# Rules of disengagement: orexin–arousal and dopamine–utility as separable axes

**DOI:** 10.64898/2026.10.25.761642

**Authors:** Nikola Grujic, Victoria Ploerer, Mattia Hamann, Alexander Tesmer, Eva Bracey, Daria Peleg-Raibstein, Denis Burdakov

**Affiliations:** Institute for Neuroscience, Department of Health Sciences and Technology, ETH Zurich, Switzerland; Chinese Institute for Brain Research, Beijing (CIBR), Beijing, China

**Keywords:** task engagement, arousal, orexin, dopamine, GLP-1

## Abstract

Staying on task, leaving it, and coming back to it are processes we experience daily. They depend on task utility and arousal, but the neurochemical processes behind this interplay are unknown. In mice performing a decision-making task with shifting reward size, we identified latent engagement states and the transitions between them. Pupil-linked arousal and dopaminergic utility coding predicted these transitions along separable axes. Orexin antagonism with almorexant disengaged mice at low reward, an effect mediated by pupil-linked arousal and not by dopamine. GLP-1 receptor agonism with semaglutide reduced the gain with which accumbal dopamine integrates reward history, which predicted the progressive disengagement it caused, with no detectable contribution from arousal or orexin. Fast striatal and slow accumbal dopamine codes predicted engagement on their own timescales. These results decouple orexin-driven arousal from dopamine-mediated utility coding in task engagement, with implications for two widely used drug classes and for disorders of motivation.

## Introduction

Everyone procrastinates. We disengage from the task we are supposed to be doing to do something else, often more immediately rewarding, and have difficulty re-engaging with it. The reason for this lack of focus is an interplay of many factors. Too little arousal (drowsiness) or too much (distractibility) pulls us away from the task and prevents us from returning to it. However, depending on how important the task at hand is, or how motivated we are for it, we can overcome the will to procrastinate and ‘lock in’.

These processes are central to everyday functioning, and their disruption is a hallmark of conditions ranging from attentiondeficit/hyperactivity disorder to apathy in depression(1, 2). Yet what determines whether we stay engaged, disengage, or return to a task remains unclear.

During motivated behavior, mice switch between different levels of task engagement(3, 4). Task engagement states and switches between them are associated with changes in pupil-linked arousal(5–8). But, despite the strong association between the two, this relationship has not been causally established by manipulating pupil-linked arousal and seeing how engagement metrics vary with it. Engagement also depends on the animal’s motivation to keep performing the task, that is, on the utility of the task, its subjective value in the animal’s current state. Here we manipulate utility through reward size. When utility is high, arousal stays close to the level optimal for engagement. When it is low, arousal drifts between low (drowsiness) and high (distractedness), toward rest or exploration of other options(5, 9), which gives rise to the inverted-U relation between arousal and performance(10) that shifts with relative arousal in humans(11). However, it is unclear which of the many systems controlling and tracking arousal(9, 12–18) are responsible for implementing this utility-dependent allocation of arousal.

The orexin system of the lateral hypothalamus encodes food cues and other stimuli, rewards, locomotion and nutritive states(19–24). It is also strong modulator of pupil-linked arousal(16), and orchestrates the activity of many other arousal systems(17). However, whether orexin integrates encoded information about nutritive state and food reward availability to recruit and allocate arousal in a taskutility-dependent manner is unknown. This arousal allocation could be mediated through their fast glutamatergic action or through orexin peptide, whose fluctuations produce slower, tonic effects on arousal(25–27) and whose levels are heightened during fasting(28). Both might be the answer, as orexin peptide antagonism has been found to decrease pupil-linked arousal in anesthetized animals, but it didn’t eliminate the orexin neuron stimulation evoked dilation(16). The orexin system is therefore a good target for investigating how utility and arousal interact, and its antagonists are increasingly prescribed for insomnia(29).

In turn, task utility is signaled by the midbrain dopamine system. Midbrain projections release dopamine in the nucleus accumbens (NAcc) and dorsolateral striatum (DLS) to encode reward value, prediction errors and movement. Dopamine in the NAcc integrates reward history(30), likely by tonic accumulation(31), to guide motivation(32) and, in foraging, the decision to leave a patch, with ventral striatal dopamine carrying the prediction errors that build the underlying value expectations(33). Dopamine in the DLS instead preferentially encodes reward and sensory prediction errors to guide decisions(34), consistent with a role in habit or skill, as well as movement and vigor(35, 36), though not consistently(37, 38). The time window over which dopamine reflects past rewards also lengthens from the dorsolateral to the ventral striatum(39). Dopamine is therefore positioned to signal the utility of the task, which determines whether it is worth continuing to perform it, as well as the vigor with which it is performed. Orexin peptide excitatory effects on the dopamine system could further change the value encoding of the environment that is part of the arousal allocation and task engagement(26, 40–42). However, whether these dopaminergic signals track task utility and predict disengagement and re-engagement, and whether this is separate from arousal allocation and potentially the orexin system, is unknown. Reinforcers in rodent experiments are usually food rewards, so utility depends strongly on the nutritive state of the animal. Given their effects on reducing appetite(43), GLP-1 agonists are a good way to lower the drive to perform the task for food. In the brain, GLP-1 agonism acts on the dopamine system through GLP-1 receptors in the amygdala, ventral tegmental area(44) (although some studies suggest GLP-1 agonism increases dopamine(45)). These effects are not constrained to food rewards, but extend to other rewarding stimuli such as drugs and wheel-running(46–48), suggesting a more general effect on dopaminergic value signaling, possibly also extending to the DLS, but this has not been tested in operant/task settings. GLP-1 agonism could additionally affect arousal directly, via GLP-1 receptors in the locus coeruleus(49), or via the orexin system described above(50). As such, GLP-1 agonism is a good manipulation to investigate how arousal and utility interact to shape task engagement. It is also highly clinically relevant, as GLP-1 receptor agonists such as semaglutide are now used by millions for weight loss and type 2 diabetes, and are being tested for substance use disorders(47, 48, 51, 52).

Here, we concurrently recorded a range of signals – pupil-linked arousal, dopamine in the nucleus accumbens (NAcc) and dorsolateral striatum (DLS), and orexin in the ventral tegmental area (VTA) and substantia nigra pars compacta (SNc), as animals performed a decision-making task with reward size switching in long trial blocks. We decomposed trials onto latent engaged and disengaged states using a hidden Markov model. We then applied orexin antagonism, orexin neuron optostimulation and GLP-1 agonism, and asked how the resulting changes in engagement relate to the measured signals.

Our recordings showed how arousal, orexin and dopamine track the amount of reward on offer and engagement, and how they predict transitions between states along separable axes. Our pharmacological and optogenetic interventions showed that these axes can be traversed separately. Orexin antagonism and orexin neuron stimulation disengaged mice mainly at low reward by disrupting the allocation of pupil-linked arousal, and antagonism also removed a protective action of orexin on engagement at low arousal. Orexin antagonism did so without changing dopamine signals, whereas GLP-1 agonism lowered engagement by reducing the gain of accumbal reward integration, with little or no effect via arousal. Together, these results define distinct routes to disengagement, set by the animal’s position on the arousal and utility axes in interaction with the current reward context.

## Results

### Reward-size-dependent modulation of task engagement

We used a standardized visual decision-making task(53) in which head-fixed mice are trained to report the side of the visual stimulus with varying contrasts appearing on the screen in front of them (left or right) by moving it to the center of the screen using a stimulus-coupled wheel under their forepaws (Fig. 1a). Correct responses were awarded with a drop of 10% sucrose solution, while incorrect responses yielded a longer inter-trial interval and a white noise (Fig. 1b, bottom). Reaction time was defined as the time between stimulus onset and outcome of the trial. To manipulate task utility, we varied the reward size in blocks of 60 trials, between 1 µL and 5 µL, always starting with low reward (Fig. 1b, top). As expected, this task revealed that animals change their reaction times with reward size, acting faster when reward was high and slower when it was low (Fig. 1c).

**Fig. 1.**
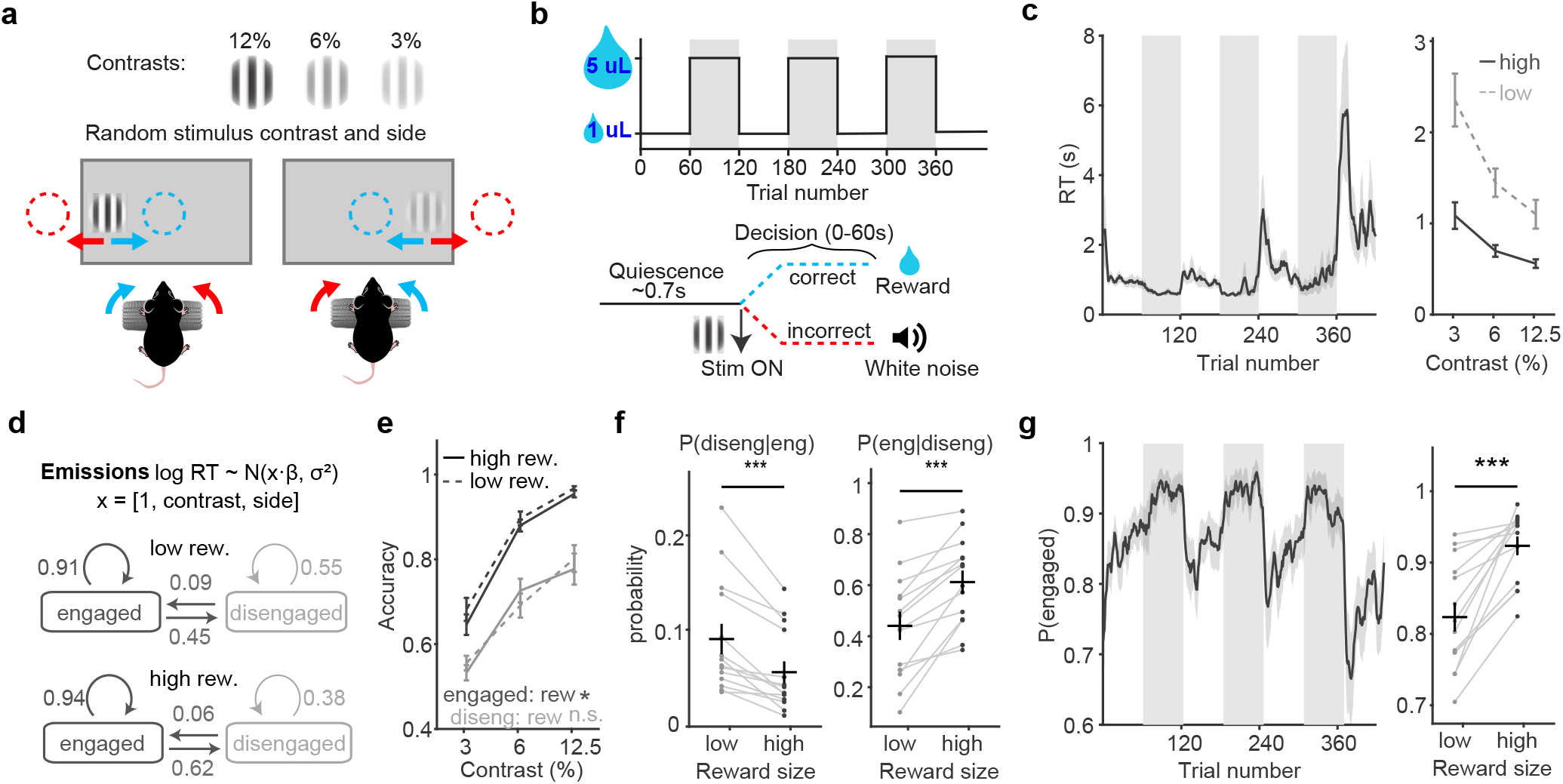
Task engagement depends on reward size. **a**, Task. Head-fixed mice turn a wheel to bring a grating (12.5, 6 or 3% contrast, left or right) to the center of the screen (blue, correct; red, incorrect). **b**, Top, reward size alternates between 1 and 5 µL in 60-trial blocks, starting with low reward (gray, high-reward blocks). Bottom, trial timeline. **c**, Reaction time across the session (left) and against contrast in high (solid) and low (dashed) reward blocks (right). **d**, The engagement model (RT-HMM) and the mean transition probabilities between the engaged and disengaged state in low and high reward blocks. **e**, Accuracy against contrast in the engaged (dark) and disengaged (gray) state, high (solid) and low (dashed) reward; text, low vs high reward within each state (paired t-tests on the mean over contrasts; disengaged state, 12 mice with enough trials). **f**, Disengage hazard, P(disengage | engaged), and re-engage hazard, P(re-engage | disengaged), per mouse in low and high reward blocks; stars, low vs high reward (paired t-tests). **g**, P(engaged) across the session (left) and per mouse (right; stars, low vs high reward, paired t-test). Vehicle sessions of both experiments, pooled within mouse; n = 14 mice. Statistics in Supplementary Table 1.

In order to better quantify task engagement we applied a twostate hidden Markov model(6) – allowing us to split trials into two states – engaged and disengaged and examined the probabilities of switching between them (the disengage and re-engage hazards, Methods) in the two reward blocks (Fig. 1d; Extended Data Fig. 1a–e). These extracted states were tracked by additional measures that were not included in the model like accuracy (Fig. 1e) which revealed a large difference between the two states and a small difference between low and high reward blocks in engaged states. Additionally, pre-reward movement and post-reward licking were reduced in disengaged trials (Extended Data Fig. 1f,g). These results are consistent with these being separate behavioral states. The extracted probabilities of disengaging and re-engaging were strongly modulated by the reward block, leading to a change in engagement probability which fell in low-reward and rose in high-reward blocks (Fig. 1f,g).

### Arousal, orexin and dopamine track reward size and predict engagement transitions along separable axes

Having verified that our paradigm revealed engagement states that depended on our manipulation of reward size, we looked at how our recordings of pupil size, dopamine (dLight) and orexin (oxLight) tracked both reward size and engagement.

Baseline pupil size (residualized and z-scored to preserve absolute differences, see Methods) replicated previous patterns, tracking engagement states, reward blocks and transitions within them(5, 6) (Fig. 2a). The relationship between pupil and engagement was weaker in high-reward blocks, making state occupancy and transitions more sensitive to arousal under low reward (Fig. 2a). Orexin release in SNc and VTA (oxLight) also showed dynamic modulation by reward block, rising through low-reward blocks and falling through high-reward blocks (Extended Data Fig. 2a,c). In SNc this dynamic was implemented through running integration of previous rewards with a time constant of around 10 trials, rather than through the block itself. VTA orexin followed mainly the last reward (Extended Data Fig. 2a,g).

**Fig. 2.**
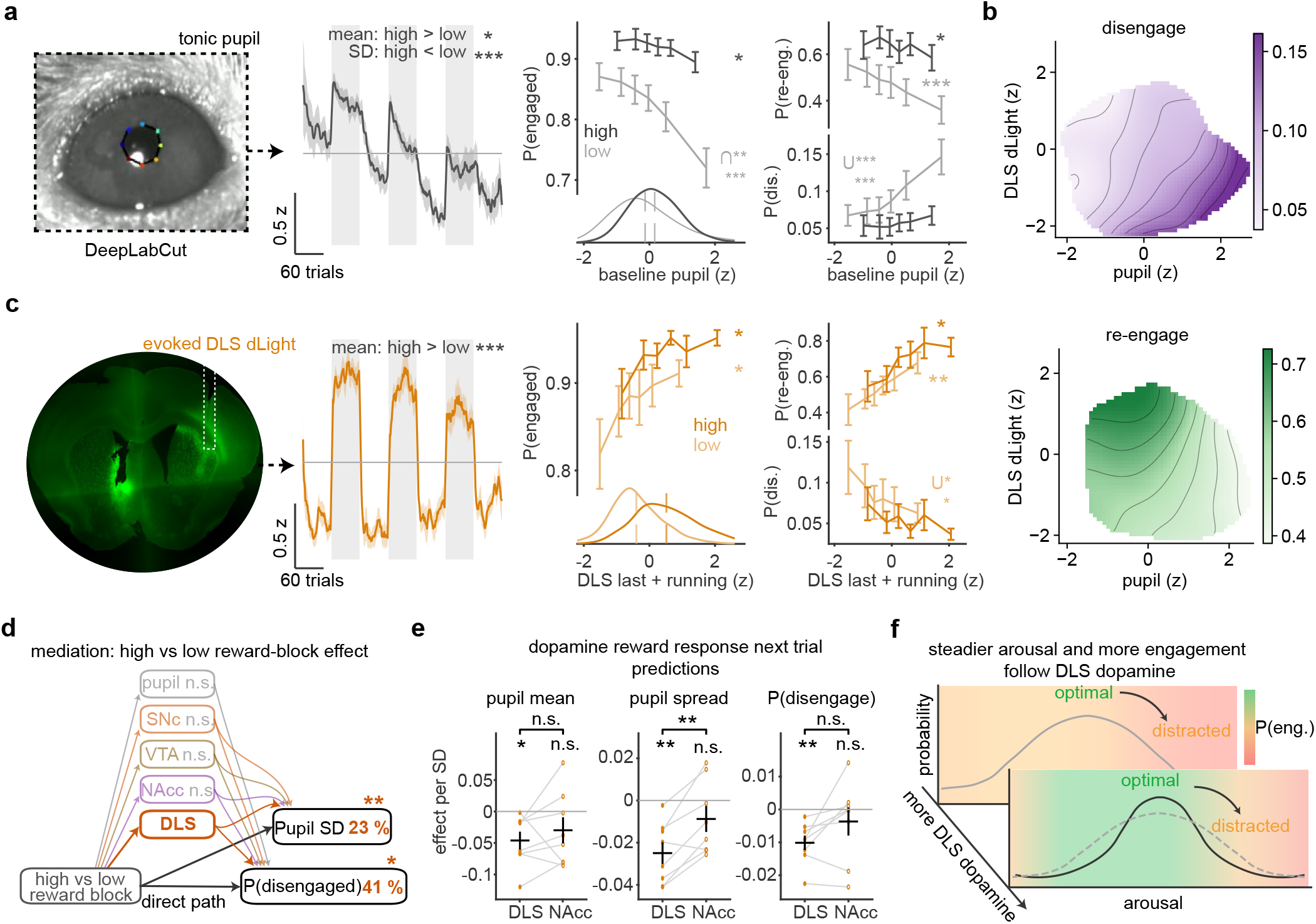
Arousal and utility are separable axes of engagement, with reward size shaping the arousal distribution. **a**, Pupil. An example frame with the tracked pupil, the baseline pupil across the session (gray, high-reward blocks), and P(engaged), P(disengage | engaged) and P(re-engage | disengaged) against the baseline pupil in high (dark) and low (light) reward blocks (six quantile bins per block), with each block’s distribution below P(engaged) (ticks, means; text, high − low difference in mean and within-block SD, paired t-tests). Hazards are the transition into trial n from the pupil of trial n (Methods). **b**, Disengage (top) and re-engage (bottom) probability in low-reward blocks over pupil and the DLS measure of c, smoothed per mouse and averaged over mice. **c**, The same as a for the DLS dLight reward response (left, recording site), as the reward response of trial n − 1 plus the running average of earlier responses (‘last + running’, Methods). **d**, Share of the high − low reward-block difference in within-block pupil SD and in P(disengaged) that each signal accounts for, on trials after a rewarded trial (quantile replacement of each signal’s high-block values by its low-block distribution, Methods); stars, one-sample t-tests across mice. The shares fitted on all trials, with wheel movement and licking as additional covariates, are in Extended Data Fig. 2f. **e**, Fast coupling. Effect of the previous trial’s reward response (per SD) on the next trial’s baseline pupil, its deviation from the block mean and the probability of disengaging, for DLS and NAcc, from per-session linear models (Methods); dots, mice (n = 8 with both signals); stars, against 0; bracket, DLS vs NAcc (paired t-test). **f**, Schematic. Engagement is most likely in an intermediate arousal range; after larger DLS reward responses the arousal distribution is steadier and engagement more likely. Vehicle sessions of both experiments, pooled within mouse; n = 14 mice (pupil), 9 (photometry; SNc 8). Statistics in Supplementary Table 1.

Expectedly, NAcc and DLS dopamine reward responses (dLight) strongly tracked the reward block, as previously described(34, 54–57) (Fig. 2c; NAcc, Extended Data Fig. 2d). In NAcc the tonic baseline tracked the block as well, implemented, as for SNc orexin, through integration of recent reward. DLS showed no such tonic tracking (Extended Data Fig. 2g).

The five measures were only weakly correlated trial by trial (|r| ≤ 0.16 except between the same sensors in different sites; Extended Data Fig. 2e). Even the correlation of orexin release with pupil size seen in free-running head-fixed settings(16, 20) was absent in both regions. Each signal’s contribution to the disengagement and re-engagement probabilities of the trial ahead could therefore be examined separately. Pupil size was the most predictive of both, with a larger baseline pupil predicting more disengagement and less re-engagement (Fig. 2a,b). Dopamine responses on the previous trial predicted the opposite, less disengagement and more re-engagement (Fig. 2b,c; NAcc, Extended Data Fig. 2d). Higher orexin predicted more disengagement and less re-engagement, depending on the recorded location (Extended Data Fig. 2a–c). The effects of the five signals on the hazards were largely additive. In a joint model with all five signals and their ten pairwise products, only two sub-additive interactions survived correction, both on disengaging in high-reward blocks, where the pupil coupling is shallow anyway (Extended Data Fig. 2h).

Because DLS reward responses are larger in high than in low-reward blocks, we asked how much of the block difference in engagement and arousal it, and other signals, account for. When each signal’s values in high-reward blocks were replaced by values from its low-block distribution, so that the signal no longer differed between blocks while everything else stayed, only the DLS response carried a significant part of the block effect, that is, of the difference in engagement between high- and low-reward blocks. On trials after a rewarded trial it accounted for about 40% of the effect on P(disengaged) (somewhat less when including all trials) and about a quarter of the effect on the within-block variability of the pupil, whereas pupil, SNc, VTA and NAcc carried no significant share (Fig. 2d; Extended Data Fig. 2f). From trial to trial, a larger DLS reward response predicted a smaller baseline pupil, a smaller deviation of the pupil from its block mean and a lower probability of disengaging on the next trial. The NAcc response, measured the same way, did not significantly predict any of the three, and DLS predicted the pupil’s deviation more strongly than NAcc (Fig. 2e). These DLS relations held within blocks and when the engagement states of the two trials were taken into account (Methods), and the smaller spread came mainly from fewer trials with a large baseline pupil. DLS dopamine thus tracks the current value of the task and predicts steadier arousal and continued engagement on the next trial (Fig. 2f).

These results set the arousal and utility axes along which the probabilities of disengagement or re-engagement move, with orexin release as a further predictor. The axes are separable, additive in the hazards, while dopamine and the reward context shape where on the arousal axis the animal sits (Fig. 2d–f).

### Reward-size-dependent modulation of engagement states by orexin antagonism and GLP-1 agonism

Next we looked at how our pharmacological manipulations affected task engagement in relation to the reward context.

Animals were injected with the dual orexin antagonist almorexant (30 mg/kg) or the GLP-1 agonist semaglutide (0.03 mg/kg), with the respective vehicles as control.

Almorexant caused disengagement specific to low-reward blocks (Fig. 3a–c, top), whereas semaglutide caused progressive disengagement independent of reward size (Fig. 3a– c, bottom). The hazard functions showed the same split. Almorexant mainly affected engagement via disengagement, and only in low-reward blocks, with a trend in re-engagement (Fig. 3e). Semaglutide both increased disengagement and reduced re-engagement in high and low reward blocks (Fig. 3f). Since engaged trials are the majority, a given change in the disengage probability affects the overall state structure more than the same change in the re-engage probability. The two engagement states showed a large difference in accuracy, independent of reward size, which neither drug affected in either state (Fig. 3d; Extended Data Fig. 3c), suggesting that the drug effects on performance reflect a change in state occupancy and not in perception.

**Fig. 3.**
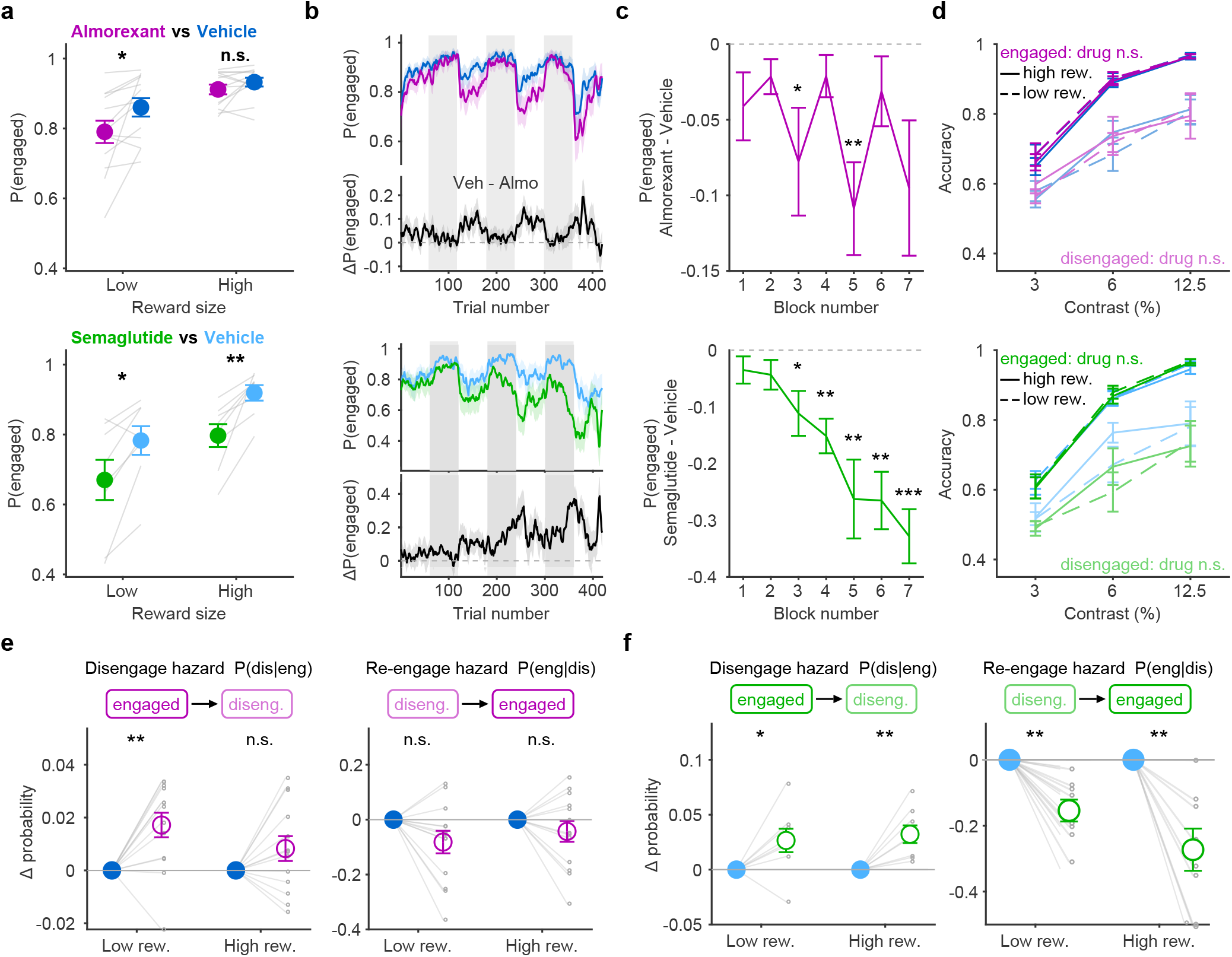
Almorexant disengages mice at low reward only, semaglutide at both reward sizes. **a**, P(engaged) at each reward size, almorexant vs vehicle (top) and semaglutide vs vehicle (bottom). **b**, P(engaged) across the session (upper; gray, high-reward blocks) and the control − drug difference (lower). **c**, Drug − control P(engaged) per reward block (odd blocks low reward; one-sample t-tests against 0). **d**, Accuracy against contrast in the engaged (dark) and disengaged (light) state, high (solid) and low (dashed) reward, drug vs control; text, drug effect within each state (paired t-tests on the mean over contrasts; contrast cells with at least 5 responded trials, so every mouse enters). **e, f**, Drug − control change in the disengage and re-engage hazards in low and high reward blocks for almorexant (**e**) and semaglutide (**f**); filled circle, control; open circle, drug. n = 13 mice (almorexant/vehicle) and 8 mice (semaglutide/vehicle). Statistics in Supplementary Table 1.

The same pattern was present in the raw reaction times, before any state assignment (Extended Data Fig. 3b,d), and in the reward consumed per day, which fell by only 3% under almorexant and by 13–14% under semaglutide, so the drug effects were not a consequence of large differences in reward consumed (Extended Data Fig. 3a).

To see whether the reward-size switching is necessary to detect these effects, we ran a control experiment with a constant reward (2 µL). Almorexant no longer lowered session engagement (even slightly raising it), whereas semaglutide produced the same progressive disengagement (Extended Data Fig. 3e–h). We also checked movement (wheel turning) and licking. The only drug effect on pre-reward movement was a slight reduction in engaged trials under almorexant (Extended Data Fig. 1f), consistent with the reduced response vigor reported in orexin deficiency and under almorexant(21, 58, 59). Licking was reduced only under semaglutide, in high-reward trials that were already disengaged, and was preserved in engaged trials (Extended Data Fig. 1g).

Orexin peptide action is thus needed to sustain engagement when reward is low, whereas GLP-1 receptor agonism causes an overall disengagement independent of reward size, consistent with a reduction in appetitive drive.

### Orexin antagonism lowers engagement on the arousal axis, but not the utility axis

Having identified the axes of engagement, we next examined how the effects of orexin peptide signaling on engagement are mediated along them. Unexpectedly, almorexant increased baseline pupil size (Fig. 4a), in both reward blocks, both engagement states and throughout the session (Extended Data Fig. 4a,b), opposite to its effect under anesthesia(16). The increase was not an effect on the eye itself. Almorexant did not change the pupillary light reflex or pupil size under task lighting outside the task, and it reduced pupil size in darkness (Extended Data Fig. 4i). Within the task it steepened the pupil–movement relation, enlarged reward-evoked dilations despite the higher baseline, and on trials without wheel movement raised the pupil on engaged trials while lowering it on disengaged ones (Extended Data Fig. 4c–h). Together these point to a central change in arousal regulation during the task, not to a peripheral effect on the pupil.

**Fig. 4.**
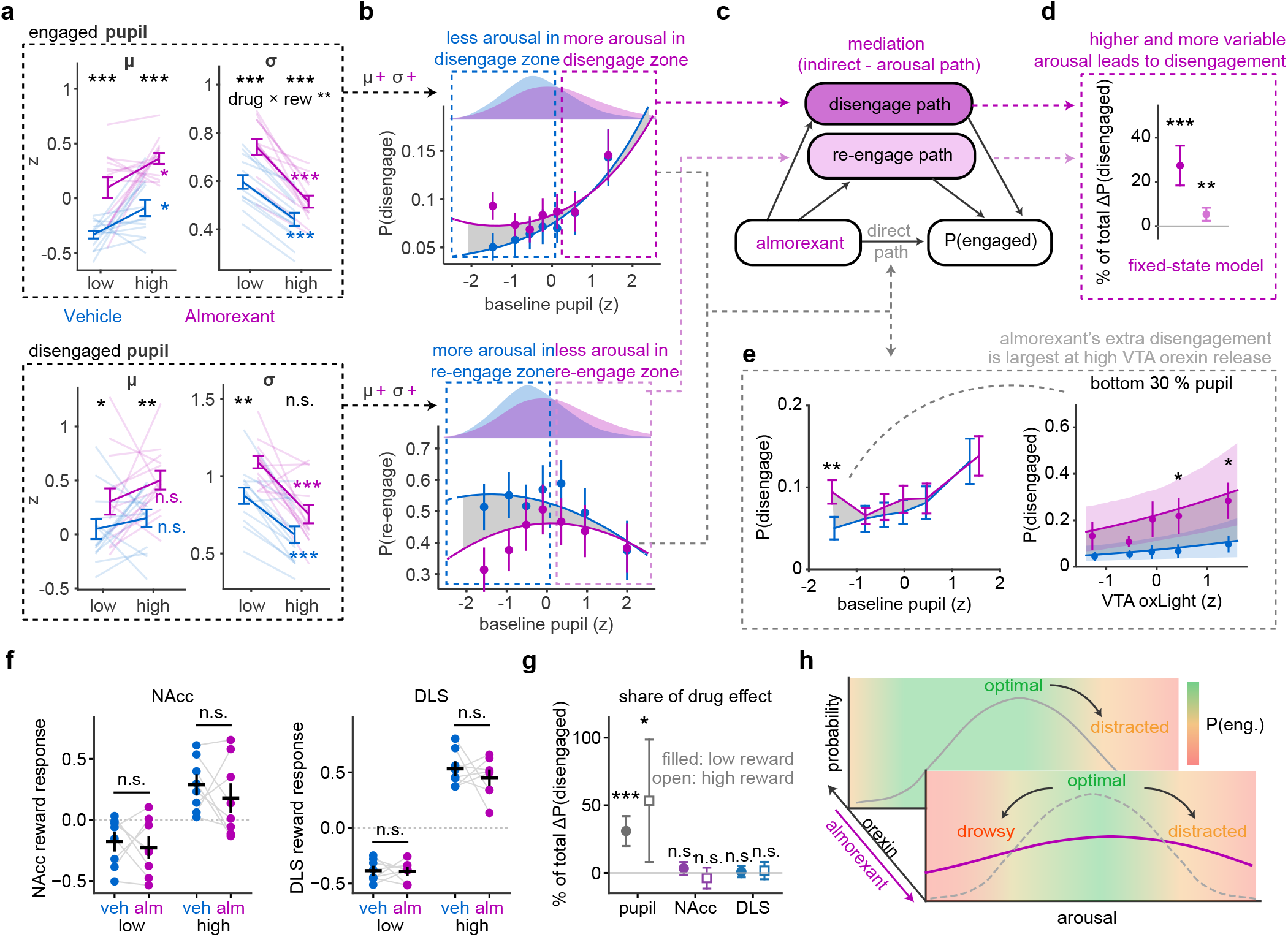
Arousal, not dopamine, mediates the reward-context-dependent disengagement under orexin antagonism. **a**, Mean (*µ*) and within-block SD (*σ*) of the baseline pupil (within-mouse z) in low and high reward blocks, on engaged (top) and disengaged (bottom) trials, vehicle vs almorexant; black stars, drug vs vehicle; colored stars, low vs high reward; text, drug *×* reward interaction where significant. **b**, Disengage (top) and re-engage (bottom) hazards against the baseline pupil in low-reward blocks. Points, binned data; lines, the fitted hazard model (Methods); gray, the difference; above, the pupil distributions of vehicle and almorexant trials; boxes, pupil ranges in which almorexant had fewer (left) or more (right) trials than vehicle. **c**, The mediation. Almorexant acts on engagement through arousal, via the disengage and re-engage hazards, and directly. **d**, Share of almorexant’s effect on P(disengaged) carried by arousal through each hazard in low-reward blocks (almorexant’s pupil distribution under the vehicle hazards, the pure indirect effect, Methods); mean and 95% CI across mice, one-sample t-tests. **e**, Left, P(disengage) against the baseline pupil in low-reward blocks (six quantile bins; stars, per-bin paired t-tests). Right, P(disengaged) against the VTA oxLight baseline within each mouse’s bottom 30% of baseline pupil, low reward; points, observed; lines and bands, hazard model with oxLight *×* drug (mean and 95% cluster-bootstrap interval); stars, per-bin paired t-tests on the observed values. **f**, NAcc and DLS reward responses (last + running, z) in low and high reward blocks, vehicle vs almorexant. **g**, Share of almorexant’s effect on P(disengaged) carried by pupil, NAcc and DLS (as d, same trials; filled, low reward; open, high). **h**, Schematic. Almorexant raises and widens the arousal distribution, moving trials into ranges where disengagement is likely, and adds disengagement at low arousal when orexin release is high. n = 13 mice (a, b, d, e left), 7 (e right), 8 (f, g). Statistics in Supplementary Table 1.

A drug can lower engagement through arousal in two ways. It can move the arousal distribution into ranges where the hazards already favor disengagement, or it can change the hazards themselves, so that the same arousal carries a higher risk. Almorexant did both, but only where the task was worth little. Under both vehicle and almorexant, pupil baselines fluctuated less in high-than in low-reward blocks, and almorexant widened them mainly in low-reward blocks (Fig. 4a). In low-reward blocks, where the hazards depend steeply on pupil size, this higher and more variable pupil placed more trials in the range where control mice disengage (Fig. 4b; Extended Data Fig. 5b, left). In high-reward blocks almorexant raised mean pupil size as well but widened the distribution far less (drug × reward interaction on *σ*, engaged trials, Fig. 4a). There the hazards depend only weakly on pupil size and the hazard curve was unchanged, so the shift changed the hazard little and engagement was spared (Fig. 3a,e; Extended Data Fig. 5b, right). To quantify this route, we passed almorexant’s pupil distribution through the vehicle hazard curves and recomputed the time spent disengaged (a pure indirect effect, Methods). Arousal accounted for about a third of almorexant’s effect in low-reward blocks, almost entirely through the disengage hazard (Fig. 4c,d), and a generative RT-HMM in which pupil size drives the transitions directly gave the same proportions (Extended Data Fig. 5a). The reward-size specificity of this route thus has two sources, the arousal–hazard coupling that is steep only when reward is low, and the loosening of arousal control mainly when reward is low, which suggests that orexin signaling is part of how high reward keeps arousal stable.

The second route appeared where arousal was low. At low pupil sizes mice disengaged more under almorexant than under vehicle at the same arousal (Fig. 4e, left). On these low-arousal trials (bottom 30% of pupil size), vehicle mice stayed engaged regardless of how much orexin was being released in the VTA, whereas under almorexant engagement fell the more orexin was released (Fig. 4e, right; Extended Data Fig. 6a). On high-arousal trials the dependence on VTA release was weaker, with no bin differing between drugs (Extended Data Fig. 6a,b), and SNc release showed no drug difference at either arousal level (Extended Data Fig. 6b). The cost of blocking orexin therefore scaled with the amount of orexin signaling being blocked, most clearly when arousal was low. Since orexin release rises as reward becomes scarce (Extended Data Fig. 2a,g), this suggests that orexin is recruited to hold mice on task when arousal sags in low-reward blocks, which is why high release predicts disengagement in control sessions even though removing it causes disengagement. Two controls support reading the VTA oxLight signal this way. The drug’s effect depended on the measured release level rather than being uniform, so the sensor was tracking behaviorally relevant dynamics, and almorexant itself did not detectably alter the oxLight signal. Reward responses, the reward-block code and the weight of the baseline in predicting the transitions were all unchanged (Extended Data Fig. 6c–e). Consistent with the two routes, almorexant’s additional disengaged trials were of two kinds, slow responses on trials with a large pupil and lapses (no response, or a response after more than 10 s) at low pupil that accumulated mid-session (Extended Data Fig. 6f–h).

Almorexant did not act through dopamine. Reward responses in NAcc and DLS, the dopamine measures that predict the hazards (Fig. 2), were unchanged under almorexant in both reward blocks (Fig. 4f; event-locked responses, Extended Data Fig. 6i), and substituting the almorexant dopamine distributions into the vehicle hazards carried none of the drug’s effect (Fig. 4g). This differs from reports of orexin effects on dopamine signaling in other settings(26, 40–42, 60–63).

Almorexant thus lowers engagement through two routes on the orexin–arousal axis, neither of which involves dopamine (Fig. 4h). If the first route is causal, raising arousal by other means should also disengage mice in low-reward blocks, which we tested next by stimulating orexin neurons.

### Orexin neuron stimulation lowers engagement along the arousal axis

To increase orexin neuron output (and pupil-linked arousal), we stimulated orexin neurons during the reward-size switching task in a repeating pattern of 30 s laser on and 1 min laser off (Fig. 5a). As expected, stimulation produced frequency-graded pupil dilation. This dilation was absent in heat/light control animals and in sham control trials from unstimulated sessions (Fig. 5b,c). Sham trials were important because selecting trials that started during laser stimulation cuts out longer RT trials, which are more likely to be disengaged, and so can bias the selection. Interestingly, the dilation caused by orexin neuron stimulation was itself modulated by the reward context and was larger in low-reward blocks (Fig. 5e, Extended Data Fig. 7c,d). The arousal evoked by orexin neuron stimulation therefore depends on the reward context, consistent with a utility-dependent recruitment of arousal.

**Fig. 5.**
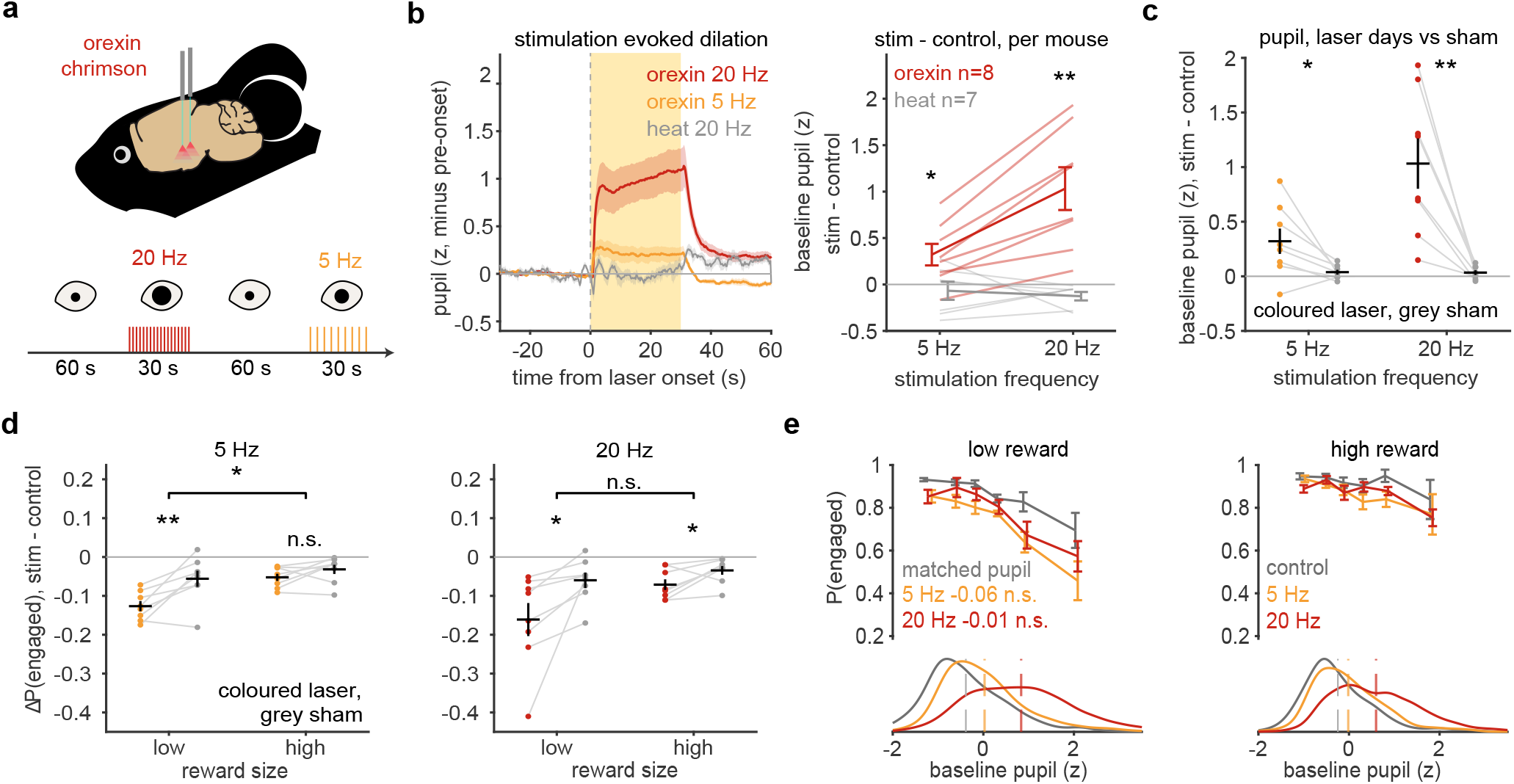
Orexin neuron stimulation disengages mice, most at low reward, in step with the arousal it evokes. **a**, ChrimsonR in lateral-hypothalamic orexin neurons; 30-s bouts at 20 or 5 Hz separated by 60 s without laser. **b**, Left, continuous pupil aligned to laser onset (z, minus the pre-onset level; yellow, laser on). Right, stimulated − control trial baseline per mouse at 5 and 20 Hz, orexin vs heat controls. **c**, The same change on laser days (colored) and for sham bouts laid over no-laser days and labeled by the same rule (gray). **d**, Stimulated − control P(engaged) at 5 Hz (left) and 20 Hz (right), laser (colored) vs sham (gray), in low and high reward; stars, laser vs sham; bracket, low vs high. **e**, P(engaged) against baseline pupil for control, 5-Hz and 20-Hz trials on laser days in low (left) and high (right) reward; text, laser − sham shift at matched pupil (low reward; 7 mice with enough trials per bin); below, pupil distributions (ticks, means). n = 8 orexin and 7 heat mice. Statistics in Supplementary Table 1.

Stimulated trials showed disengagement at both reward sizes. It was larger in low-reward blocks at 5 Hz, with the same tendency at 20 Hz (Fig. 5d). In the engagement versus baseline pupil curves, the large increase in baseline pupil under 20 Hz stimulation pushed the animal into an arousal range in which disengagement is likely, in low reward. The smaller dilation in high reward, together with the lower transition rates there, produced a smaller drop in engagement (Fig. 5e). Engagement did not change in heat/light control mice (Extended Data Fig. 7a).

At 20 Hz, disengagement and re-engagement at matched pupil did not differ from non-stimulated trials in the same mice, consistent with an effect carried by arousal (Extended Data Fig. 7b). In a mediation analysis, the larger pupil under 20 Hz stimulation accounted for a third to a half of the extra disengagement through the disengage hazard alone, with a smaller contribution through slower re-engagement (Extended Data Fig. 7e). At 5 Hz, baseline pupil changed little, yet disengagement still rose (Fig. 5e, Extended Data Fig. 7b). Mediation therefore assigned only a small share of the 5 Hz effect to arousal.

Because stimulation sessions were interleaved with sham sessions, we could compare sham trials with no-laser trials from laser sessions in the same mice. In these post-stimulation trials the fast activation of orexin neurons had ended, while its slower consequences may still have been present. We matched sham and post-laser trials in pupil size to exclude the effect of still elevated pupil-linked arousal. Here engagement increased in low-reward trials, and the increase scaled with pupil size (Extended Data Fig. 7f). The effect persisted, less robustly, when conditioned on an engaged previous trial (Extended Data Fig. 7f). Engagement therefore moved in opposite directions during stimulation and in the period after it.

Orexin neurons are a small subset of the glutamatergic (vGlut2) neurons of the lateral hypothalamus, which as a population also promote arousal(64, 65). To ask whether a broader arousal drive from the same region shows the same reward-size dependence, we stimulated this wider population at the same location. The stimulation caused strong dilation (Extended Data Fig. 7h,i), paired with strong disengagement that scaled only with stimulation frequency and was independent of reward size (Extended Data Fig. 7g). A strong enough increase in arousal therefore disengaged mice even at high reward.

Together, these experiments show that orexin neuron stimulation increases disengagement mainly through arousal, and that the size of this effect depends on reward size. They also implicate wider LH glutamatergic populations in arousal control, since their stimulation disrupted engagement even in high-reward trials.

### GLP-1 agonism lowers engagement along the utility axis

Next, we looked at which signals semaglutide changed and how they mediated its effect on engagement. Disruptions of dopaminergic signaling in the NAcc are well known to impair motivation to work for reward(30, 32, 66), and the tonic level has been suggested to carry the motivation signal. At reward block switches, the tonic NAcc level rose progressively when switching from low to high reward, and fell when switching from high to low (Fig. 6a). Semaglutide disrupted this updating (Fig. 6a, right), and with it the tonic coding of utility (Fig. 6c; Extended Data Fig. 9c). Less of the reward history was integrated into the tonic level, at block changes and within a block (Extended Data Fig. 8a–d). The tonic level integrated the reward obtained rather than confidence-weighted prediction errors (Extended Data Fig. 8e).

**Fig. 6.**
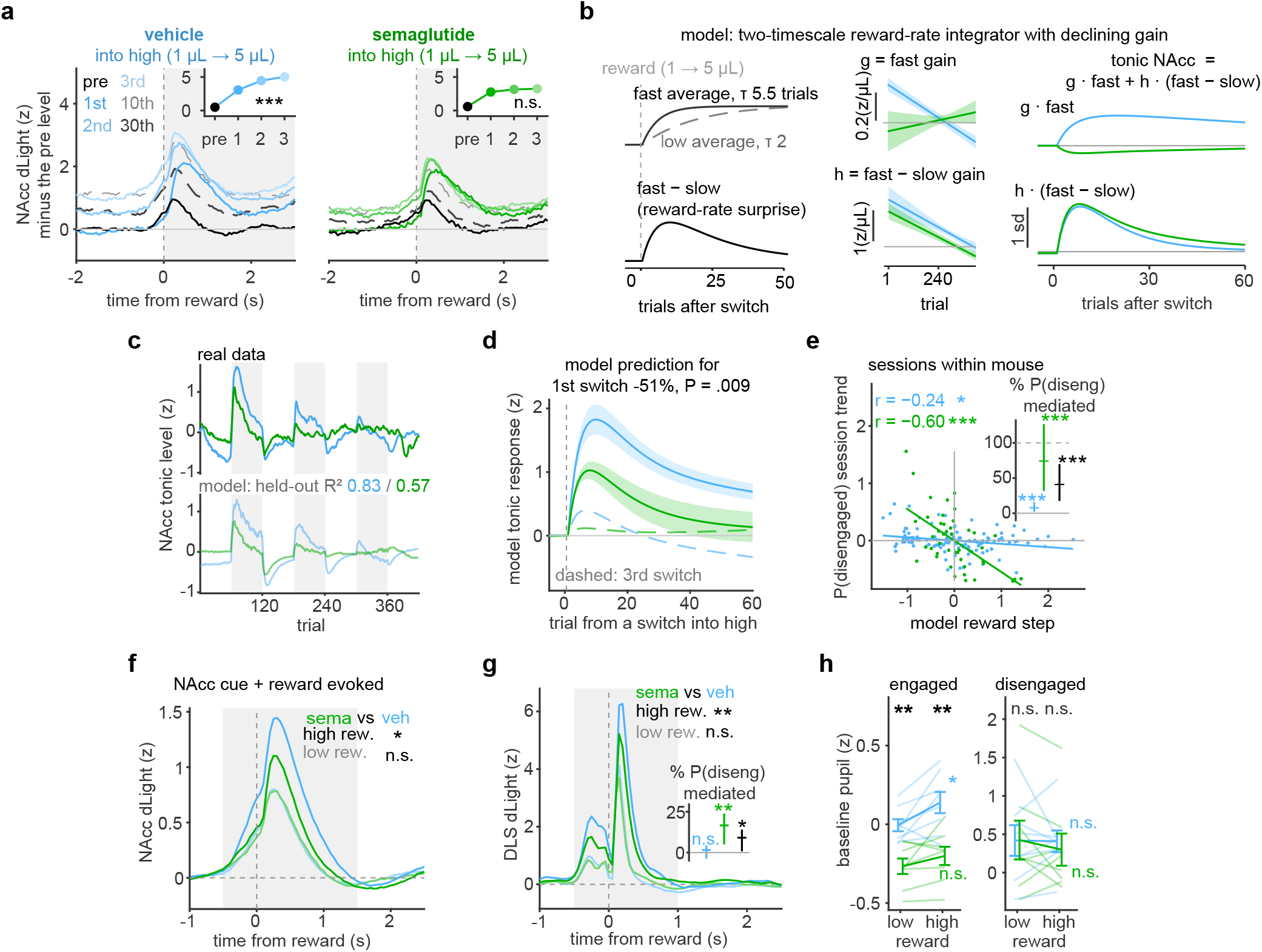
The gain of accumbal reward integration, not arousal, mediates disengagement under semaglutide. **a**, NAcc dLight at switches into high reward (1 *→* 5 µL), vehicle (left) and semaglutide (right); reward-aligned traces of the last pre-switch trial and of early and late trials of the new block, minus the pre-switch tonic level. Insets, area 0–3 s after reward; star, rise from the 1st trial to the 2nd and 3rd. **b**, The two-timescale reward-rate integrator (Methods). Left, fast and slow reward averages after a step and their difference. Middle, fitted gains g and h across the session. Right, the two terms after a switch into high reward. **c**, Tonic NAcc level by trial, averaged over sessions (top), and the integrator’s held-out prediction (bottom); R^2^, vehicle / semaglutide. **d**, Integrator response to a step up in reward at the 1st (solid) and 3rd (dashed) switch into high reward; above, semaglutide − vehicle at the 1st switch. **e**, Per session, the integrator’s step at the first switch against the session trend of P(disengaged) (Methods), centered within mouse and condition; vehicle sessions of both experiments (9 mice) and semaglutide (8 mice); r, within-mouse correlation (permutation P). Inset, share of semaglutide’s effect on the trend mediated by the step, with the step–trend relation from vehicle sessions, from semaglutide sessions, and their mean; 95% cluster-bootstrap intervals. **f**, NAcc dLight aligned to reward in high (dark) and low (light) reward blocks, vehicle vs semaglutide; gray, AUC window; text, semaglutide vs vehicle on the AUC. **g**, The same for DLS (per mouse, Extended Data Fig. 10b). Inset, share of semaglutide’s increase in P(disengaged) mediated by the trial-level DLS response, same three estimates; the two insets have different outcomes and do not add. **h**, Mean baseline pupil (within-mouse z) in low and high reward blocks, engaged and disengaged trials, vehicle vs semaglutide; black stars, drug vs vehicle; colored, low vs high reward. n = 8 mice per arm. Statistics in Supplementary Table 1.

We modeled the tonic level as a two-timescale reward-rate integrator (Fig. 6b). A fast average tracked the reward of the last ∼5 trials and a slow one the reward over the last 20– 30 trials. The tonic level followed the fast average plus the recent reward minus the slow average. The second term produced the overshoot after a switch and its decay within the block. Each term had a gain that changed with the amount of reward consumed. Both gains declined as reward accumulated, like a decreasing motivational drive, and semaglutide lowered them further and almost eliminated the fast gain (Fig. 6b). Fitted per mouse and predicted on sessions not used for the fit, the model reproduced the session-averaged tonic level (Fig. 6c). Semaglutide strongly reduced the model’s response to a reward step (Fig. 6d), whereas almorexant did not (Extended Data Fig. 8f). This explained the tonic compression of the utility code, while the phasic responses to the first rewards of a new block were unchanged (Extended Data Fig. 8g,h).

The suppression of phasic NAcc utility signaling (Extended Data Fig. 9b,c) and of the model’s response to a reward step (Fig. 6d) was already present in the first two blocks, before engagement had changed significantly under semaglutide (Fig. 3c, bottom). We therefore asked whether these early measures predicted the later rise in disengagement. Per session, a smaller integrator step at the first switch into high reward predicted a steeper rise of P(disengaged) over the session, within semaglutide sessions and within the vehicle sessions of both experiments (Fig. 6e). In vehicle sessions, a larger early step also predicted a smaller decline of the phasic DLS response and a smaller rise of pupil variability over the session (Extended Data Fig. 10f). Semaglutide reduced this step. How much of the drug’s effect on the session trend that reduction explains depends on which step– trend relation is used. With the relation measured in vehicle sessions it accounted for under a tenth, with the steeper relation within semaglutide sessions about three quarters, and about two fifths on their mean (Fig. 6e, inset; Methods). Between a tenth and three quarters of the extra disengagement that builds up over a session is thus attributable to the smaller step.

Semaglutide also compressed the phasic reward responses. It reduced the high-reward response in NAcc and DLS while leaving the low-reward response unchanged (Fig. 6f,g; Extended Data Fig. 9b,c), so the contrast between high and low reward in the evoked response fell, while the baseline differences between blocks remained (Extended Data Fig. 9f,g). The reduction held when vehicle and semaglutide recordings were matched within mouse by signal brightness (Extended Data Fig. 9e), and it was absent in the constant reward control task (Extended Data Fig. 9a).

The effect of semaglutide was more confined in the DLS, where we did not observe tonic reward integration. The DLS phasic code was also compressed (Extended Data Fig. 10a–e,g). Semaglutide lowered the DLS reward response only in high-reward blocks (Fig. 6g; Extended Data Fig. 10b), whereas it raised disengagement in both, and in high blocks it shrank mainly the largest responses, in a range where the disengage hazard is already low and flat, so lowering those responses changed the hazard little. Accordingly, the trial-level DLS change carried only a small share of the extra disengagement through the vehicle relation, and under a tenth on the mean of both relations (Fig. 6g, inset). The DLS compression is therefore mainly a parallel change in reward-size signaling, a minor route at most, which points to pathway-specific action of GLP-1 agonism rather than an overall dopamine effect. Semaglutide also reduced the reward-size prediction error at block switches in the DLS, and the high-reward response was reduced within blocks as well (Extended Data Fig. 10c–e). This was not seen in the NAcc (Extended Data Fig. 8g,h). Early-session VTA and SNc oxLight codes did not predict late disengagement (Extended Data Fig. 9h).

Finally, we asked whether semaglutide also acted on the arousal axis. It lowered pupil-linked arousal overall, but only in engaged trials (Fig. 6h), a change that should protect against disengagement (Fig. 2a). Unlike almorexant, it did not change the variability of baseline pupil in engaged trials, only in disengaged trials (Extended Data Fig. 5c). An arousal route to the disengage hazard is therefore unlikely, with at most an effect on re-engagement through the variability change. Indeed, the mediation analyses found no effect (Extended Data Fig. 5a).

Semaglutide therefore lowers engagement along the utility axis. It reduced the gain of tonic reward integration in the NAcc, and this reduction predicted its progressive disengagement (Fig. 6a–e). The compression of high reward phasic signals in NAcc and DLS (Fig. 6f,g) accompanied the behavioral effect, and arousal carried little or none of it (Fig. 6h).

## Discussion

### Two separable axes of task engagement

The ability to focus on the task at hand is of great personal and societal importance, and so is understanding how the brain implements it. Many studies have related arousal and utility to engagement, but the neural systems behind this modulation(17) have not been disentangled. Here we recorded signals that integrate arousal, utility or both, and manipulated them with two classes of drugs already in widespread use (GLP-1 agonists and orexin antagonists(29)), to investigate the physiological and pharmacological principles that govern task engagement. We show that the probability of being engaged depends on two separable axes, pupil-linked arousal and the dopaminergic utility code, with orexin acting on the arousal axis and, when arousal is low, directly on engagement. We show how animals traverse this space and how the two classes of commonly used drugs change the way they do so (Fig. 7c).

**Fig. 7.**
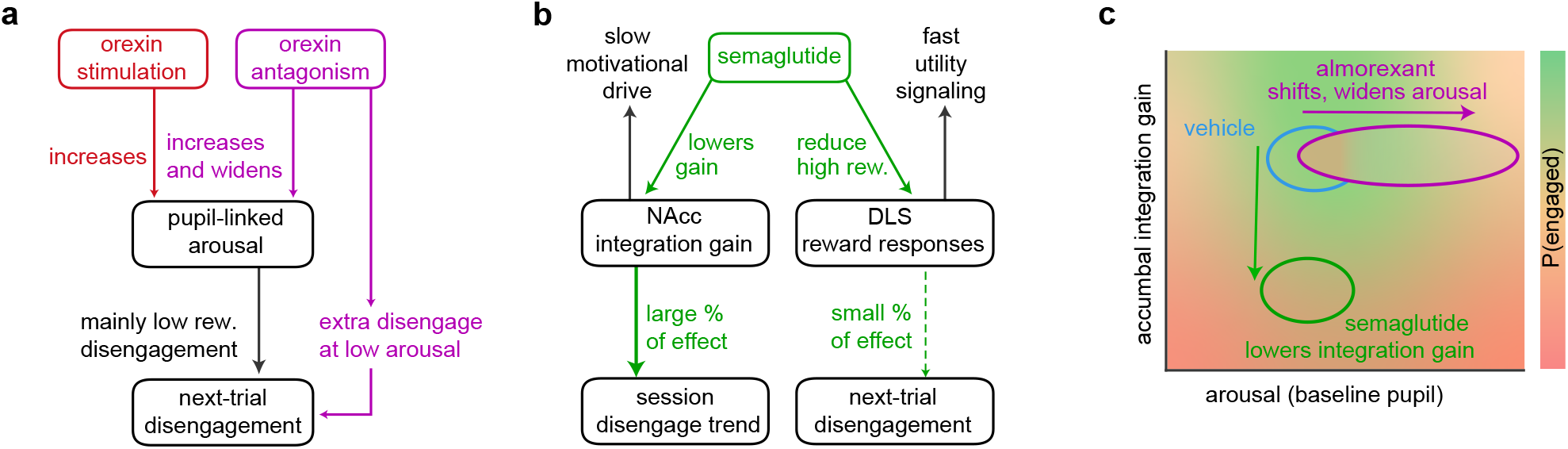
Two axes of engagement and two routes to disengagement. **a**, The arousal axis. Orexin neuron stimulation (red) and orexin antagonism (magenta) raise pupillinked arousal, and the raised arousal increases next-trial disengagement, mainly at low reward (Figs. 4 and 5). Antagonism adds a direct route, extra disengagement at low arousal where orexin release is high (Fig. 4e), and leaves the dopamine measures unchanged (Fig. 4f,g). **b**, The utility axis. Semaglutide lowers the gain of accumbal reward-rate integration, which carries between a tenth and three quarters of the rise of disengagement over a session (Fig. 6e), and compresses the DLS reward response, which carries little of the trial-level effect (Fig. 6g); arrow weight, share of the effect; the two outcomes differ and the shares do not add. Pupil-linked arousal carries little or none of the effect (Fig. 6h, Extended Data Fig. 5a,c). **c**, The manipulations on the plane of pupil-linked arousal and accumbal integration gain (schematic; color, P(engaged)). Almorexant shifts and widens the arousal distribution, semaglutide lowers the gain. Statistics in Supplementary Table 1.

Orexin, dopamine and arousal each predicted transitions between engagement states, with largely additive effects (two sub-additive interactions, both in high-reward blocks where the arousal coupling is shallow), which sets them up as separable axes. Rising pupil and orexin predicted disengagement, whereas the previous trial’s dopamine response predicted engagement. These signals predicted not only the current state but the next transition, disengagement or re-engagement. The transitions depended strongly on the reward context, and in low-reward blocks they depended more on where the animal sat along these axes.

### Two routes from orexin antagonism to disengagement

High orexin levels, as measured by oxLight, predicted disengagement, which would suggest that blocking orexin receptors should increase engagement. However, it did the opposite, a clear case of correlation not equaling causation. High orexin levels may instead be deployed to prevent disengagement, which would be higher still without them. Indeed, a large part of almorexant’s disengaging effect occurred at low arousal in low-reward trials and was not explained by the arousal changes. It occurred only in trials where orexin concentration was high but prevented from acting by the antagonist. OxLight also showed that orexin levels integrate reward over previous trials, rising when reward is scarce. We suggest that these raised levels sustain engagement when arousal drops during reward droughts, keeping mice from giving up and resting. The stimulation results also support this idea, with pupil-matched engagement higher after stimulation, and only in low reward.

The other part of the antagonist effect was mediated through arousal. Almorexant raised baseline arousal and widened its distribution, more so in low-reward blocks. Together these changes meant arousal spent more time in the range where a mouse is likely to disengage, especially in low-reward blocks, where engagement is more vulnerable to arousal. The low-reward-specific widening suggests that orexin signaling uses utility information to keep arousal stable and close to engagement-optimal levels.

Contrary to what previous work would suggest(26, 40–42, 60–63), orexin antagonism did not change dopamine signaling. Task engagement can thus be changed along the orexin and arousal axes without any dopaminergic alteration.

### Orexin-linked arousal dissociates from pupil-linked arousal

These data point to a dissociation within arousal itself. Orexin, an arousing peptide, kept engagement high when pupil-linked arousal was low. Orexin neuron activity usually correlates with pupil size(16, 17), yet orexin here acted most strongly when pupil size was low. This may be a component of arousal distinct from pupil-linked arousal, an orexin-linked arousal. We have argued that arousal should be split into components defined by the system that generates them, rather than treated as one variable indexed by the pupil(17), and the present data support that view. The classic approach of correlating neural populations with arousal in head-fixed wheel-running settings outside a task(13, 16) may therefore not translate to task settings, where these populations can diverge from pupil-linked arousal according to the needs of the task.

Another surprise was the robust increase in pupil-linked arousal under antagonism of an arousing peptide. This increase depended on the animal’s movement, on light and on the type of error it was making. Orexin can constrict the pupil through direct actions in the retina(67), but other accounts conflict(16, 17, 68–73). We saw a slight pupil decrease in darkness and no change in the pupillary light reflex or under the task lighting without the task. As such, this effect is probably due to orexin antagonism disruption of arousal control specifically in the task setting.

### Driving arousal up through orexin neurons

Raising arousal by stimulating orexin neurons also increased disengagement, and more so at low reward. Our axes attribute this mainly to the increase in pupil-linked arousal. The dilation evoked by stimulation itself depended on the reward context, being larger in low-reward blocks, consistent with gating of orexin neuron output by the animal’s valuation of the task. At 5 Hz, stimulation increased disengagement with little change in arousal, so part of the stimulation effect runs through a route we did not measure. After stimulation, by contrast, engagement in non-stimulated trials was higher than in sham sessions. This was visible only at matched pupil size, which controls for leftover pupil-linked arousal, and again only in low reward. The disengaging effect during stimulation is thus dissociable from an engaging effect after it, which may reflect slower consequences of orexin neuron activation. Stimulating the wider glutamatergic population of the lateral hypothalamus, which includes orexin neurons, raised arousal further and disengaged mice at both reward sizes, so the engagement protection at high reward holds only within a range of arousal.

### GLP-1 agonism turns down the gain of reward integration, not arousal

Semaglutide, a GLP-1 agonist, produced a large decrease in engagement, consistent with its well documented effects on nutrient-motivated behavior(43), at both reward sizes alike. In the NAcc it disrupted how dopamine integrates reward history into the tonic level. Slow tracking of reward rate by accumbal dopamine has been linked to motivation and vigor(30–32, 66). A reward-rate integrator model showed that this tracking has a gain, that the gain declines as reward accumulates and falls further under semaglutide, and that this reduction predicted the progressive disengagement. Semaglutide changed the gain of the integration but not its time constants. GLP-1 agonism therefore lowers the weight dopamine gives to recent reward, not the number of trials over which it is counted, a window that can itself change with the environment(74). The integration and its breakdown were specific to the NAcc. The DLS showed no tonic integration, and its phasic change carried little of the disengagement, so GLP-1 agonism does not act on engagement through dopamine in general. Semaglutide also compressed the phasic reward-size code in both regions. It reduced responses to high reward while responses to low reward were unchanged. This can reconcile reports of both dopamine increases and decreases under GLP-1 agonism(44– 46, 75, 76). It may also explain the effects of semaglutide on addiction(46, 47, 51, 77), if repeated use of a highly rewarding substance, like consecutive puffs of a cigarette, relies on the same tonic accumulation of accumbal dopamine. Reducing responses to highly rewarding stimuli while preserving regular reward signaling could at the same time help maintain normal everyday behavior(78, 79). A recent report found altered accumbal dopamine dynamics under semaglutide alongside reduced motivated running(46). Our data identify the gain of reward-rate integration as the affected quantity and link it to the loss of engagement. Here, lower motivation came with smaller phasic responses, whereas when motivation is lowered by cost or pre-feeding, phasic release grows(80). The two manipulations therefore likely act on different parts of the dopamine signal, the value of each reward as it arrives in one case and the weight given to the recent reward rate in the other.

Semaglutide slightly lowered pupil-linked arousal in engaged trials, which if anything should push toward engagement, and widened the pupil distribution only in disengaged trials.

The only arousal route left is through the spread of arousal in already disengaged trials, away from the range where reengagement is likely. In the mediation analysis this route was small to absent. Semaglutide thus moved engagement along the dopaminergic axis without a detectable contribution of orexin, as measured by oxLight, or of pupil-linked arousal.

### Fast DLS and slow NAcc dopamine shape engagement and arousal on their own timescales

The two dopamine measures related to engagement on different timescales. The DLS reward response tracked reward size trial by trial and predicted the next trial’s pupil, its spread and disengagement (Fig. 2), a fast coupling that carried the reward-block effect on engagement. NAcc dopamine instead integrated reward over tens of trials, and the gain of this integration predicted how engagement drifted over a session. In patch foraging, ventral striatal dopamine carries prediction errors that bias stay-or-leave decisions by reward context(33). Leaving a patch and disengaging from a task may draw on the same signal at two timescales, a phasic error that teaches what the environment is worth and a slow integration that sets whether staying is worth it now. In vehicle sessions, a larger gain also predicted a smaller decline of the phasic DLS code and a smaller rise of pupil variability over the session (Extended Data Fig. 10f), whereas the DLS response predicted pupil and engagement trial by trial (Fig. 2d–f). Dopamine transients across the striatum follow a gradient of reward time horizons, fastest in the DLS and slowest in the ventral striatum(39), and our data assign a behavioral role to each end of it. Dopamine has rarely been linked to pupil-linked arousal(81), yet here both dopamine codes shaped the variability of the pupil, the DLS from one trial to the next and the NAcc over the session, which makes the variability of arousal a shared readout of dopaminergic motivation on both timescales.

### Rules of disengagement

Our study splits task engagement along separable axes of pupil-linked arousal and dopaminergic utility coding, and shows that engagement can be maintained or lost along each of them separately (Fig. 7). For the orexin system we show that it can disengage mice through pupil-linked arousal and protect engagement through the orexin peptide, and that it uses task-utility information to stabilize arousal. We thereby link the utility-dependent stabilization of arousal to orexin signaling, a question at the forefront of arousal research(6, 17). For GLP-1 agonists we show region-specific dopaminergic effects, on the gain of reward-rate integration in the NAcc and on high-reward responses, which compress the utility code. Both GLP-1 agonists and orexin antagonists are increasingly used, so it matters how they affect engagement-driven productivity. Many neuro-behavioral disorders target the systems that maintain appropriate arousal and allow task focus, and the signatures described here may help identify and treat their disruption.

## ACKNOWLEDGMENTS

Study was funded by ETH Zurich.

## AUTHOR CONTRIBUTIONS

N.G. conceptualized the study, designed the experiments, performed surgeries, collected data, carried out data analyses and wrote the manuscript. V.P., M.H. and

E.B. collected data. A.T. and E.B. performed some of the surgeries. All authors contributed to discussion of the results and manuscript. D.B. assisted with study concept and experiment design. D.P.R. and D.B. raised funding and supervised the study.

## Methods

### Animals

All animal experiments followed Swiss Federal Food Safety and Veterinary Office Welfare Ordinance (Animal Welfare Ordinance 455.1, approved by the Zürich Cantonal Veterinary Office). Adult female and male C57BL/6 mice were studied, except for the vGlut stimulation experiments, which used vGlut2-Cre mice (Slc17a6^tm2(cre)Lowl^). Animals were housed in a 12-h reversed light–dark cycle at 22 ^*°*^C with 55% humidity. All experiments were performed during the dark phase. Animals had ad libitum access to water solution of 2% citric acid (82), which allowed stable motivation for 10% sucrose reward over the course of experiment, and ad libitum access to normal food (chow).

### Surgeries and viral vectors

For stereotaxic surgeries, mice were anesthetized using 2–5% isoflurane and administered analgesia using buprenorphine and site-specific lidocaine. We injected the dopamine sensor ssAAV-9/2-hSyn1-chl-dLight1.3b-WPRE-bGHp(A) (200 nL, 7.9 × 10^12^ vg/mL, UZH Viral Vector Facility) in the dorsolateral striatum (DLS, AP: +0.8 mm ML: 2.2 mm DV: 3.0 mm), and medial nucleus accumbens shell (NAcc, AP: +1.3 mm ML: 0.5 mm DV: 4.25 mm). In the same mice we also injected orexin sensor AAVDJ.hSynapsin1.OxLight1 (200 nL, 7 × 10^12^ vg/mL, UZH Viral Vector Facility) into the substantia nigra pars compacta (SNc, AP: -3.2 mm ML: 1.4 mm DV: 4.1 mm) and the ventral tegmental area (VTA, AP: -3.2 mm ML: 0.5 mm DV: 4.6 mm). Injections were counterbalanced for side between mice. We implanted 200 µm diameter optic fibers, 0.39 numerical aperture (NA) with a 1.25 mm ceramic ferrule (Thorlabs) 0.1 mm above each of the injected areas, and implanted the head-fixation bar, fixating everything with dental cement (C&B Metabond).

For the optogenetic experiments, animals were injected with AAV9-hORX-ChrimsonR-mCherry (200 nL, 2 × 10^12^ vg/mL, UZH Viral Vector Facility), with previously verified orexin promoter to selectively target orexin neurons at the given concentrations (23, 83). In this case the injections were performed bilaterally in the lateral hypothalamus (LH, AP: -1.35 mm ML: ±0.95 mm DV: 5 mm). Fibers were implanted 0.15 mm above the injection sites. The vGlut experiments used the same surgery in vGlut2-Cre mice, with 200 nL of AAV9-EF1a-DIO-ChrimsonR-mRuby2-KV2.1-WPRE-SV40 (124603-AAV9, Addgene, stock 1 × 10^13^ vg/mL diluted 20-fold to 5 × 10^11^ vg/mL) in place of the orexin construct. The heat controls had no virus injections, just the fiber implants in the same location.

### Histology

Animals were terminally anesthetized using pentobarbitone and perfused with a sterile PBS solution at pH 7.4 followed by 4% paraformaldehyde in PBS. Brains were removed and then kept in a 4% paraformaldehyde solution overnight, and then in 30% sucrose for another night. Brains were frozen using dry ice and then sectioned at 50 µm with a cryostat. Images were acquired using a fluorescence microscope (Eclipse Ti2, Nikon).

### Behavioral task

The task was based on the International Brain Lab standardized visual decision making task (53), with some modifications. Namely the contrasts were changed to make the task somewhat more difficult to 3%, 6% and 12.5% contrasts only. Additionally we introduced a block structure similar to other previous work (5) where we modulated reward size per block every 60 trials to be either 1 µL or 5 µL, always starting with a low reward block.

The trial begins with a quiescent period in which the headfixed animal is not allowed to move the wheel under its forepaws for a duration of between 0.5 and 0.7 seconds. Following, there is a brief 12 kHz sound signaling trial start, and the stimulus (a sinusoidal grating) appears on the left or right side of the screen in front of the animal (50% chance of either side). The animal uses the wheel, which then becomes locked to the position of the stimulus on screen, to indicate its belief about which side the stimulus is on. Moving the stimulus to the center of the screen is the correct outcome and it leads to a drop of 10% sucrose being delivered to the spout in front of the animal’s mouth. The wrong outcome leads to a longer inter-trial interval (2 seconds instead of 1-1.5 – not including the quiescent period) and a white noise being played to the animal. Maximum of 420 trials were run per training session in the reward task, while the constant reward, contingency switching task ran till up to 570 trials. There were conditions that led to earlier termination of the session to prevent overt stress to animals that were no longer motivated to perform the task. The session was terminated after 45 minutes if less than 25 trials were performed in the last 5 minutes, or if the session took longer than 90 minutes.

### Pharmacological manipulations

Animals were always injected i.p. 45 minutes before being placed into the setup and starting the session. For dual orexin antagonist Almorexant (MedChem Express), it was dissolved in vehicle (used for control) of 2% dimethyl sulfoxide and 25% beta-cyclodextrin in PBS, to achieve an injection volume of about 125 µL at 30 mg/kg. Semaglutide was dissolved just in PBS (used as control vehicle) to achieve an injection volume of about 100 µL at 0.03 mg/kg.

### Fiber photometry

Fiber photometry was recorded at 20 Hz using a multifiber camera-based photometry system (Doric) with illumination that was alternating between 405 nm and 470 nm excitation. For the dopamine (dLight) channels, photobleaching and motion artifacts were removed using the isosbestic control: within each session the 470 nm trace was linearly regressed onto the simultaneous 405 nm trace, and ΔF/F was computed as (F470 − F_0_)/F_0_, where F_0_ is the 405-predicted 470 signal. For the orexin (oxLight) channels, which lack a usable isosbestic, the slow bleaching baseline of the 470 nm trace was estimated by a sum-of-exponentials fit and removed (biexponential baseline for ΔF/F). ΔF/F was divided by one number per mouse and drug experiment, the standard deviation of the per-trial pre-stimulus level (mean 1 to 0.05 s before stimulus onset) over all of that mouse’s trials, pooled across sessions and reward blocks (z); because the same divisor applies to drug and vehicle sessions, it leaves each mouse’s drug/vehicle ratio unchanged and weights the mice equally. Absolute response sizes were compared between drug and control sessions in ΔF/F on brightness-matched sessions (see below). Recording sites without a stimulus- or reward-aligned change in a session were identified by visual inspection and excluded from all analyses for that session. Event-locked reward responses (Fig. 6f,g, Extended Data Figs. 6i and 10b) were the reward-aligned trace minus the trial’s pre-stimulus level (0.25–0 s before stimulus onset), averaged per mouse, drug and reward size over rewarded trials (at least 10; both engagement states), in z. Fig. 6f,g and Extended Data Figs. 6i and 10b compared drug with vehicle on the AUC over−0.5 to 1.5 s (NAcc) and −0.5 to 1.0 s (DLS, due to quicker responses) after reward (paired t-tests over 8 mice).

### Pupil size recordings

Mouse pupils were recorded during the entire experiment using an infrared camera (Blackfly S USB3 Teledyne FLIR) at 20 Hz with a custom written code for camera acquisition in Python using SpinnakerSDK. The camera capture was synchronized via an output line that generated a square wave representing each frame. Pupil size was estimated by extracting 8 edges around the edge of the pupil with DeepLabCut (84) and a circle was fitted to these points with custom code in Matlab. In case of low-confidence point estimates of some points, down to 5 points with the highest estimates were chosen for fitting. In case less than 5 points fit the criteria for fitting the circle (<95% likelihood reported by DeepLabCut), such as during blinks, the datapoints were interpolated. Sessions in which tracking failed were identified by an objective criterion (more than 4% of trial baselines further than 5 robust standard deviations from the session median, or a session standard deviation more than 2.5 times the robust standard deviation), and their pupil data were excluded (3 of 246 reward-task sessions); behavior and photometry from these sessions were kept. The baseline pupil of a trial was the mean diameter 0.25–0 s before stimulus onset. For the arousal analyses it was residualized within each session on the previous trial’s contrast and outcome (keeping the session mean) and z-scored within mouse over the drug and control sessions of a drug pair, so that drug-induced shifts are preserved.

### Optogenetic stimulation

The optogenetic stimulation was performed using 635 nm lasers (Laserglow) at 10 mW power at the fiber end. The stimulation schedule was randomized to either 5 Hz or 20 Hz with 5 ms pulses applied in 30-s bouts separated by 60 s without light (90 s from one bout onset to the next; the first bout about 60 s after the first trial) throughout the session, as read from the recorded laser channel (n = 8). Heat controls (n = 7) received light with the same parameters, without virus, in one 30-s bout about every 5 min. LH glutamate-neuron (vGlut) mice (n = 5) were stimulated at 5 and 20 Hz in 30-s every 5 minutes. Each orexin mouse also ran sessions without laser, used for the sham comparison (below).

### Engagement-state HMM (RT-HMM)

To infer trial-by-trial task engagement we fit a two-state switching linearregression hidden Markov model to reaction times, following Johnson et al.(6). The latent state z_*t*_ ∈ {engaged, disengaged} followed a first-order Markov chain (initial distribution *π*, transition matrix A), and the log reaction time was modeled as a state-specific linear function of the trial’s stimulus properties: y_*t*_ = log RT_*t*_ ∼ *N* (x_*t*_*β*_*k*_, *σ*_*k*_^2^), with x_*t*_ = [1, contrast, side] (side coded ±1). Each state had its own intercept, contrast/side sensitivity, and residual variance, capturing the fast, low-variance engaged regime and the slow, high-variance disengaged regime typical of extended rodent sessions.

Reaction times were log-transformed (capped at 60 s beforehand to bound timed-out trials); trials with missing or nonpositive RT were excluded and left unassigned. Parameters were estimated by maximum likelihood using EM (Baum– Welch), with forward–backward recursions in the log domain and a small ridge term (10^−6^) on the regression update, run to a log-likelihood tolerance of 10^−4^ (≤200 iterations) and repeated from 20 random initializations (highest-likelihood solution retained). States were labeled by emission intercept (smaller = engaged), and each trial assigned to its most probable state (MAP).

To prevent the definition of “engaged” from drifting between sessions or drug conditions, we fit one shared emission/transition model per mouse × drug-pair: all of a mouse’s sessions under a drug and its matched control (semaglutide/vehicle or almorexant/vehicle) were fit jointly, each session contributing its own Markov chain (no transitions across session boundaries) but sharing {*β*_*k*_, *σ*_*k*_}, A, and *π*. This enforces a single, drug-agnostic state definition within each animal, so drug-versus-control differences reflect changes in state occupancy or dynamics rather than in how the states were defined. Hard state labels were merged into the trial table and used to split all behavioral and photometry measures by engagement state. Implemented in Python (NumPy/SciPy).

### Mediation analysis of drug/stimulation effect on engagement via arousal

Mediation of disengagement by arousal. For each drug pair and reward level, the two transition hazards were estimated on contiguous trial pairs – the disengage hazard P(disengaged(t) | engaged(t ™ 1)) and the re-engage hazard P(engaged(t) | disengaged(t ™ 1)) – as logistic functions of the baseline pupil of trial t (pupil and pupil^2^, each also interacting with a drug indicator) and of the trial’s position in its session (z-scored within session) and its interaction with drug, with a random intercept and random pupil and pupil^2^ slopes per mouse × condition, so that each condition’s hazard curves are estimated from its own sessions (Fig. 4b). From the fitted hazards we computed the steadystate disengaged occupancy *π*D = āED / (āED + āDE), where āED and āDE are the mean hazards over a set of trials. The arousal-mediated (pure indirect) effect was the change in *π*D when the vehicle hazards are evaluated on the drug sessions’ trials instead of the vehicle sessions’ trials (each trial keeping its own session position), i.e. the drug’s pupil distribution under the vehicle hazard curves; the disengage and the re-engage route swapped the pupil distribution in only one of the two hazards, so the two routes need not add up to the total. The total drug effect was computed from the hazards of each condition; all quantities were expressed per mouse as a percentage of that mouse’s total drug effect and averaged across mice (Fig. 4d; 95% confidence intervals and one-sample ttests across mice).

Because this model conditions on the RT-HMM engagement states, we confirmed the mediation generatively by refitting the reaction-time HMM with the transition probabilities driven directly by baseline pupil while the emissions (and thus the state definitions) remained governed by reaction time (Extended Data Fig. 5a); the inferred states were 97.9% identical to the original. The transition into trial t was a logistic of the baseline pupil of trial t and its square, reward, drug, and the trial’s position in its session (z-scored within session) with its interaction with drug. Two estimators are reported: a per-mouse fit, in which × drug and reward shifted the intercepts, tested with exact Wilcoxon signed-rank tests; and a hierarchical fit with the pupil terms × drug pooled across mice and fitted separately per reward block, with one intercept per mouse × condition, whose 95% intervals and P values came from a percentile bootstrap over mice (1,000 resamples; P from the fraction of resampled estimates on either side of 0, floored at 0.001). The indirect effect was the pure indirect effect described above. Each fit was started from 12 random initializations. For Fig. 4e (right) and Extended Data Fig. 6b, both hazards were fitted in low-reward blocks within each mouse’s bottom or top 30% of baseline pupil, with the oxLight baseline of trial t and its interaction with drug, the pupil and the trial’s position (*×* drug) as inputs (ridge-penalized logistic regression, one intercept per mouse × condition); P(disengaged) = a/(a + b) was evaluated along the oxLight axis per mouse and averaged, with 95% intervals from a cluster bootstrap over mice (2,000 resamples), and the growth of the almorexant ™ vehicle difference was taken from ™1 to +1.5 z. In Fig. 4g, the NAcc and DLS measures of Fig. 2 (last + running; z per mouse and experiment) entered the pure-indirect decomposition in place of the pupil, on the trials with that measure, and the pupil share was recomputed on the same trials.

### Mice, sessions and trials

Almorexant and vehicle were tested in 13 mice: 8 with fiber photometry and pupil recordings and 5 with pupil recordings only. Semaglutide and its vehicle were tested in 8 mice with fiber photometry and pupil recordings, 7 of which also took part in the almorexant experiment. Each mouse received each drug and its control on different days. Sessions with an accuracy below 0.65 over the first 420 trials were excluded, which left 112 almorexant, 127 vehicle (almorexant experiment), 65 semaglutide and 53 vehicle (semaglutide experiment) sessions. In 9 semaglutide sessions photometry was not recorded (failed synchronization signal or LED); these sessions entered the behavioral and pupil analyses only. Analyses of control sessions alone (Fig. 2) pooled the vehicle sessions of both experiments within each mouse (14 mice with pupil, 9 with photometry, 8 with SNc). The constant-reward task (Extended Data Figs. 3e–h and 9a) used 8 mice.

### Engagement measures

P(engaged) was the fraction of trials in the engaged state. Transition hazards were computed on consecutive trials of a session: the disengage hazard, P(disengaged on trial n + 1 | engaged on trial n), and the re-engage hazard, P(engaged on n + 1 | disengaged on n), assigned to the reward block of trial n. Block-by-block drug ™ control differences (Fig. 3c) were computed per session and block and then averaged over sessions. Accuracy was computed on trials with a response. Disengaged trials were classified by reaction time as anticipatory (< 0.3 s), slow (0.3–10 s) or lapses (> 10 s or no response; Extended Data Fig. 6f–h).

### Signals and engagement (Fig. 2)

Each signal entered as one pre-specified measure: for pupil, SNc and VTA the prestimulus level of trial n (mean 0.25–0 s before stimulus onset, residualized per session on the previous trial’s contrast and outcome); for NAcc and DLS the sum of the z-scored reward response of the previous trial (mean 0.1–1 s after reward minus that trial’s pre-stimulus level; defined when the previous trial was rewarded and immediately preceded trial n) and the z-scored running average of the reward responses up to trial n ™ 2 (exponentially weighted over the earlier rewarded trials, time constant 5 rewarded trials), z-scored again (‘last + running’). Hazards were the transition into trial n (the state of trial n given the immediately preceding trial), so that the levels of trial n are measured before the state they predict. Dopamine was used as ΔF/F (405 regression), oxLight as the 470-nm trace minus a four-exponential bleaching fit, zscored per session; every measure was z-scored within mouse and experiment over all sessions.

For the curves, each mouse’s trials were cut into six quantile bins of the signal, separately within each reward block, and the rate computed per bin and reward block (at least 5 trials). Per mouse and reward block, a linear and a quadratic orthogonal-polynomial term were fitted to the binned rates; each term was tested against 0 across mice (two-sided t-tests; two contrasts per panel, uncorrected). The linear term tests a monotone trend, the quadratic term a bend (for example a peak inside the range). The same conclusions followed from per-mouse logistic slopes on single trials, a random-slope mixed model, rank correlation over bins and an any-shape test (rate ∼ bin + (1|mouse)). The traces across the session and around switches (Fig. 2a,c, Extended Data Fig. 2a,c,d) are means over each mouse’s control sessions, then over mice; the reward-size test in the distribution panels compared each mouse’s mean in high- and low-reward blocks (paired ttest). For the maps (Fig. 2b, Extended Data Fig. 2b), the probability of the transition was smoothed per mouse over the two signals with a Gaussian kernel (bandwidth 0.45 z for disengaging and 0.6 z for re-engaging; points with less than 4 or 3 trials of kernel weight left out) and averaged over mice where every mouse contributed (low-reward blocks). The slope on recent reward (Extended Data Fig. 2g) regressed each signal’s pre-stimulus level on a leaky average of the reward obtained (time constant 10 trials) within session. Per-trial correlations (Extended Data Fig. 2e) were computed per mouse and experiment over trials with both measures (at least 100), Fisher-z averaged per mouse, separately on engaged and disengaged trials (state of trial n), and tested against 0 across mice (Benjamini–Hochberg over the 10 pairs of each state). For the block mediation (Fig. 2d, Extended Data Fig. 2f), logistic models of the two transitions were fitted per mouse with reward block, trial position and log run length as covariates (Extended Data Fig. 2f: also wheel movement before the stimulus, movement and licking around the previous reward and their running averages); each signal’s values in high-reward blocks were replaced by the same quantiles of its low-reward distribution, and P(disengaged) = a/(a + b) was recomputed from the two hazards. The share of the block effect was the ratio of means over mice, with a 95% bootstrap interval over mice and a one-sample t-test across mice on the indirect part; all five signals entered one model with missing measurements set to 0 plus a missing indicator.

For the within-block pupil spread of Fig. 2d, the outcome was each trial’s absolute deviation of the baseline pupil from the mean of its block instance (session × 60-trial block), modeled per mouse by least squares with reward block, trial position and experiment; the indirect part was computed as above. Fig. 2d shows the shares on the trials after a rewarded trial (where the NAcc and DLS measures exist); Extended Data Fig. 2f and the all-trials values in the legend of Fig. 2d were fitted on all trials, with a missing measure set to 0 plus a missing indicator, so that every row is a share of the same effect. Fast coupling (Fig. 2e): per mouse, the next trial’s baseline pupil, its absolute deviation from the mean of its block instance within engagement state, and the disengage indicator (trials after an engaged trial) were regressed by least squares within session on the previous trial’s reward response (last + running, per within-mouse SD), with reward block, trial position and a last-block step as covariates and one intercept per session; DLS and NAcc were fitted separately on the same trials (8 mice with both), and the slopes were tested against 0 and against each other across mice (one-sample and paired ttests). Controls: one offset per block instance in place of the block covariate (only fluctuations within a block), and the engagement states of the two trials as covariates (or only trials engaged on both).

Interactions (Extended Data Fig. 2h): per mouse, the two hazards were fitted with all five measures, their ten pairwise products, reward block, trial position and log run length (ridge-penalized logistic regression; missing measures set to 0 with a missing indicator), separately for low and high reward blocks; each product term was tested against 0 across mice (two-sided t-tests), Benjamini–Hochberg corrected over the ten pairs within each hazard and block. High-reward disengaging is the rarest transition, so these terms rest on the fewest events.

### Pupil level and variability (Figs. 4 and 6h, Extended Data Fig. 5c)

The mean baseline pupil was averaged over trials per mouse, reward block and engagement state. Within-block variability was the pooled standard deviation of the baseline pupil after subtracting each block’s own mean, computed per mouse, reward block and engagement state over all blocks of that reward size (sum of squares divided by N ™ B for N trials in B blocks), so that slow changes between blocks do not count as variability. P(engaged) against baseline pupil used six quantile bins of the pupil per drug pair, at least 5 trials per mouse and bin, and paired tests per bin. The change of the baseline over the session (Extended Data Fig. 4a) was the slope on trial number per mouse and drug, with one intercept per session. The detrended baseline (Extended Data Fig. 4a, right) was the baseline minus each session’s linear trend in trial number, keeping the session mean. For Extended Data Fig. 6a, each mouse’s trials (both drugs) were split at the 30th and 70th percentiles of its baseline pupil, and P(engaged) was computed in five quantile bins of the oxLight baseline within each pupil group; bins were compared between drugs with paired t-tests (uncorrected).

### Accumbal value code and disengagement (Fig. 6e, Extended Data Figs. 9 and 10)

The utility codes of a session (Extended Data Fig. 9c,d) were computed on correct trials more than 10 trials into a block: the tonic code from the pre-stimulus level (mean 1 to 0.05 s before stimulus onset), the phasic code from the reward response (0.1–1 s after reward minus the trial’s pre-stimulus level), each as block 2 (high reward) minus the mean of blocks 1 and 3 (Extended Data Fig. 9c) or block 2 minus block 1 (Extended Data Fig. 9d). Late disengagement was P(disengaged) from block 3 on (Extended Data Fig. 9d) or block 4 on (Extended Data Fig. 9h). Per mouse and drug (at least 3 sessions), P(disengaged) was regressed on the session measure by least squares; r was the correlation within mouse and drug, or within mouse over both drugs, Fisher-z averaged and tested across mice. The early value code of Extended Data Fig. 9b,h was the reward response (0–1 s after reward minus the pre-stimulus level) in the first high-reward minus the first low-reward block (at least 1™ 0 correct trials each).

The early-code comparison (Extended Data Fig. 9b) averaged the early value code, and the same high low difference over blocks 3 onwards, per mouse.

### Comparisons of dopamine between sessions (Extended Data Fig. 9e–g)

Because ΔF/F and z-scores are relative to each session, drug and control sessions were compared on brightness-matched pairs. The raw 470-nm light of each NAcc fiber was taken as the session median, and each semaglutide session was paired one-to-one with a vehicle session of the same mouse whose light differed by less than 10% (optimal assignment; rule fixed before any response was computed). Responses were computed from the raw 405- and 470-nm traces with a 405-nm fit within each reward block (first 10 trials of each block dropped), as ΔF/F 0.1–0.6 s after reward minus the pre-stimulus level, on correct trials. The tonic code (Extended Data Fig. 9g) was the high ™ low difference of the pre-stimulus level (ΔF/F, without baseline subtraction); Extended Data Fig. 9f shows the high ™ low difference of the reward-aligned responses on the same pairs, and Extended Data Fig. 9e the low- and high-reward responses.

### Reward integration by tonic dopamine (Fig. 6a–d, Extended Data Fig. 8)

These analyses used NAcc and DLS in z (see Fiber photometry). The tonic level of a trial was the mean NAcc signal from 1 to 0.05 s before stimulus onset. Around each reward-size switch, the reward-aligned traces of the pre-switch trial and of the 1st, 2nd, 3rd, 10th and 30th trial of the new block (rewarded trials) were taken relative to the pre-switch trial’s tonic level and averaged per mouse over switches (at least 3); the reward AUC was the area 0–3 s after reward (Fig. 6a; Extended Data Fig. 8a,b). For Fig. 6c, each session’s tonic levels were centered on the session mean and averaged by trial number over all sessions. The two-timescale reward-rate integrator (Fig. 6b,c) modeled the tonic level of trial n as B(n) = b + g·V_f(n) + h· [V_f(n) V_s(n)] + {g_t·V_f(n) + h_t·[V_f(n) ™ V_s(n)] + c} · z(n), with b a session intercept, V_f and V_s exponentially weighted averages of the reward obtained (µL) on the session’s earlier trials with a fast (1–10 trials) and a slow (10–80 trials) time constant, and z(n) the trial’s position (z-scored within session); the time constants were chosen on a grid and the weights fitted by least squares per mouse and drug, each session being predicted from the mouse’s other sessions. The fitted time constants did not differ between vehicle and semaglutide (fast 5.5 vs 4.8 trials, P = 0.36; slow 20.0 vs 27.5 trials, P = 0.43; medians over mice, signed-rank tests). The two integration terms over the session (Fig. 6b) are g + g_t·z(n) and h + h_t z(n). The response to a reward step (Fig. 6d; almorexant, Extended Data Fig. 8f) was the model’s tonic level for the task schedule with every trial rewarded (1 µL in low, 5 µL in high reward blocks, 420 trials), computed with the mouse and drug’s fitted terms, minus its level on the trial before a switch into high reward, averaged over trials 1–30 of the new block, for the 1st (trial 61) and the 3rd switch (trial 301). Integration within a block (Extended Data Fig. 8c) used trials more than 10 into a block: the tonic level minus its block’s mean was regressed, per mouse, drug and reward size, on the number of rewarded trials among the 2nd to 5th preceding trials of the same block, with the position in the block as a covariate. The inside-block kernel (Extended Data Fig. 8d,f) regressed the tonic level on the reward obtained on each of the previous 10 trials of the same block, with one intercept per block, on trials more than 10 trials into a block; its summary was the summed weight of lags 2–5. For confidence weighting (Extended Data Fig. 8e), the lagged rewards were split by the stimulus contrast of their trial and compared with the weights expected from an integrator of prediction errors whose expectation is the mouse’s accuracy at that contrast times the reward size.

Prediction errors at switches (Extended Data Figs. 8g,h and 10c–e) were taken on the phasic response (0.1–1 s after reward minus the 0.25 s before it; DLS responses corrected for stimulus contrast by subtracting each contrast’s mean response more than 10 trials into a block): the switch step was the response to the first reward of a new size minus that to the last reward of the old block, and the surprise the first reward minus the mean of the second and third, with the sign reversed for switches into low reward and both directions averaged per mouse. The DLS model (Extended Data Fig. 10e) described the response to each rewarded trial as a + b_size·(U ™ V) + b_conf (1™ p)·V, where U is the value of the reward size (1 for 1 µL, u for 5 µL), V an expected value updated after each reward with learning rate *α* (V = 1 at the session start) and p the mouse’s accuracy at the trial’s contrast; u and *α* were fitted on a grid and the weights by least squares per mouse and drug, with session intercepts; the reward-size prediction error at a switch was b_size (u 1). Cue value and cue confidence were the high ™ low rewardblock and easy ™ hard contrast differences of the cue response (0.1–0.5 s after stimulus onset minus the pre-stimulus level), all on correct trials more than 10 trials into a block. Changes under semaglutide (Extended Data Fig. 10g) were 100 × mean(semaglutide ™ vehicle) / mean(vehicle) over mice, with 95% intervals from a paired bootstrap over mice (10,000 resamples).

For the early step response per session (Fig. 6e), the integrator of Fig. 6b,c was refitted to each session (at least 40 trials with a tonic level in blocks 1–2), with the fast time constant of its mouse and drug, the slow time constant fixed at 20 trials, a fast gain free in each third of the session (blocks 1–2, 3–5 and 6–7) and a recent-minus-slow gain linear in trial position; the step was the model’s tonic level on the task schedule with every trial rewarded, mean of trials 61–90 minus trial 60 (the first switch into high reward), from the blocks 1–2 gains. The outcome was the session trend of P(disengaged): the coefficient of trial position (z-scored, trial 1 to 420) in a least-squares fit of the disengaged state (0/1) on reward block and trial position, in sessions with at least 100 state-labeled trials, reported as the slope *×* 3.45, the change along the fitted line from trial 1 to 420. Session values were centered within mouse and condition; r was the pooled within-mouse correlation, with P from a within-mouse permutation (10,000). For Extended Data Fig. 10f, the same step was related, over the same vehicle sessions and with the same trend fit and centering, to two session trends, the phasic DLS reward response (0.1–1 s after reward minus the trial’s prestimulus level, rewarded trials, z) and the within-block SD of the baseline pupil (per 60-trial block with at least 20 trials, fitted on reward block and the block’s mean trial position, in sessions with at least six blocks). The share of semaglutide’s effect on the trend carried by the step was a × b / total, with a the semaglutide ™ vehicle difference in the step and the total the difference in the trend (mouse means, 8 mice), and b the within-mouse slope of the trend on the step in the vehicle sessions of both experiments (lower bound) or in the semaglutide sessions (upper bound); their mean is reported, as for the arousal routes. 95% intervals and P came from a cluster bootstrap over mice that refits the links (5,000 resamples).

For Fig. 6g, the disengage and re-engage hazards on blocks 3 onwards were fitted per mouse and drug by ridge-penalized logistic regression (intercept, reward block, trial position, log run length, the session’s NAcc step and the DLS measure last + running with a missing indicator); the DLS values of one drug were mapped onto the other drug’s distribution by quantile within mouse and reward block, and P(disengaged) = a/(a + b) was recomputed: vehicle hazards with semaglutide’s DLS distribution (vehicle rule, the pure indirect effect, lower bound), semaglutide hazards with vehicle’s distribution (semaglutide rule, upper bound) and their mean, as shares of the semaglutide ™ vehicle difference in P(disengaged), with 95% intervals and P from a bootstrap over mice (5,000 resamples). The main panel shows the disengage hazard by quintile of the DLS measure (edges per mouse and reward block over both drugs, the two block types averaged), and *β* is the DLS coefficient of the same hazard fits in log-odds per within-mouse SD.

### Stimulation analyses (Fig. 5, Extended Data Fig. 7)

Trials were labeled from the recorded laser channel. A trial was stimulated when the laser was on at stimulus onset; the trial during which the laser switched on was excluded; control trials started at least 45 s after the end of the last bout and had no laser during the whole trial. Because these rules favor short trials (a long trial is more likely to overlap a laser edge), the same labels were applied to sham bouts: each mouse’s laser schedules laid over its sessions without laser, with the same timing relative to the first trial. Sham sessions were restricted to days on which the reward task ran. Stimulation effects are reported as laser sham, per mouse. Matched-pupil comparisons (Fig. 5e, Extended Data Fig. 7b) compared stimulated and control trials within pupil bins. Dilation by reward block (Extended Data Fig. 7c,d) was the laser ™ sham change of the stimulated ™ control baseline in low- and high-reward blocks, and the continuous pupil 0–30 s after bout onset minus the 30 s before it. For vGlut stimulation (Extended Data Fig. 7i), the pupil slope over the last second before each stimulated trial’s stimulus was related to the trial’s time into the bout. For post-stimulation trials (Extended Data Fig. 7f), every laser-free trial starting 0–60 s after the end of a bout was matched to all sham trials of the same mouse and reward block whose baseline pupil (residualized, and z-scored once per mouse over laser and no-laser sessions, so that differences between days are kept) was within 0.05 z; the difference in P(engaged) was averaged per mouse (the 8 orexin mice of Fig. 5b–e). The arousal routes (Extended Data Fig. 7e) used the hazard model of the drug mediation (pure indirect effect), with stimulation in place of the drug, expressed as % of the observed laser ™ sham change in P(disengaged) (ratio of means over mice; 95% bootstrap interval over mice; paired t-tests on the absolute routes; high-reward routes from the 7 mice with enough trials).

### Light test (Extended Data Fig. 4i)

In part of the stimulation cohort, the setup light was switched off for about 2 min and then on for about 2 min before the first trial and after the last trial of each session in the final weeks of the experiment (63 sessions). Lights-on was detected from the pupil step. DARK was the mean pupil ™70 to ™10 s and LIGHT +50 to +110 s from lights-on (stopping at the first trial); the pupillary unrest index was the summed absolute change between consecutive 0.65-s block means per minute; t50 was the time to half of the constriction after lights-on (or of the dilation after lights-off). Values were expressed in units of each mouse’s task baseline (z with the mean and standard deviation of its laser-free task baselines over all sessions).

### Statistics. Figure conventions

Unless a legend states otherwise, n is the number of mice and all values were averaged within mouse before averaging across mice. Error bars and bands are mean ± s.e.m. across mice, or 95% intervals where the legend says so; thin lines or small dots are individual mice; black crosses are mean ± s.e.m. Drug and control were compared with two-sided paired t-tests across mice, and per-mouse slopes or differences with one-sample t-tests against 0. ^*^ P < 0.05, ^**^ P < 0.01, ^***^ P < 0.001, n.s. not significant. Every statistical test shown on a figure panel is listed in Supplementary Table 1, with the test used, n, the test statistic and the exact P value. Where a test uses fewer mice than the legend states, because a signal is missing in a mouse or a mouse has too few trials of that kind, the n of that test is the one given there. In the engagement curves, stars mark a linear trend across bins and ∩ or U a downward or upward curvature (orthogonal-polynomial terms per mouse, t-tests across mice, uncorrected). Dopamine and oxLight units are given in Fiber photometry. Vehicle colors are blue, almorexant magenta, semaglutide green, orexin stimulation red (20 Hz) and orange (5 Hz), heat controls gray, vGlut purple. In Fig. 2 the DLS dLight traces are orange. Repeated-measures ANOVAs used mouse as a random factor. Per-mouse generative estimates were tested with exact Wilcoxon signed-rank tests (Hodges–Lehmann estimates with exact 95% intervals). Pooled within-mouse slopes were tested with the wild cluster bootstrap-t, and pooled model quantities with bootstraps over mice (percentile intervals; two-sided P from the fraction of resampled estimates on either side of 0). Mixed-effects logistic regressions had a random intercept per mouse. No correction for multiple comparisons was applied unless stated (Benjamini–Hochberg in Extended Data Fig. 2e,h). Analyses were done in MATLAB R2023a (Statistics and Machine Learning Toolbox) and Python 3 (NumPy, SciPy).

## Extended Data

**Extended Data Fig. 1.**
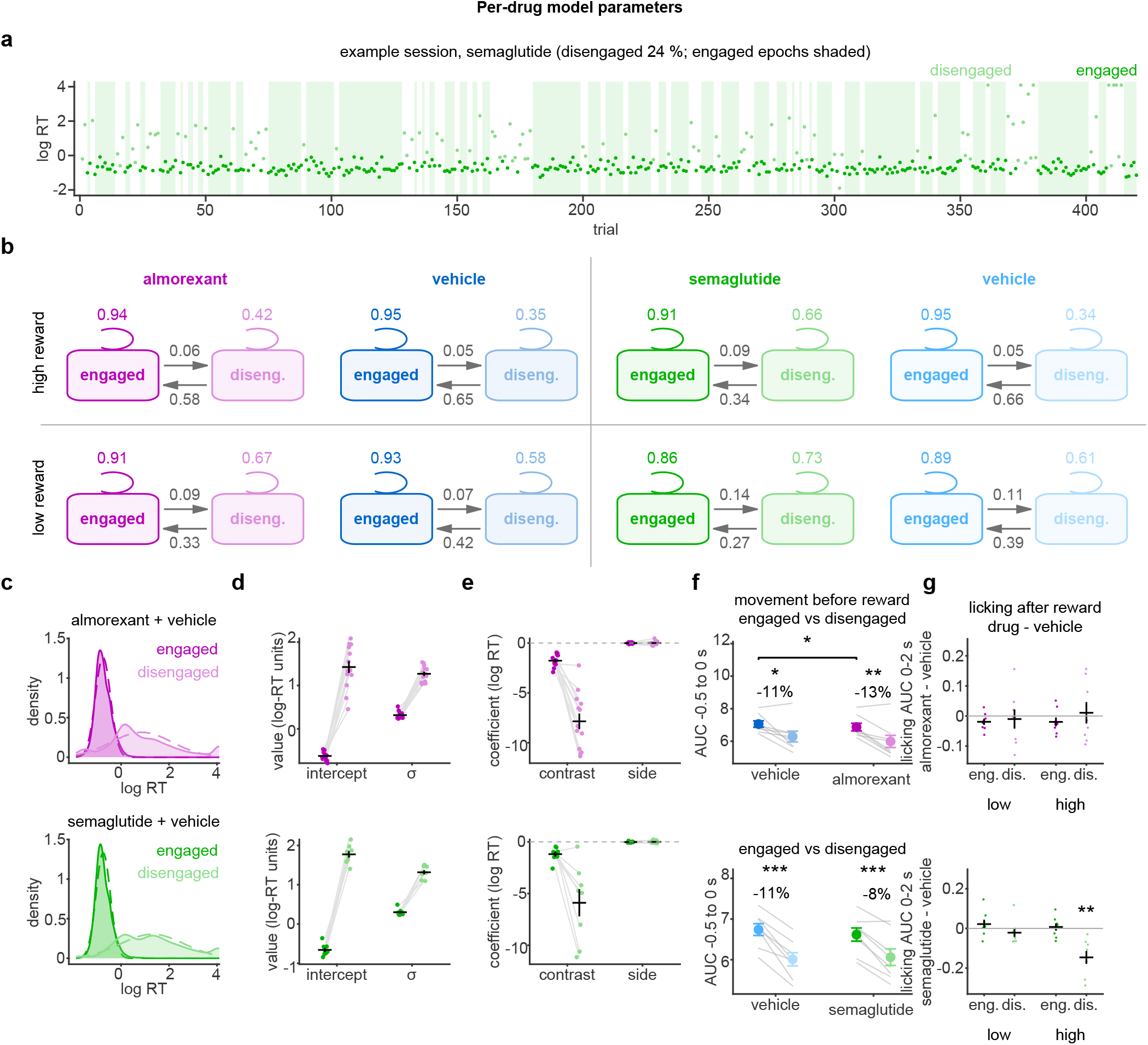
Engagement states: model fits and behavior. **a**, Example session (semaglutide); log reaction time per trial colored by the inferred state (shaded, engaged epochs). **b**, Transition probabilities per drug condition and reward size (consecutive trial pairs within a block, pooled over mice). **c**, Distributions of log reaction time in the engaged and disengaged state (dashed, fitted Gaussians). **d, e**, Emission intercept and SD (**d**) and contrast and side slopes (**e**) per state. **f**, Wheel movement before reward (AUC −0.5 to 0 s), engaged vs disengaged within each condition (top, almorexant experiment; bottom, semaglutide experiment; text, % change; brackets, drug vs vehicle in the same state where P < 0.05). **g**, Licking after reward (AUC 0–2 s), drug − vehicle per mouse in low and high reward blocks on engaged (eng.) and disengaged (dis.) trials (top, almorexant; bottom, semaglutide); stars, one-sample t-tests against 0. Model details in Methods. n = 13 mice (almorexant/vehicle) and 8 mice (semaglutide/vehicle); f, g, 8 mice per pair with wheel and lick recordings. Statistics in Supplementary Table 1.

**Extended Data Fig. 2.**
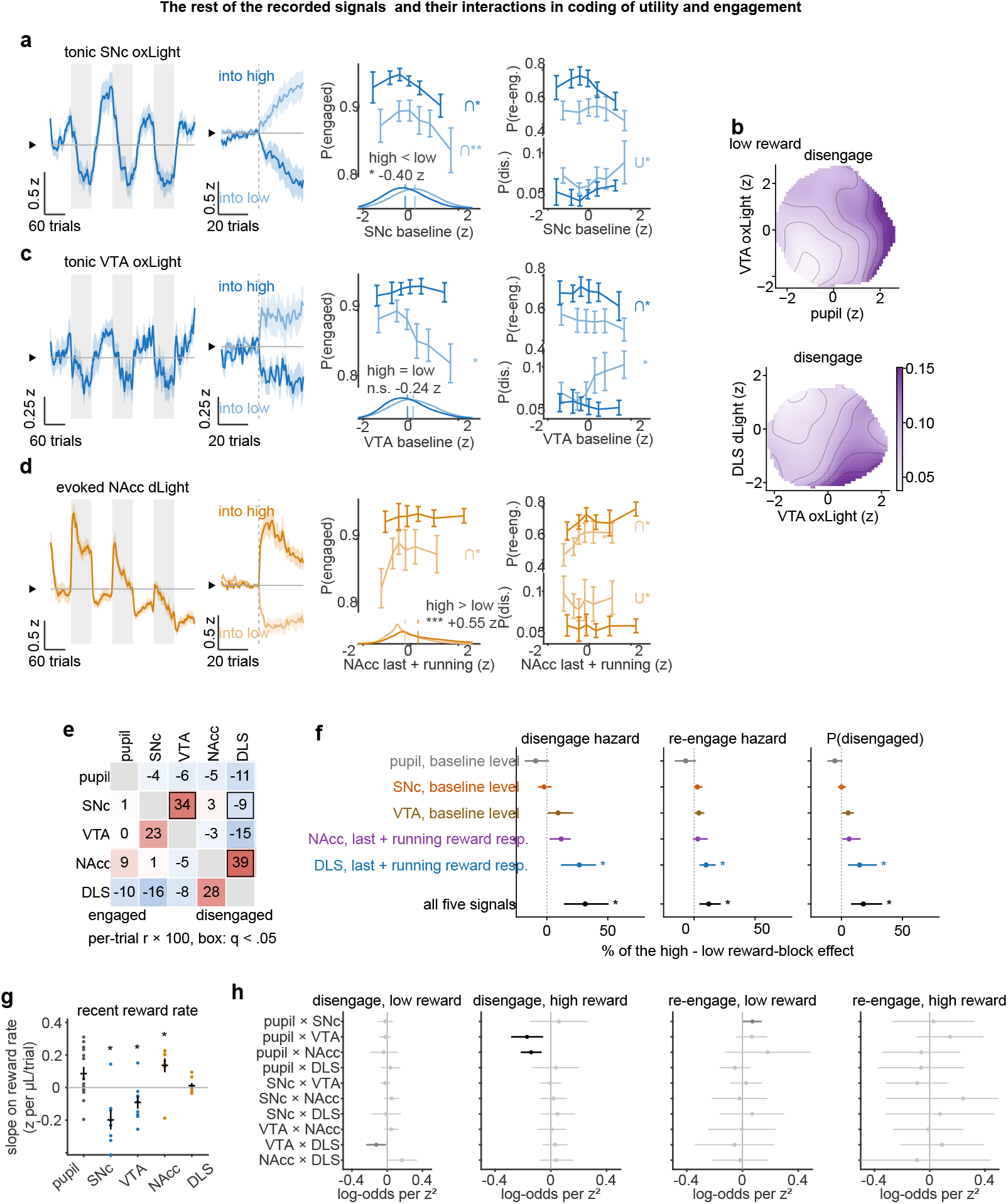
The other recording sites, correlations between the signals and the block mediation with movement and licking, and pairwise interactions. **a**, SNc oxLight baseline, as Fig. 2a. **b**, Disengage probability in low-reward blocks over pupil *×* VTA (top) and VTA *×* DLS (bottom), as Fig. 2b. **c**, VTA oxLight baseline, as a. **d**, NAcc dLight reward response (last + running), as a. **e**, Per-trial correlations (r *×* 100) between the five Fig. 2 measures on engaged (lower triangle) and disengaged trials (upper triangle), mean over mice; boxes, q < 0.05 (Benjamini–Hochberg over the 10 pairs of each state). **f**, Share of the high − low reward-block effect on the disengage hazard, the re-engage hazard and P(disengaged) that each signal accounts for (as Fig. 2d), with wheel movement and licking around the previous reward as additional covariates, every row fitted on all trials (Methods); dots, share; lines, 95% bootstrap interval over mice; stars, one-sample t-tests. **g**, Slope of each signal’s pre-stimulus level on the recent reward rate (time constant 10 trials); dots, mice; stars, against 0. **h**, Pairwise interactions between the five measures in one per-mouse logistic model of each hazard (Methods), low and high reward; dots, mean coefficient over mice (log-odds per z^2^); lines, 95% CI; black, Benjamini–Hochberg q < 0.05 over the ten pairs of each panel; dark gray, P < 0.05 uncorrected. The two surviving terms were negative in 8/9 and 9/9 mice. Vehicle sessions of both experiments; n = 9 mice (SNc 8; pupil 14). Statistics in Supplementary Table 1.

**Extended Data Fig. 3.**
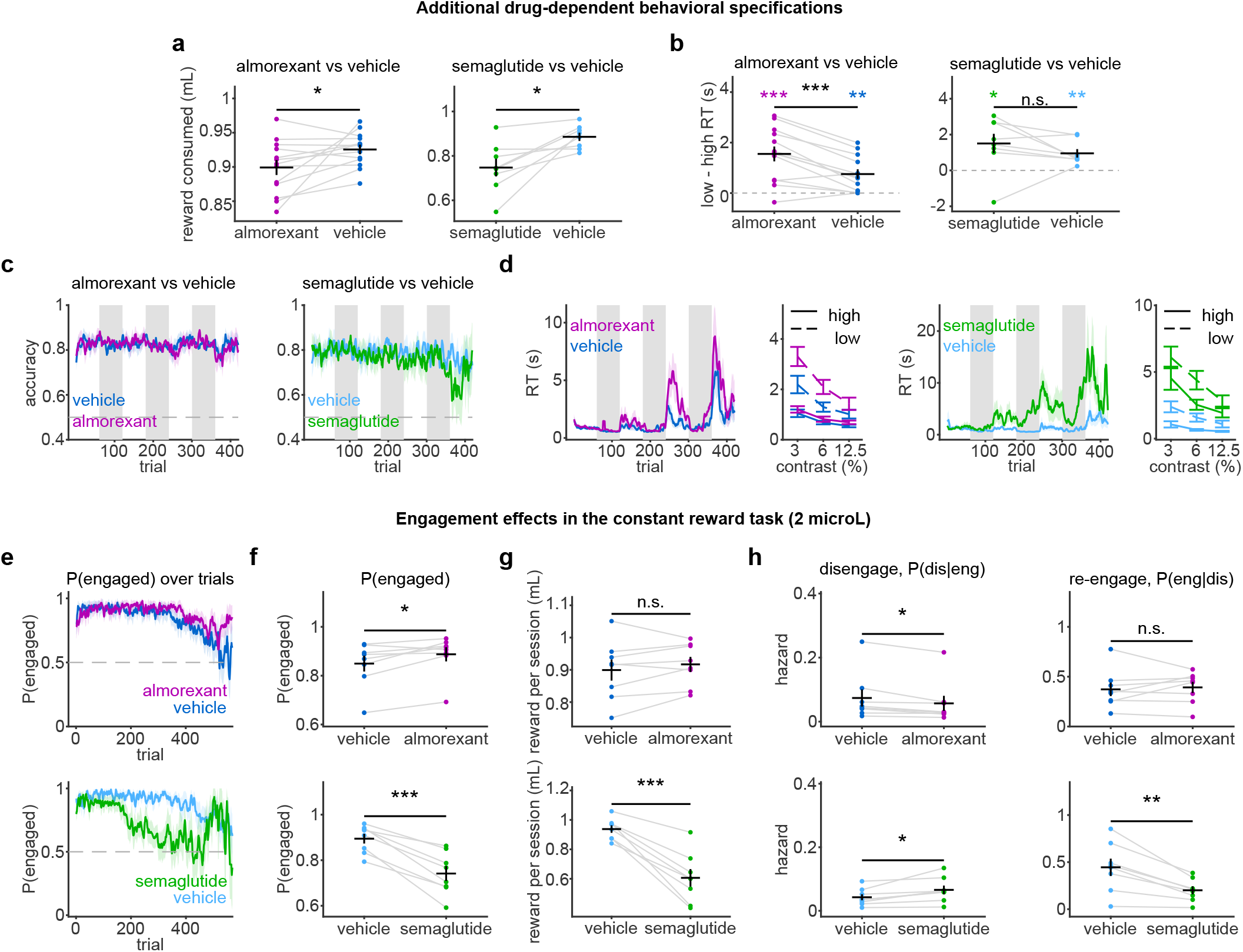
Drug effects on behavior; the constant-reward task. **a**, Reward consumed per session, drug vs control. **b**, Effect of reward size on reaction time (low − high reward) per condition (stars above dots, against 0) and drug vs control (bracket). **c**, Accuracy across the session (gray, high-reward blocks). **d**, Reaction time across the session (left) and against contrast (right; solid, high reward; dashed, low), drug vs control. **e**–**h**, Constant-reward task, the same mice and drugs with a constant 2-µL reward. Each session was cut after trial 570, where it had offered as much reward as a reward-task session (1,140 µL in 420 trials), and sessions with an accuracy below 0.65 were excluded, as in the reward task. **e**, P(engaged) across the session. **f**, P(engaged) per mouse. **g**, Reward consumed per session. **h**, Disengage and re-engage hazards. n = 13 mice (almorexant/vehicle) and 8 mice (semaglutide/vehicle) in a–d; 8 mice per pair in e–h. Statistics in Supplementary Table 1.

**Extended Data Fig. 4.**
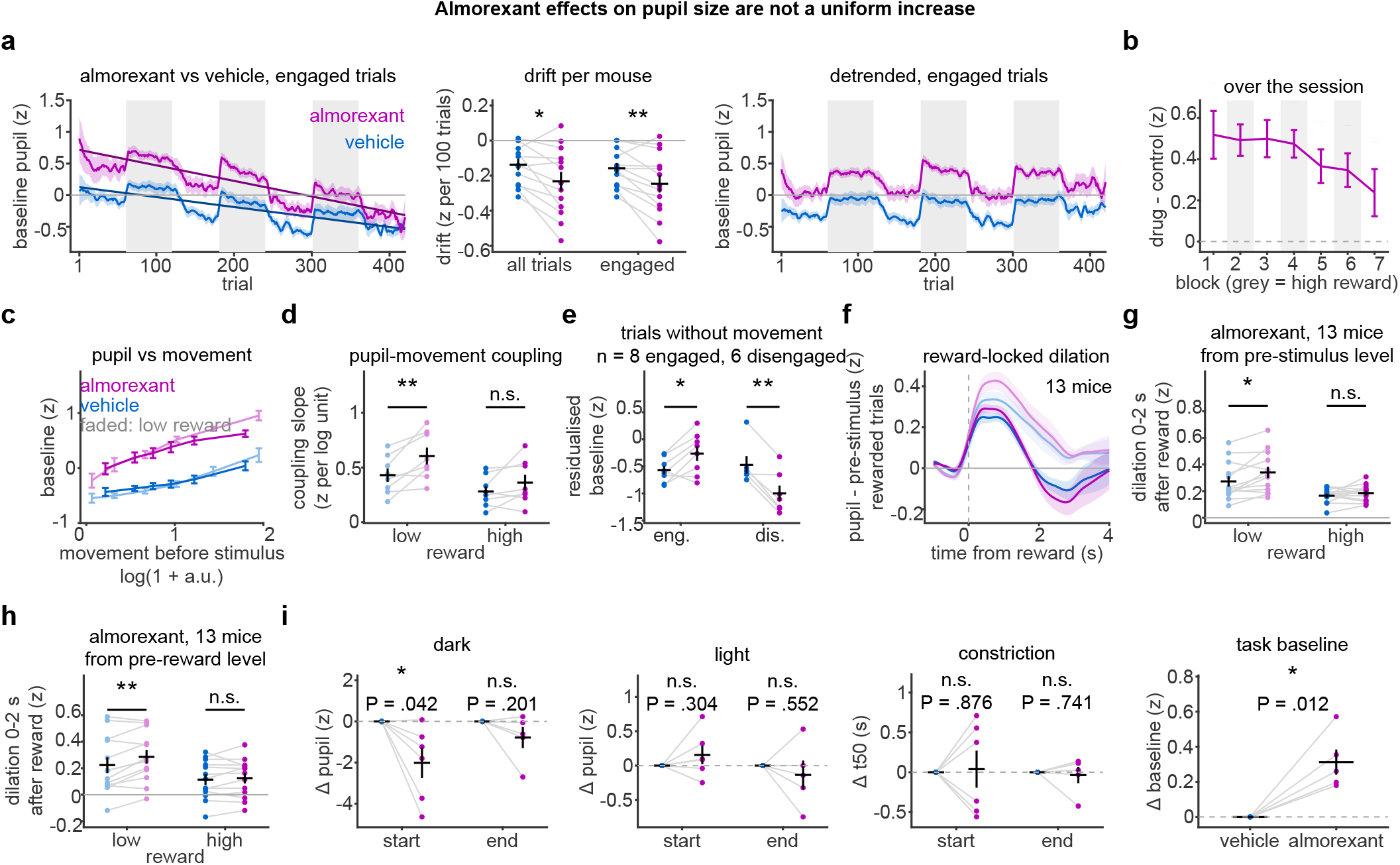
Almorexant effects on pupil size are not a uniform increase. **a**, Baseline pupil across the session on engaged trials with fitted lines (left), the per-mouse drift (middle; all and engaged trials) and the pupil after removing each session’s linear trend (right). **b**, Almorexant − vehicle baseline pupil per 60-trial block (gray, high reward). **c**, Residualized baseline pupil against wheel movement before the stimulus (six within-mouse quantile bins; faded, low reward). **d**, Slope of that relation per mouse, low and high reward. **e**, Baseline pupil on trials without wheel movement, engaged vs disengaged. **f**, Reward-locked pupil minus the prestimulus baseline on rewarded trials. **g, h**, Dilation 0–2 s after reward, from the pre-stimulus baseline (**g**) and from the pre-reward pupil (**h**). **i**, Light test in a second cohort (n = 6 mice; 5 at the end of the session and for the task baseline). Almorexant − vehicle change in the dark and light pupil and in the constriction t50 at the start and end of the session, and in the task baseline (z: each mouse centered on the mean of its task baselines, one SD pooled over mice; Methods). n = 13 mice (a, b, f–h); 8 mice with wheel recordings (c, d; e, 8 engaged and 6 disengaged). Statistics in Supplementary Table 1.

**Extended Data Fig. 5.**
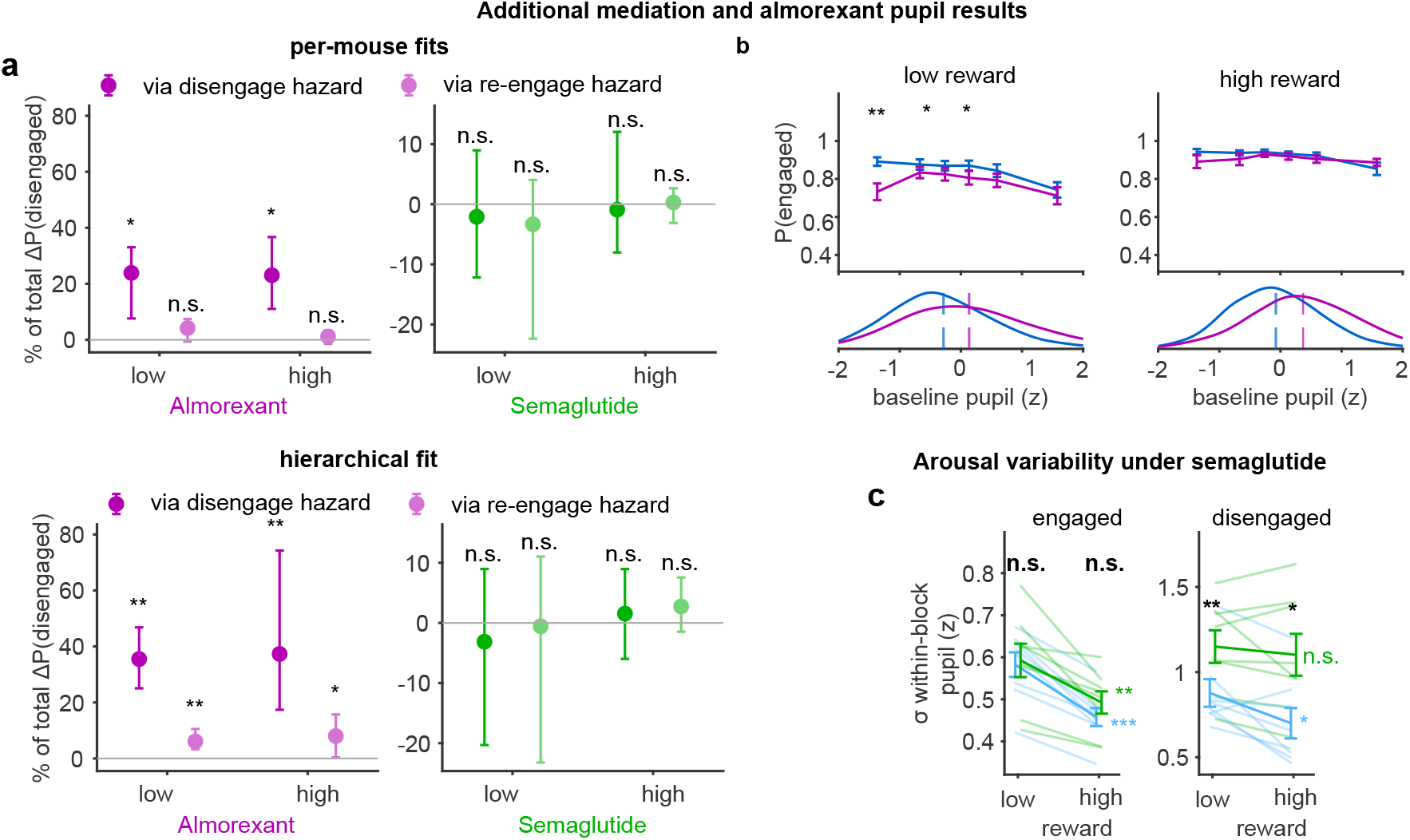
Additional arousal mediation results; arousal variability under semaglutide. **a**, Arousal routes from the generative model, the RT-HMM refitted with pupil-driven transitions (Methods), as the pure indirect effect of Fig. 4d. Per-mouse fits (top; Hodges–Lehmann estimate, exact Wilcoxon 95% CI, signed-rank test) and hierarchical fit (bottom; estimate and 95% cluster-bootstrap interval), almorexant and semaglutide, low and high reward. **b**, P(engaged) against baseline pupil in low (left) and high (right) reward blocks, vehicle vs almorexant; below, pupil distributions (ticks, means); stars, per-bin paired t-tests. **c**, Within-block SD of the baseline pupil (within-mouse z) in low and high reward blocks, engaged (left) and disengaged (right) trials, vehicle vs semaglutide; black stars, drug vs vehicle; colored stars, low vs high reward. n = 13 mice (almorexant/vehicle) and 8 mice (semaglutide/vehicle). Statistics in Supplementary Table 1.

**Extended Data Fig. 6.**
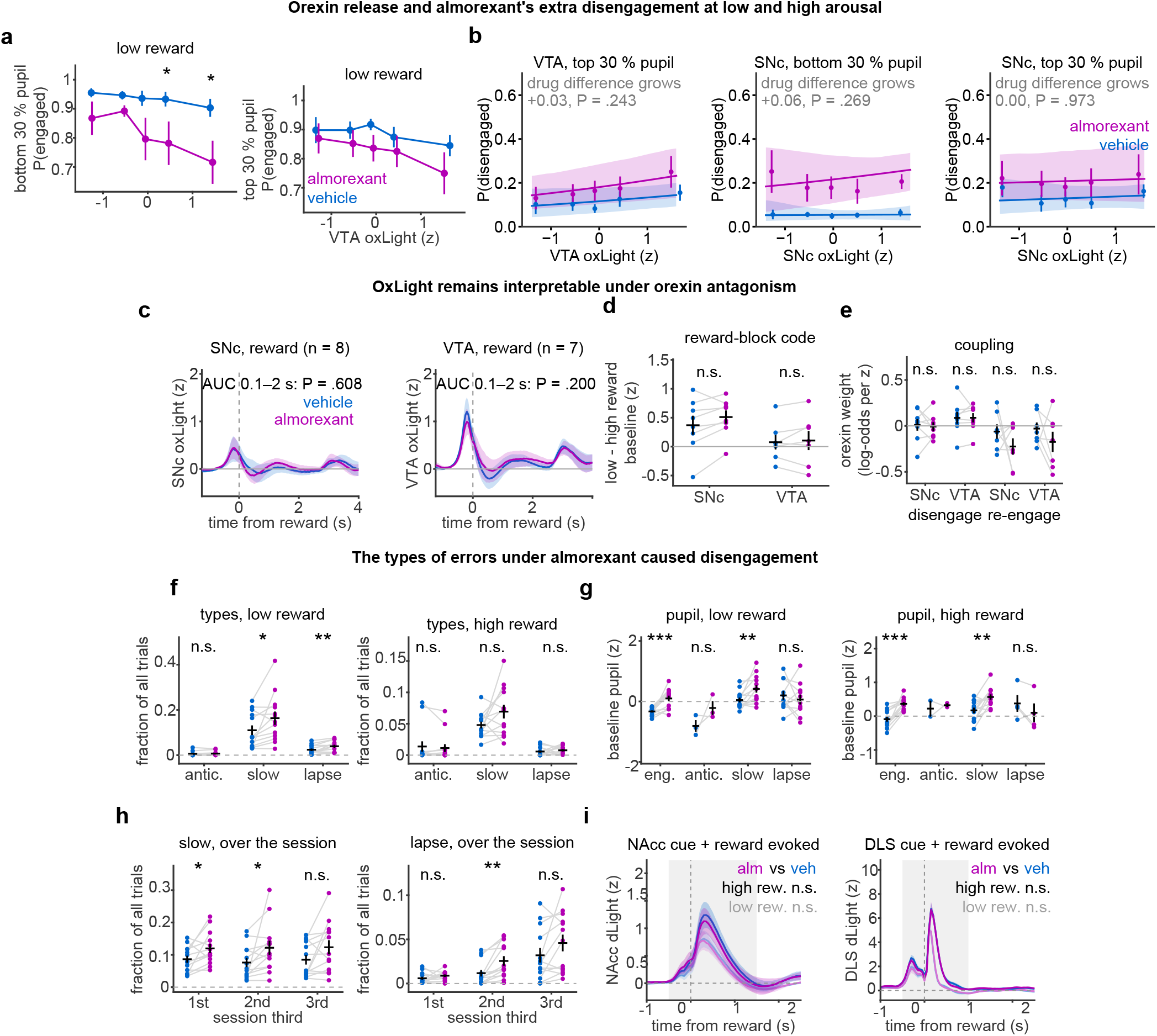
Orexin release at low and high arousal, oxLight under orexin antagonism, kinds of disengagement and striatal dopamine responses. **a**, P(engaged) against the VTA oxLight baseline within each mouse’s bottom (left) and top (right) 30% of baseline pupil, low reward, vehicle vs almorexant; stars, bins in which the drugs differ (paired t-tests, uncorrected). **b**, Model-based P(disengaged) along oxLight release (as Fig. 4e, right) for VTA at high pupil and for SNc at low and high pupil; text, growth of the drug difference from −1 to +1.5 z (bootstrap P). **c**, Reward-aligned SNc (n = 8) and VTA (n = 7) oxLight under vehicle and almorexant; text, AUC 0.1–2 s, paired t-test. **d**, Low − high reward-block code of the oxLight baseline. **e**, Weight of the oxLight baseline in predicting the two transitions (per-mouse logistic regression, log-odds per z). **f**, Disengaged trials split by reaction time (anticipatory < 0.3 s; slow 0.3–10 s; lapse > 10 s or no response), as fractions of all trials, low and high reward. **g**, Baseline pupil on engaged trials and on each type of disengaged trial. **h**, Fraction of slow and lapse trials in each third of the session. **i**, NAcc (left) and DLS (right) dLight aligned to reward, in z, in high (dark) and low (light) reward blocks, vehicle vs almorexant; gray, AUC window as Fig. 6f,g; text, almorexant vs vehicle on the AUC per mouse. n = 7 (VTA) and 8 (SNc) mice in a–e; 13 mice in f–h; 8 mice in i. Statistics in Supplementary Table 1.

**Extended Data Fig. 7.**
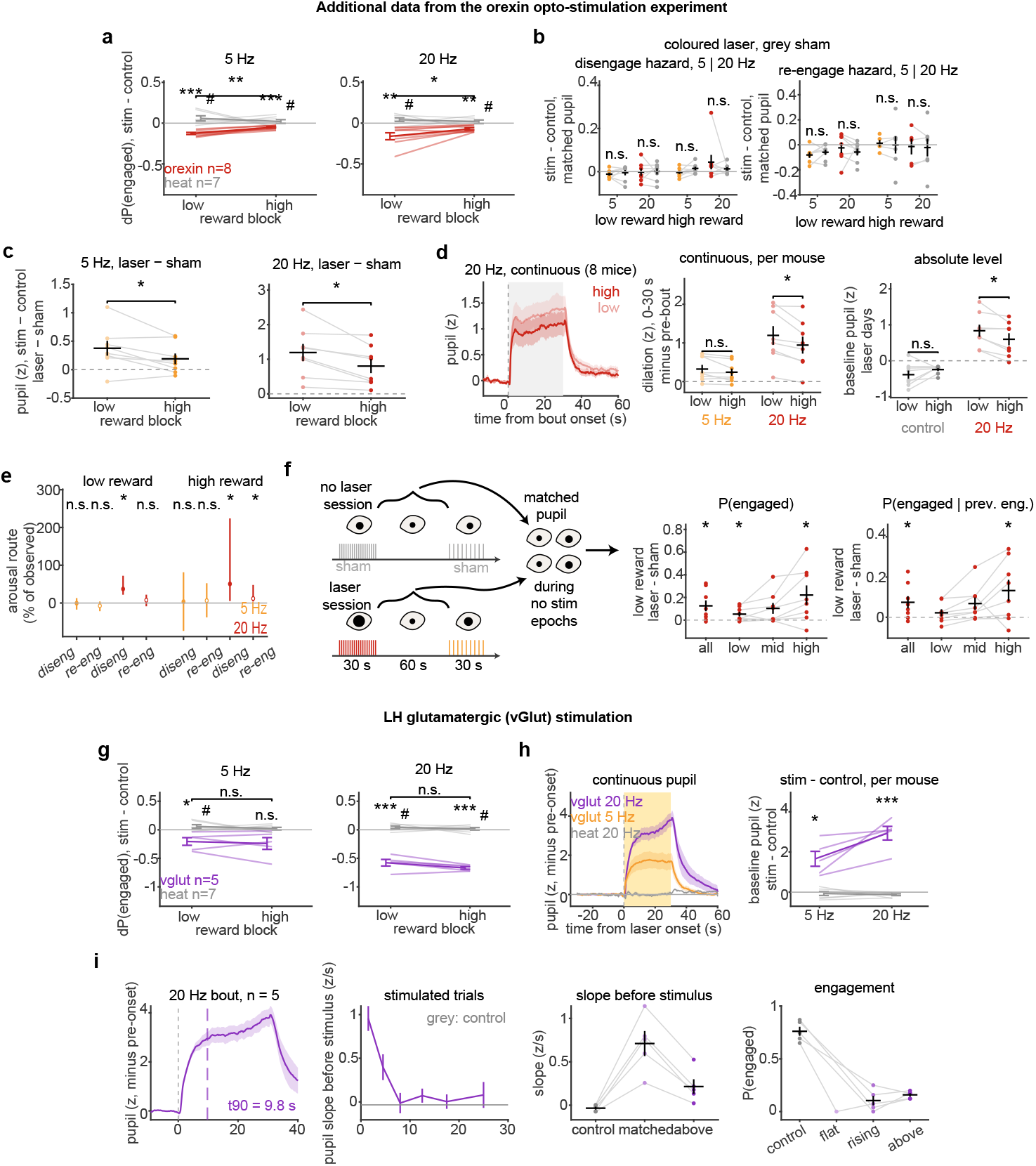
Orexin and vGlut stimulation: reward dependence, matched pupil, post-stimulation trials and arousal routes. **a**, Stimulated − control P(engaged) on laser days at 5 Hz (left) and 20 Hz (right), orexin vs heat controls (^*^ against 0; # vs heat; bracket, low vs high). **b**, Stimulated − control hazards at matched baseline pupil, laser (colored) vs sham (gray), 5 and 20 Hz, low and high reward. **c**, Stimulated − control baseline pupil, laser − sham, at 5 and 20 Hz in low and high reward blocks; bracket, low vs high (paired t-test). **d**, Dilation by reward block. Left, continuous pupil around 20-Hz bouts in high and low reward; middle, dilation 0–30 s after bout onset minus the pre-bout level, 5 and 20 Hz; right, baseline pupil of control and 20-Hz trials on laser days (z); brackets, low vs high reward (paired t-tests). **e**, Arousal routes through each hazard (stimulated pupil distribution under the control hazards, Methods), laser − sham, as % of the observed change in P(disengaged); 95% bootstrap interval over mice; stars, paired t-tests on the absolute route (8 mice in low, 7 in high reward). **f**, Post-stimulation trials. Left, design (laser-free trials 0–60 s after a bout, matched in baseline pupil to sham trials of the same mouse and reward block; Methods). Right, laser − sham P(engaged) in low reward over all matched trials, in thirds of the matched pupil range, and conditioned on an engaged previous trial; stars, against 0. **g**, vGlut stimulation. As a, for vGlut (n = 5) vs heat controls (n = 7). **h**, Continuous pupil aligned to laser onset (vGlut 5 and 20 Hz, heat 20 Hz) and stimulated − control baseline pupil per mouse. **i**, Pupil derivative control. Continuous pupil during a 20-Hz vGlut bout (t90 *≈* 10 s), pupil slope at trial onset by time into the bout (gray, control), and P(engaged) for control trials and for stimulated trials with a control-range or higher pupil slope. n = 8 orexin and 7 heat mice unless stated. Statistics in Supplementary Table 1.

**Extended Data Fig. 8.**
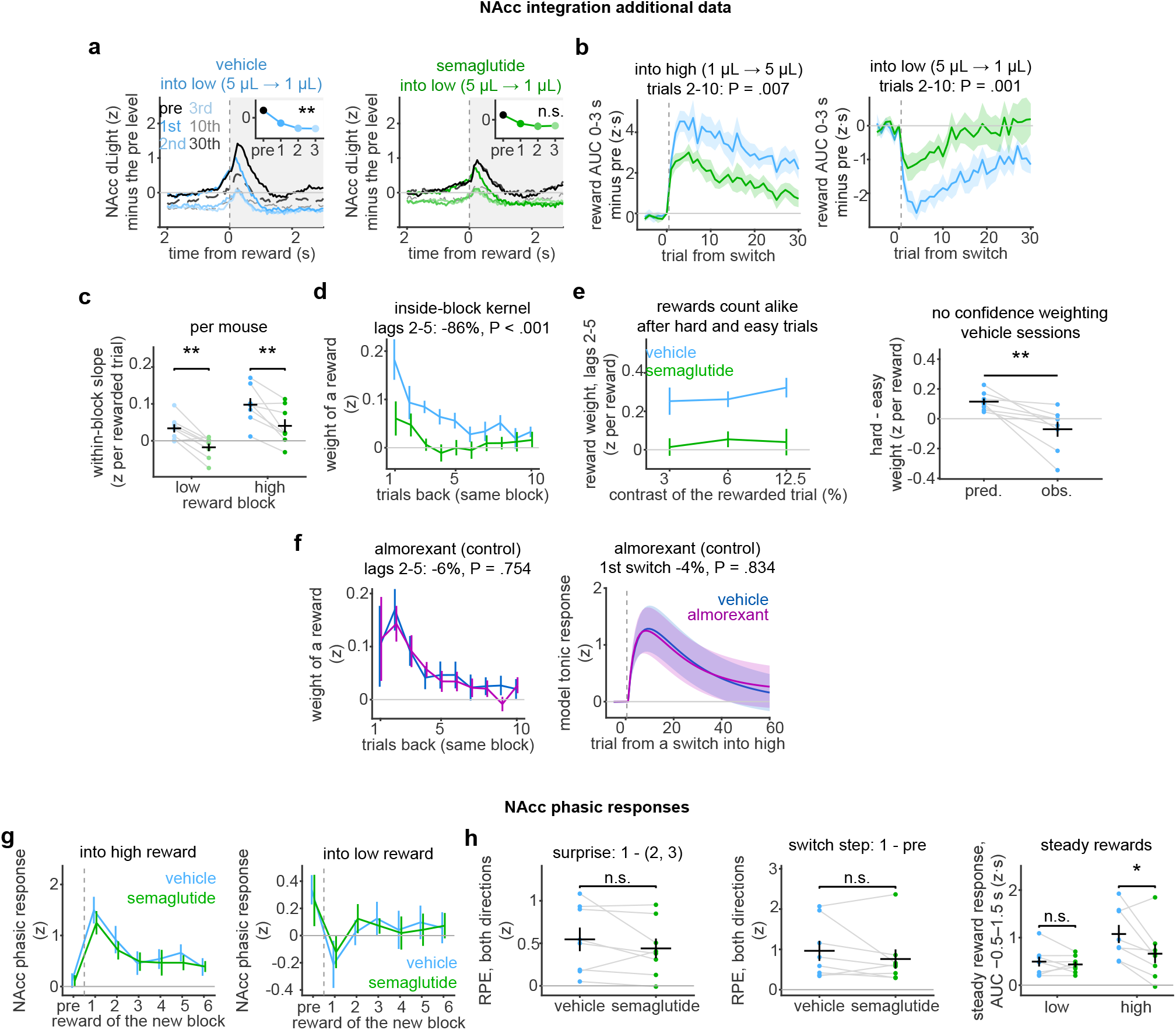
Reward integration by tonic NAcc dopamine under semaglutide. **a**, NAcc dLight at switches into low reward (5 *→* 1 µL), vehicle (left) and semaglutide (right), as Fig. 6a; insets, area 0–3 s after reward; star, the 2nd and 3rd trial vs the 1st. **b**, Reward AUC 0–3 s after reward (from the pre-switch tonic level) minus that of the pre-switch trial, around switches into high (left) and low (right) reward, vehicle vs semaglutide; above, test over trials 2–10. **c**, Integration within a block. Slope of the tonic NAcc level (minus its block mean) on the number of rewarded trials among the 2nd to 5th preceding trials (Methods), per mouse in low and high blocks. **d**, Weights of the rewards on the previous 10 trials of the same block on the tonic NAcc level, semaglutide vs vehicle; above, summed weight of lags 2–5. **e**, Reward weight (lags 2–5) after hard, mid and easy trials, vehicle vs semaglutide (left), and the hard − easy difference predicted by an integrator of confidence-weighted prediction errors against the observed one (right; vehicle sessions of the semaglutide experiment; paired t-test). **f**, Almorexant control: the reward-history kernel (left, as d) and the integrator response to the first switch into high reward with the same input for both drugs (right, as Fig. 6d), almorexant vs vehicle. **g**, NAcc phasic responses to the rewards of a new block after switches into high (left) and low (right) reward. **h**, The surprise (1st reward minus the mean of the 2nd and 3rd) and the switch step (1st reward minus pre), both directions combined, and the steady reward response (rewarded trials more than 10 into a block; AUC −0.5 to 1.5 s minus the pre-stimulus level, as Fig. 6f; low and high reward). n = 8 mice per pair. Statistics in Supplementary Table 1.

**Extended Data Fig. 9.**
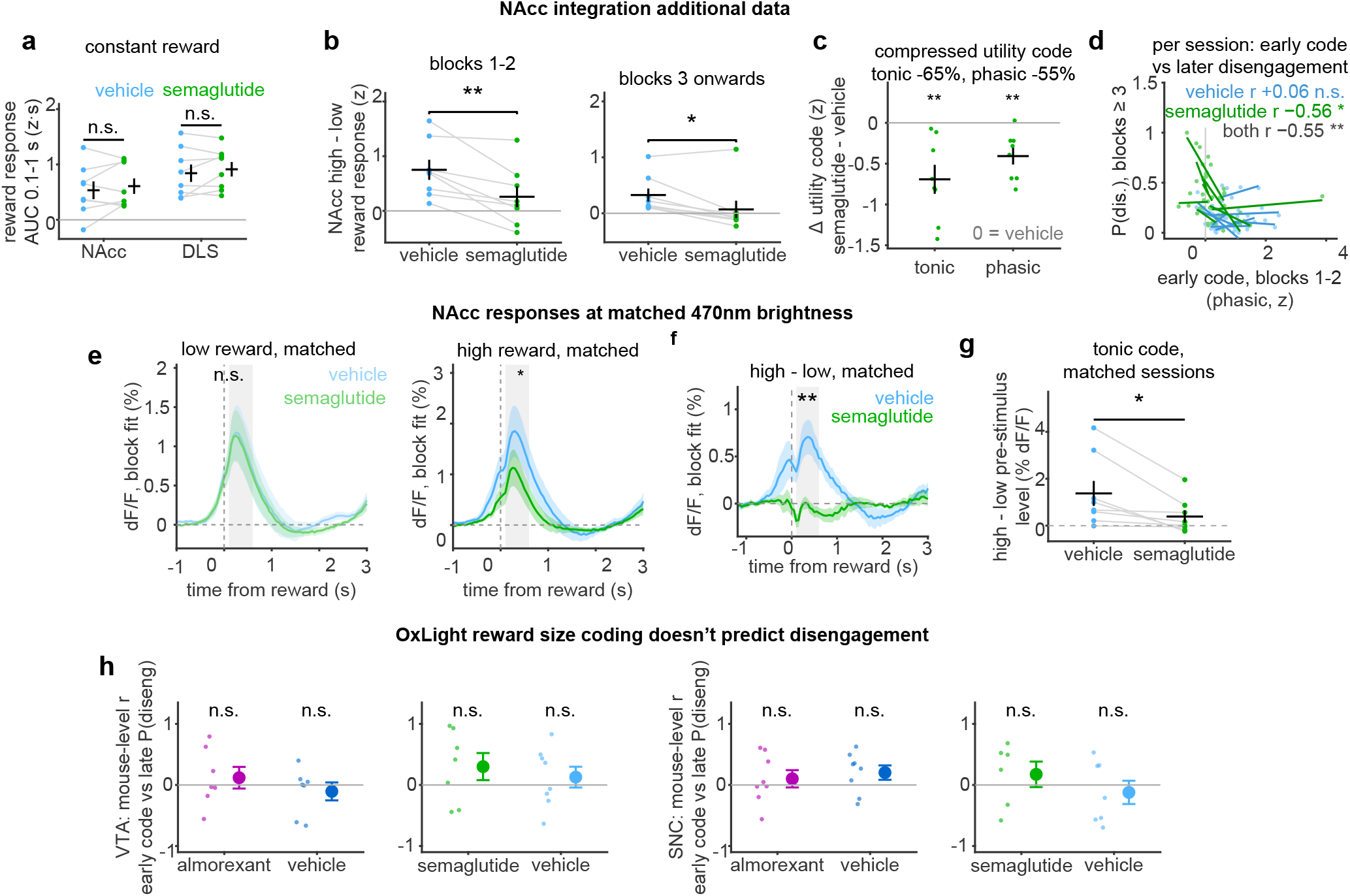
The NAcc value code between sessions, the constant-reward task and the dopamine changes under semaglutide. **a**, Constant-reward task. NAcc and DLS reward responses (AUC 0.1–1 s minus the pre-stimulus level; engaged rewarded trials), vehicle vs semaglutide per mouse. **b**, NAcc high − low reward response over blocks 1–2 and over blocks 3 onwards, vehicle vs semaglutide. **c**, Tonic and phasic utility codes (first high-reward block minus the flanking low-reward blocks, trials more than 10 into a block), semaglutide minus each mouse’s vehicle value. **d**, Per session, the early phasic utility code (block 2 minus block 1) against P(disengaged) from block 3 on; r, within mouse and drug or within mouse over both drugs. The code predicted later disengagement within semaglutide but not within vehicle sessions, so no vehicle-rule share is reported. **e**, NAcc reward responses on brightness-matched session pairs in low (left) and high (right) reward blocks (gray, 0.1–0.6 s; star, semaglutide vs vehicle). **f**, High − low reward-aligned NAcc response on the same pairs (gray, 0.1–0.6 s; star, paired t-test on that window). **g**, High − low pre-stimulus NAcc level on the same pairs (Methods). **h**, Mouse-level correlation between the early VTA (left) and SNc (right) oxLight code and late disengagement, almorexant and semaglutide experiments with their vehicles. e–g in ΔF/F, otherwise z. n = 8 mice per pair (oxLight 6–8 mice). Statistics in Supplementary Table 1.

**Extended Data Fig. 10.**
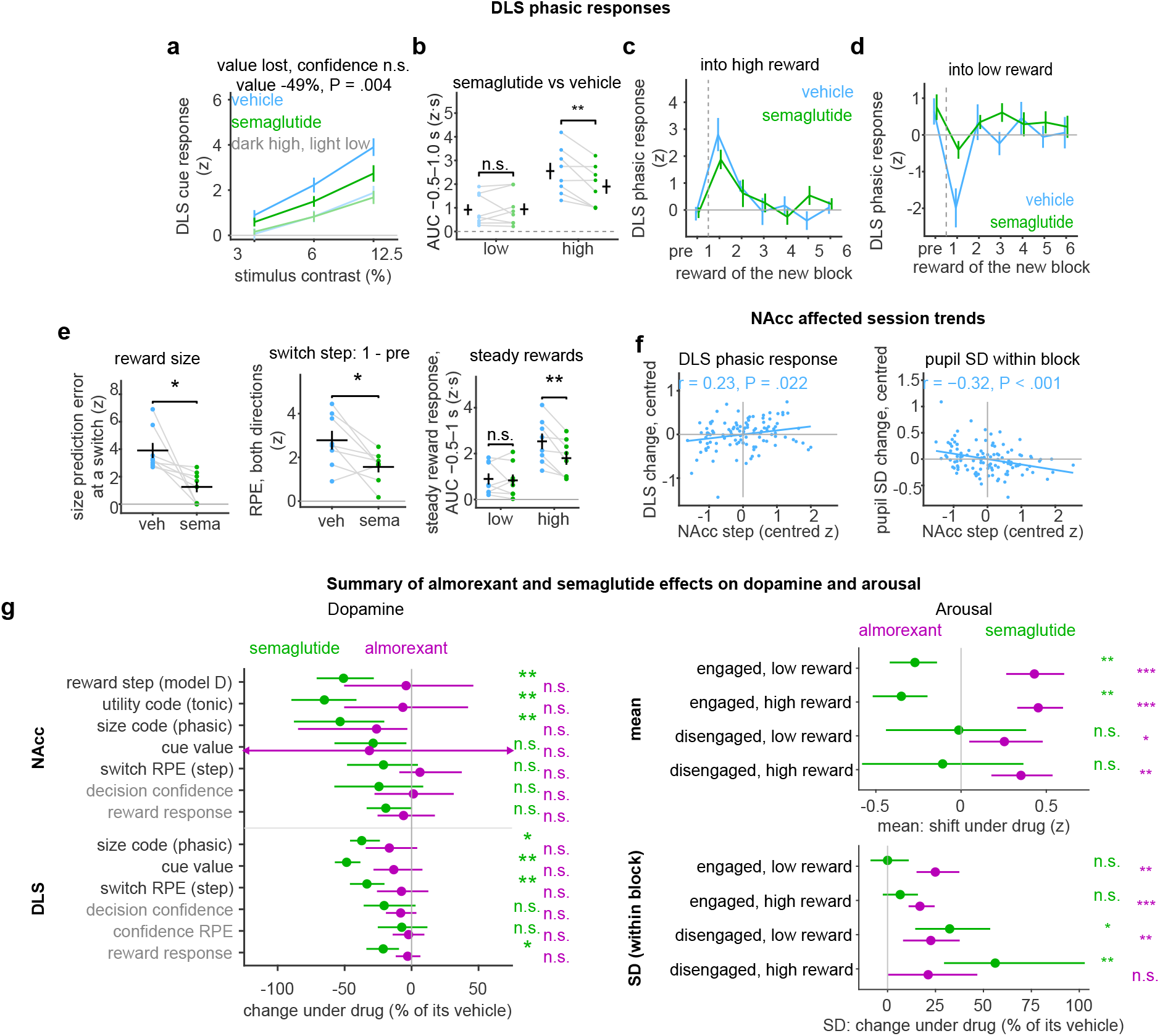
DLS dopamine under semaglutide. **a**, DLS cue response against stimulus contrast in high (dark) and low (light) reward blocks, vehicle vs semaglutide; text, change of the high − low cue value and of the confidence slope. **b**, DLS reward response (AUC −0.5 to 1.0 s, as Fig. 6g) per mouse in low and high reward blocks, vehicle vs semaglutide. The response fell only in high-reward blocks, whereas disengagement rose in both (Fig. 3c); its share of the extra disengagement is in Fig. 6g. **c, d**, DLS phasic response (corrected for stimulus contrast) to the last reward of the old block (pre) and the first six rewards after a switch into high (**c**) and low (**d**) reward. **e**, Left, reward-size prediction error at a switch from a model of the DLS reward response with a reward-size and a confidence prediction error (Methods); middle, the switch step (1st reward minus pre, both directions combined); right, the steady reward response within blocks (rewarded trials more than 10 into a block; AUC −0.5 to 1.0 s minus the pre-stimulus level, as Fig. 6g; low and high reward). **f**, Per vehicle session (both experiments; 111–113 sessions, 9 mice), the integrator’s step at the first switch into high reward (Fig. 6e) against the session trends of the phasic DLS reward response and the within-block pupil SD, centered within mouse (the trend of P(disengaged) is in Fig. 6e); r, pooled within-mouse correlation, P from a within-mouse permutation. Contrast with the trial-level DLS coupling, Fig. 2d–f. **g**, Summary. Left, change under semaglutide (green) and almorexant (magenta) of NAcc and DLS measures (% of vehicle; 95% bootstrap interval over mice; stars, paired t-tests; black labels, reward size or value; gray, confidence or mean reward response; definitions in Methods). Right, mean shift (z, top) and change of the within-block SD (% of vehicle, bottom) of the baseline pupil under each drug in each state and reward block. n = 8 mice per arm. Statistics in Supplementary Table 1.

## References

1. M. Husain and J.P. Roiser. Neuroscience of apathy and anhedonia: a transdiagnostic approach. Nat. Rev. Neurosci., 19:470–484, 2018. doi: 10.1038/s41583-018-0029-9.

2. N.D. Volkow, G.-J. Wang, J.H. Newcorn, S.H. Kollins, T.L. Wigal, F. Telang, J.S. Fowler, R.Z. Goldstein, N. Klein, J. Logan, et al. Motivation Deficit in ADHD is Associated with Dysfunction of the Dopamine Reward Pathway. Mol. Psychiatry, 16:1147–1154, 2011. doi: 10.1038/mp.2010.97.

3. Z.C. Ashwood, N.A. Roy, I.R. Stone, A.E. Urai, A.K. Churchland, A. Pouget, and J.W. Pillow. Mice alternate between discrete strategies during perceptual decision-making. Nat. Neurosci., 25:201–212, 2022. doi: 10.1038/s41593-021-01007-z.

4. S.A. Nuiten, M.N.O. Lohuis, A.C. Schaub, S. van Gaal, U. Olcese, C.M.A. Pennartz, P. Sterzer, and J.W. de Gee. Baseline activity of V1 interneurons connects pupil-linked arousal to engaged behavioral state. bioRxiv, 2026. doi: 10.64898/2026.02.05.704022.

5. J.W. de Gee, Z. Mridha, M. Hudson, Y. Shi, H. Ramsaywak, S. Smith, N. Karediya, M. Thompson, K. Jaspe, H. Jiang, et al. Strategic stabilization of arousal boosts sustained attention. Curr. Biol., 34:4114–4128.e6, 2024. doi: 10.1016/j.cub.2024.07.070.

6. P.A. Johnson, S. Nieuwenhuis, J. Mejías, and A.E. Urai. A dynamical systems model of arousal-driven behavioural state transitions. bioRxiv, 2025. doi: 10.1101/2025.10.31.685593.

7. D.A. Ganea, A. Bexter, M. Günther, P.M. Gardères, B.M. Kampa, and F. Haiss. Pupillary Dilations of Mice Performing a Vibrotactile Discrimination Task Reflect Task Engagement and Response Confidence. Front. Behav. Neurosci., 14, 2020. doi: 10.3389/fnbeh.2020.00159.

8. M.S. Gilzenrat, S. Nieuwenhuis, M. Jepma, and J.D. Cohen. Pupil diameter tracks changes in control state predicted by the adaptive gain theory of locus coeruleus function. Cogn. Affect. Behav. Neurosci., 10:252–269, 2010. doi: 10.3758/CABN.10.2.252.

9. G. Aston-Jones and J.D. Cohen. An integrative theory of locus coeruleus-norepinephrine function: adaptive gain and optimal performance. Annu Rev Neurosci, 28:403–450, 2005. doi: 10.1146/annurev.neuro.28.061604.135709.

10. R.M. Yerkes and J.D. Dodson. The Relation of Strength of Stimulus to Rapidity of Habit Formation. J. Comp. Neurol. Psychol., 18:459–482, 1908. doi: 10.1002/cne.920180503.

11. L. Beerendonk, J.F. Mejías, S.A. Nuiten, J.W. de Gee, J.B. Zantvoord, J.J. Fahrenfort, and S. van Gaal. Adaptive arousal regulation: Pharmacologically shifting the peak of the Yerkes– Dodson curve by catecholaminergic enhancement of arousal. Proc. Natl. Acad. Sci., 122: e2419733122, 2025. doi: 10.1073/pnas.2419733122.

12. F. Cazettes, D. Reato, J.P. Morais, A. Renart, and Z.F. Mainen. Phasic Activation of Dorsal Raphe Serotonergic Neurons Increases Pupil Size. Curr. Biol., 31:192–197.e4, 2021. doi: 10.1016/j.cub.2020.09.090.

13. J. Reimer, M.J. McGinley, Y. Liu, C. Rodenkirch, Q. Wang, D.A. McCormick, and A.S. Tolias. Pupil fluctuations track rapid changes in adrenergic and cholinergic activity in cortex. Nat. Commun., 7:13289, 2016. doi: 10.1038/ncomms13289.

14. M. Megemont, J. McBurney-Lin, and H. Yang. Pupil diameter is not an accurate real-time readout of locus coeruleus activity. eLife, 11:e70510, 2022. doi: 10.7554/eLife.70510.

15. M. Privitera, K.D. Ferrari, L.M. von Ziegler, O. Sturman, S.N. Duss, A. Floriou-Servou, P.-L. Germain, Y. Vermeiren, M.T. Wyss, P.P. De Deyn, et al. A complete pupillometry toolbox for real-time monitoring of locus coeruleus activity in rodents. Nat. Protoc., 15:2301–2320, 2020. doi: 10.1038/s41596-020-0324-6.

16. N. Grujic, A. Tesmer, E. Bracey, D. Peleg-Raibstein, and D. Burdakov. Control and coding of pupil size by hypothalamic orexin neurons. Nat. Neurosci., 26:1160–1164, 2023. doi: 10.1038/s41593-023-01365-w.

17. N. Grujic, R. Polania, and D. Burdakov. Neurobehavioral meaning of pupil size. Neuron, 112:3381–3395, 2024. doi: 10.1016/j.neuron.2024.05.029.

18. M. Maheu, T.H. Donner, and J.S. Wiegert. Serotonergic and Noradrenergic Interactions in Pupil-linked Arousal. bioRxiv, 2025. doi: 10.1101/2025.03.26.644382.

19. P. Viskaitis, A.L. Tesmer, Z. Liu, M.M. Karnani, M. Arnold, D. Donegan, E. Bracey, N. Grujic, T. Patriarchi, D. Peleg-Raibstein, et al. Orexin neurons track temporal features of blood glucose in behaving mice. Nat. Neurosci., 27:1299–1308, 2024. doi: 10.1038/s41593-024-01648-w.

20. A.L. Tesmer, P. Viskaitis, D. Donegan, E.F. Bracey, N. Grujic, T. Patriarchi, D. Peleg-Raibstein, and D. Burdakov. Orexin population activity precisely reflects net body movement across behavioral and metabolic states. eLife, 14, 2025. doi: 10.7554/eLife.103738.2.

21. A.L. Tesmer, X. Li, E. Bracey, C. Schmandt, R. Polania, D. Peleg-Raibstein, and D. Burdakov. Orexin neurons mediate temptation-resistant voluntary exercise. Nat. Neurosci., 27:1774–1782, 2024. doi: 10.1038/s41593-024-01696-2.

22. M.M. Karnani, J. Apergis-Schoute, A. Adamantidis, L.T. Jensen, L. de Lecea, L. Fugger, and D. Burdakov. Activation of Central Orexin/Hypocretin Neurons by Dietary Amino Acids. Neuron, 72:616–629, 2011. doi: 10.1016/j.neuron.2011.08.027.

23. M.M. Karnani, C. Schöne, E.F. Bracey, J.A. González, P. Viskaitis, H.-T. Li, A. Adamantidis, and D. Burdakov. Role of spontaneous and sensory orexin network dynamics in rapid locomotion initiation. Prog. Neurobiol., 187:101771, 2020. doi: 10.1016/j.pneurobio.2020.101771.

24. P. Viskaitis, M. Arnold, C. Garau, L.T. Jensen, L. Fugger, D. Peleg-Raibstein, and D. Burdakov. Ingested non-essential amino acids recruit brain orexin cells to suppress eating in mice. Curr. Biol., 32:1812–1821.e4, 2022. doi: 10.1016/j.cub.2022.02.067.

25. I.A. Azeez, F. Del Gallo, L. Cristino, and M. Bentivoglio. Daily Fluctuation of Orexin Neuron Activity and Wiring: The Challenge of “Chronoconnectivity.” Front. Pharmacol., 9:1061, 2018. doi: 10.3389/fphar.2018.01061.

26. G. Aston-Jones, R.J. Smith, D.E. Moorman, and K.A. Richardson. Role of lateral hypothalamic orexin neurons in reward processing and addiction. Neuropharmacology, 56:112–121, 2009. doi: 10.1016/j.neuropharm.2008.06.060.

27. M.D.C. Belle, A.T.L. Hughes, D.A. Bechtold, P. Cunningham, M. Pierucci, D. Burdakov, and H.D. Piggins. Acute Suppressive and Long-Term Phase Modulation Actions of Orexin on the Mammalian Circadian Clock. J. Neurosci., 34:3607–3621, 2014. doi: 10.1523/JNEUROSCI.3388-13.2014.

28. V. Linehan and M. Hirasawa. Short-term Fasting Induces Alternate Activation of Orexin and Melanin-concentrating Hormone Neurons in Rats. Neuroscience, 491:156–165, 2022. doi: 10.1016/j.neuroscience.2022.04.006.

29. E. Mignot, D. Mayleben, I. Fietze, D. Leger, G. Zammit, C.L.A. Bassetti, S. Pain, D.S. Kinter, and T. Roth. Safety and efficacy of daridorexant in patients with insomnia disorder: results from two multicentre, randomised, double-blind, placebo-controlled, phase 3 trials. Lancet Neurol., 21:125–139, 2022. doi: 10.1016/S1474-4422(21)00436-1.

30. A. Mohebi, J.R. Pettibone, A.A. Hamid, J.-M.T. Wong, L.T. Vinson, T. Patriarchi, L. Tian, R.T. Kennedy, and J.D. Berke. Dissociable dopamine dynamics for learning and motivation. Nature, 570:65–70, 2019. doi: 10.1038/s41586-019-1235-y.

31. Y. Niv, N.D. Daw, D. Joel, and P. Dayan. Tonic dopamine: opportunity costs and the control of response vigor. Psychopharmacology (Berl.), 191:507–520, 2007. doi: 10.1007/s00213-006-0502-4.

32. A.A. Hamid, J.R. Pettibone, O.S. Mabrouk, V.L. Hetrick, R. Schmidt, C.M. Vander Weele, R.T. Kennedy, B.J. Aragona, and J.D. Berke. Mesolimbic dopamine signals the value of work. Nat. Neurosci., 19:117–126, 2016. doi: 10.1038/nn.4173.

33. L.E. Kingsbury, H. May, and N. Uchida. Dopaminergic reward signals shape foraging decisions through learned value expectations. bioRxiv, 2026. doi: 10.64898/2026.10.21.760765.

34. M. Fritsche, C. Toschi, Y. Shevchuk, O. Didenko, R. Puliyadi, A. Majumdar, R. Bogacz, and A. Lak. Dorsal striatal dopamine integrates sensory and reward prediction errors to guide perceptual decisions. bioRxiv, 2026. doi: 10.64898/2026.01.01.696999.

35. M.W. Howe and D.A. Dombeck. Rapid signalling in distinct dopaminergic axons during locomotion and reward. Nature, 535:505–510, 2016. doi: 10.1038/nature18942.

36. J.E. Markowitz, W.F. Gillis, M. Jay, J. Wood, R.W. Harris, R. Cieszkowski, R. Scott, D. Brann, D. Koveal, T. Kula, et al. Spontaneous behaviour is structured by reinforcement without explicit reward. Nature, 614:108–117, 2023. doi: 10.1038/s41586-022-05611-2.

37. H. Liu, R. Melani, M. Maltese, J. Taniguchi, A. Sankaramanchi, R. Zeng, J.R. Martin, and N.X. Tritsch. Subsecond dopamine fluctuations do not specify the vigor of ongoing actions. Nat. Neurosci., 28:2432–2438, 2025. doi: 10.1038/s41593-025-02102-1.

38. X. Cai, C. Liu, I. Tsutsui-Kimura, J.-H. Lee, C. Guo, A. Banerjee, J. Lee, R. Amo, Y. Xie, T. Patriarchi, et al. Dopamine dynamics are dispensable for movement but promote reward responses. Nature, 635:406–414, 2024. doi: 10.1038/s41586-024-08038-z.

39. A. Mohebi, W. Wei, L. Pelattini, K. Kim, and J.D. Berke. Dopamine transients follow a striatal gradient of reward time horizons. Nat. Neurosci., 27:737–746, 2024. doi: 10.1038/s41593-023-01566-3.

40. G.C. Harris and G. Aston-Jones. Arousal and reward: a dichotomy in orexin function. Trends Neurosci., 29:571–577, 2006. doi: 10.1016/j.tins.2006.08.002.

41. C.B. Pantazis, M.H. James, S. O’Connor, N. Shin, and G. Aston-Jones. Orexin-1 receptor signaling in ventral tegmental area mediates cue-driven demand for cocaine. Neuropsychopharmacology, 47:741–751, 2022. doi: 10.1038/s41386-021-01173-5.

42. D.E. Moorman and G. Aston-Jones. Orexin/Hypocretin Modulates Response of Ventral Tegmental Dopamine Neurons to Prefrontal Activation: Diurnal Influences. J. Neurosci., 30: 15585–15599, 2010. doi: 10.1523/JNEUROSCI.2871-10.2010.

43. G.A. Christou, N. Katsiki, J. Blundell, G. Fruhbeck, and D.N. Kiortsis. Semaglutide as a promising antiobesity drug. Obes. Rev., 20:805–815, 2019. doi: 10.1111/obr.12839.

44. Z. Zhu, R. Gong, V. Rodriguez, K.T. Quach, X. Chen, and S.M. Sternson. Hedonic eating is controlled by dopamine neurons that oppose GLP-1R satiety. Science, 387:eadt0773, 2025. doi: 10.1126/science.adt0773.

45. K.L. Kooij, D.Ij. Koster, E. Eeltink, M. Luijendijk, L. Drost, F. Ducrocq, and R.A.H. Adan. GLP-1 receptor agonist semaglutide reduces appetite while increasing dopamine reward signaling. Neurosci. Appl., 3:103925, 2024. doi: 10.1016/j.nsa.2023.103925.

46. E.P. Foscue, J.R. Trinko, J.L. Jiménez, E. Kong, S.L. Thompson, K. Stankewich, A.M. Corstens, M.J. Serlie, J.R. Taylor, and R.J. DiLeone. The GLP-1R Agonist Semaglutide Reduces Motivated Running and Alters Dopamine Dynamics in the Nucleus Accumbens. bioRxiv, 2025. doi: 10.1101/2025.10.08.681212.

47. C. Aranäs, A. Caffrey, C.E. Edvardsson, H.D. Schmidt, and E. Jerlhag. Semaglutide suppresses cocaine taking, seeking, and cocaine-evoked dopamine levels in the nucleus accumbens. Eur. Neuropsychopharmacol., 98:1–10, 2025. doi: 10.1016/j.euroneuro.2025.07.001.

48. C. Aranäs, C.E. Edvardsson, O.T. Shevchouk, Q. Zhang, S. Witley, S.B. Sköldheden, L. Zentveld, D. Vallöf, M. Tufvesson-Alm, and E. Jerlhag. Semaglutide reduces alcohol intake and relapse-like drinking in male and female rats. eBioMedicine, 93, 2023. doi: 10.1016/j.ebiom.2023.104642.

49. S.M. Fortin, J.C. Chen, M.C. Petticord, F.J. Ragozzino, J.H. Peters, and M.R. Hayes. The locus coeruleus contributes to the anorectic, nausea, and autonomic physiological effects of glucagon-like peptide-1. Sci. Adv., 9:eadh0980, 2023. doi: 10.1126/sciadv.adh0980.

50. C. Acuna-Goycolea and A. van den Pol. Glucagon-Like Peptide 1 Excites Hypocretin/Orexin Neurons by Direct and Indirect Mechanisms: Implications for Viscera-Mediated Arousal. J. Neurosci., 24:8141–8152, 2004. doi: 10.1523/JNEUROSCI.1607-04.2004.

51. C.S. Hendershot, M.P. Bremmer, M.B. Paladino, G. Kostantinis, T.A. Gilmore, N.R. Sullivan, A.C. Tow, S.S. Dermody, M.A. Prince, R. Jordan, et al. Once-Weekly Semaglutide in Adults With Alcohol Use Disorder: A Randomized Clinical Trial. JAMA Psychiatry, 82:395, 2025. doi: 10.1001/jamapsychiatry.2024.4789.

52. C.S. Hendershot, M.P. Bremmer, M.B. Paladino, G. Kostantinis, T.A. Gilmore, N.R. Sullivan, A.C. Tow, M.A. Prince, S.S. Dermody, R. Jordan, et al. Once-Weekly Semaglutide in Adults With Daily Cigarette Use: A Randomized Clinical Trial. JAMA Netw. Open, 9:e2614898, 2026. doi: 10.1001/jamanetworkopen.2026.14898.

53. V. Aguillon-Rodriguez, D. Angelaki, H. Bayer, N. Bonacchi, M. Carandini, F. Cazettes, G. Chapuis, A.K. Churchland, Y. Dan, E. Dewitt, et al. Standardized and reproducible measurement of decision-making in mice. eLife, 10, 2021. doi: 10.7554/ELIFE.63711.

54. M.R. Roesch, D.J. Calu, and G. Schoenbaum. Dopamine neurons encode the better option in rats deciding between differently delayed or sized rewards. Nat. Neurosci., 10:1615–1624, 2007. doi: 10.1038/nn2013.

55. P.N. Tobler, C.D. Fiorillo, and W. Schultz. Adaptive coding of reward value by dopamine neurons. Science, 307:1642–1645, 2005. doi: 10.1126/science.1105370.

56. M.P. Saddoris, F. Cacciapaglia, R.M. Wightman, and R.M. Carelli. Differential Dopamine Release Dynamics in the Nucleus Accumbens Core and Shell Reveal Complementary Signals for Error Prediction and Incentive Motivation. J. Neurosci., 35:11572–11582, 2015. doi:10.1523/JNEUROSCI.2344-15.2015.

57. A. Lak, M. Okun, M.M. Moss, H. Gurnani, K. Farrell, M.J. Wells, C.B. Reddy, A. Kepecs, K.D. Harris, and M. Carandini. Dopaminergic and Prefrontal Basis of Learning from Sensory Confidence and Reward Value. Neuron, 105:700–711.e6, 2020. doi: 10.1016/j.neuron.2019.11.018.

58. C. Anaclet, R. Parmentier, K. Ouk, G. Guidon, C. Buda, J.-P. Sastre, H. Akaoka, O.A. Sergeeva, M. Yanagisawa, H. Ohtsu, et al. Orexin/Hypocretin and Histamine: Distinct Roles in the Control of Wakefulness Demonstrated Using Knock-Out Mouse Models. J. Neurosci., 29:14423–14438, 2009. doi: 10.1523/JNEUROSCI.2604-09.2009.

59. J. Hara, C.T. Beuckmann, T. Nambu, J.T. Willie, R.M. Chemelli, C.M. Sinton, F. Sugiyama, K. Yagami, K. Goto, and M. Yanagisawa. Genetic ablation of orexin neurons in mice results in narcolepsy, hypophagia, and obesity. Neuron, 30:345–354, 2001. doi: 10.1016/s0896-6273(01)00293-8.

60. T.M. Korotkova, O.A. Sergeeva, K.S. Eriksson, H.L. Haas, and R.E. Brown. Excitation of Ventral Tegmental Area Dopaminergic and Nondopaminergic Neurons by Orexins/Hypocretins. J. Neurosci., 23:7–11, 2003. doi: 10.1523/JNEUROSCI.23-01-00007.2003.

61. S.L. Borgland, S.A. Taha, F. Sarti, H.L. Fields, and A. Bonci. Orexin A in the VTA Is Critical for the Induction of Synaptic Plasticity and Behavioral Sensitization to Cocaine. Neuron, 49: 589–601, 2006. doi: 10.1016/j.neuron.2006.01.016.

62. M. Narita, Y. Nagumo, S. Hashimoto, M. Narita, J. Khotib, M. Miyatake, T. Sakurai, M. Yanagisawa, T. Nakamachi, and S. Shioda. Direct involvement of orexinergic systems in the activation of the mesolimbic dopamine pathway and related behaviors induced by morphine. J. Neurosci., 26:398–405, 2006.

63. D.L. Bernstein, P.S. Badve, J.R. Barson, C.E. Bass, and R.A. España. Hypocretin receptor 1 knockdown in the ventral tegmental area attenuates mesolimbic dopamine signaling and reduces motivation for cocaine. Addict. Biol., 23:1032–1045, 2018. doi: 10.1111/adb.12553.

64. C. Schöne, J. Apergis-Schoute, T. Sakurai, A. Adamantidis, and D. Burdakov. Coreleased orexin and glutamate evoke nonredundant spike outputs and computations in histamine neurons. Cell Rep., 7:697–704, 2014. doi: 10.1016/J.CELREP.2014.03.055.

65. R. Wang, H. Guo, S. Jiang, Z. Liu, W. Qu, Z. Huang, and L. Wang. Control of wakefulness by lateral hypothalamic glutamatergic neurons in male mice. J. Neurosci. Res., 99:1689–1703, 2021. doi: 10.1002/jnr.24828.

66. J.D. Salamone and M. Correa. The Mysterious Motivational Functions of Mesolimbic Dopamine. Neuron, 76:470–485, 2012. doi: 10.1016/j.neuron.2012.10.021.

67. W. Zhou, L.-Q. Wang, Y.-Q. Shao, X. Han, C.-X. Yu, F. Yuan, X. Wang, S.-J. Weng, Y.-M. Zhong, and X.-L. Yang. Orexin-A intensifies mouse pupillary light response by modulating intrinsically photosensitive retinal ganglion cells. J. Neurosci., 41:2566–2580, 2021. doi: 10.1523/jneurosci.0217-20.2021.

68. R.E. Yoss, N.J. Moyer, and K.N. Ogle. The pupillogram and narcolepsy: A method to measure decreased levels of wakefulness. Neurology, 19:921–921, 1969. doi: 10.1212/WNL.19.10.921.

69. J. Newman and R. Broughton. Pupillometric Assessment of Excessive Daytime Sleepiness in Narcolepsy-Cataplexy. Sleep, 14:121–129, 1991. doi: 10.1093/sleep/14.2.121.

70. W.D. O’Neill, A.M. Oroujeh, and S.L. Merritt. Pupil noise is a discriminator between narcoleptics and controls. IEEE Trans. Biomed. Eng., 45:314–322, 1998. doi: 10.1109/10.661156.

71. S. Schreyer, J.A. Büttner-Ennever, X. Tang, M.J. Mustari, and A.K.E. Horn. Orexin-A inputs onto visuomotor cell groups in the monkey brainstem. Neuroscience, 164:629–640, 2009. doi: 10.1016/j.neuroscience.2009.08.039.

72. L. Chrobok, J.S. Jeczmien-Lazur, K. Pradel, J.D. Klich, M. Bubka, M. Wojcik, M. Kepczynski, and M.H. Lewandowski. Circadian actions of orexins on the retinorecipient lateral geniculate complex in rat. J. Physiol., 599:231–252, 2021.

73. H.G. Cruz, J.L. Hay, P. Hoever, F. Alessi, E.T. Te Beek, J.M.A. Van Gerven, and J. Dinge-manse. Pharmacokinetic and pharmacodynamic interactions between almorexant, a dual orexin receptor antagonist, and desipramine. Eur. Neuropsychopharmacol., 24:1257–1268, 2014. doi: 10.1016/j.euroneuro.2014.05.002.

74. J.R. Floeder, H. Jeong, A. Mohebi, and V.M.K. Namboodiri. Mesolimbic dopamine ramps reflect environmental timescales. eLife, 13:RP98666, 2025. doi: 10.7554/eLife.98666.

75. E.N. Godschall, T.B. Gungul, I.R. Sajonia, A.K. Buyukaksakal, O. Li, S. Ogilvie, A.B. Keeler, G. Tian, Y. Shi, O. Koita, et al. A brain reward circuit inhibited by next-generation weight-loss drugs in mice. Nature, 654:1055–1064, 2026. doi: 10.1038/s41586-026-10444-4.

76. V.R. Konanur, T.M. Hsu, S.E. Kanoski, M.R. Hayes, and M.F. Roitman. Phasic dopamine responses to a food-predictive cue are suppressed by the glucagon-like peptide-1 receptor agonist Exendin-4. Physiol. Behav., 215:112771, 2020. doi: 10.1016/j.physbeh.2019.112771.

77. N.S. Hernandez, B. O’Donovan, P.I. Ortinski, and H.D. Schmidt. Activation of glucagon-like peptide-1 receptors in the nucleus accumbens attenuates cocaine seeking in rats. Addict. Biol., 24:170–181, 2019. doi: 10.1111/adb.12583.

78. K.C. Berridge and T.E. Robinson. What is the role of dopamine in reward: hedonic impact, reward learning, or incentive salience? Brain Res. Rev., 28:309–369, 1998. doi: 10.1016/S0165-0173(98)00019-8.

79. K.C. Berridge. The debate over dopamine’s role in reward: the case for incentive salience. Psychopharmacology (Berl.), 191:391–431, 2007. doi: 10.1007/s00213-006-0578-x.

80. N. Eshel, G.C. Touponse, A.R. Wang, A.K. Osterman, A.N. Shank, A.M. Groome, L. Taniguchi, D.F. Cardozo Pinto, J. Tucciarone, B.S. Bentzley, et al. Striatal dopamine integrates cost, benefit and motivation. Neuron, 112:500–514.e5, 2024. doi: 10.1016/j.neuron.2023.10.038.

81. J.W. de Gee, O. Colizoli, N.A. Kloosterman, T. Knapen, S. Nieuwenhuis, and T.H. Donner. Dynamic modulation of decision biases by brainstem arousal systems. eLife, 6:e23232, 2017. doi: 10.7554/eLife.23232.

82. A.E. Urai, V. Aguillon-Rodriguez, I.C. Laranjeira, F. Cazettes, Z.F. Mainen, and A.K. Churchland. Citric acid water as an alternative to water restriction for high-yield mouse behavior. eNeuro, 8:1–8, 2021. doi: 10.1523/ENEURO.0230-20.2020.

83. J.A. González, L.T. Jensen, P. Iordanidou, M. Strom, L. Fugger, and D. Burdakov. Inhibitory Interplay between Orexin Neurons and Eating. Curr. Biol., 26:2486–2491, 2016. doi: 10.1016/J.CUB.2016.07.013.

84. T. Nath, A. Mathis, A.C. Chen, A. Patel, M. Bethge, and M.W. Mathis. Using DeepLabCut for 3D markerless pose estimation across species and behaviors. Nat. Protoc., 14:2152–2176, 2019. doi: 10.1038/s41596-019-0176-0.

