## Supplementary Table 1 for "Rules of disengagement: orexin–arousal and dopamine–utility as separable axes"

### Supplementary Table 1 | Statistics of the main and Extended Data figures

Every statistical test shown on a figure panel. All tests are two-sided. n is the number of mice unless stated; for paired comparisons each mouse contributes one value per condition. t, t statistic with degrees of freedom; r, correlation coefficient; q, Benjamini–Hochberg (BH) adjusted P. —, statistic not reported by the analysis code (P from the stated test); †, t computed as mean / s.e.m. across mice. Significance marks on the figures: \* P < 0.05, \*\* P < 0.01, \*\*\* P < 0.001.

| Panel | Comparison | Test | n | Statistic | P |
| --- | --- | --- | --- | --- | --- |
| **Fig. 1** |  |  |  |  |  |
| e | engaged state: low vs high reward | Paired t-test | 14 mice | t(13) = 2.28 | .040 |
| e | disengaged state: low vs high reward | Paired t-test | 12 mice | t(11) = -0.28 | .79 |
| f (left) | disengage hazard P(dis\|eng): low vs high reward | Paired t-test | 14 mice | t(13) = 5.32 | $1.4 \times 10^{-4}$ |
| f (right) | re-engage hazard P(eng\|dis): low vs high reward | Paired t-test | 14 mice | t(13) = -5.81 | $6.1 \times 10^{-5}$ |
| g (right) | P(engaged): high vs low reward | Paired t-test | 14 mice | t(13) = 5.83 | $5.9 \times 10^{-5}$ |
| **Fig. 2** |  |  |  |  |  |
| a | high vs low reward block | Paired t-test | 14 mice | t(13) = 2.74 | .017 |
| a | high vs low reward block | Paired t-test | 14 mice | t(13) = -19.75 | $4.4 \times 10^{-11}$ |
| a | high-reward blocks: linear trend across the 6 bins | One-sample t-test | 14 mice | t(13) = -2.25 | .042 |
| a | low-reward blocks: linear trend across the 6 bins | One-sample t-test | 14 mice | t(13) = -5.39 | $1.2 \times 10^{-4}$ |
| a | low-reward blocks: curvature (quadratic term) across the 6 bins | One-sample t-test | 14 mice | t(13) = -4.15 | .0011 |
| a | high-reward blocks: linear trend across the 6 bins | One-sample t-test | 13 mice | t(12) = -2.57 | .025 |
| a | low-reward blocks: linear trend across the 6 bins | One-sample t-test | 14 mice | t(13) = -4.67 | $4.4 \times 10^{-4}$ |
| a | low-reward blocks: linear trend across the 6 bins | One-sample t-test | 14 mice | t(13) = 6.05 | $4.1 \times 10^{-5}$ |
| a | low-reward blocks: curvature (quadratic term) across the 6 bins | One-sample t-test | 14 mice | t(13) = 5.72 | $7.1 \times 10^{-5}$ |
| c | high vs low reward block | Paired t-test | 9 mice | t(8) = 24.06 | $9.5 \times 10^{-9}$ |
| c | high-reward blocks: linear trend across the 6 bins | One-sample t-test | 9 mice | t(8) = 2.33 | .049 |
| c | low-reward blocks: linear trend across the 6 bins | One-sample t-test | 9 mice | t(8) = 2.67 | .028 |
| c | high-reward blocks: linear trend across the 6 bins | One-sample t-test | 8 mice | t(7) = 2.44 | .045 |
| c | low-reward blocks: linear trend across the 6 bins | One-sample t-test | 9 mice | t(8) = 3.83 | .005 |
| c | low-reward blocks: linear trend across the 6 bins | One-sample t-test | 9 mice | t(8) = -2.59 | .032 |
| c | low-reward blocks: curvature (quadratic term) across the 6 bins | One-sample t-test | 9 mice | t(8) = 2.59 | .032 |
| d | share of the high - low block effect on within-block pupil spread carried by DLS (quantile replacement) | One-sample t-test; bootstrap over mice | 9 mice | — | .003 |
| d | share of the high - low block effect on P(disengaged) carried by DLS (quantile replacement) | One-sample t-test; bootstrap over mice | 9 mice | — | .045 |
| d | share of the high - low block effect carried by pupil | One-sample t-test | 14 mice | — | n.s. |
| d | share of the high - low block effect carried by SNc | One-sample t-test | 8 mice | — | n.s. |
| d | share of the high - low block effect carried by VTA | One-sample t-test | 9 mice | — | n.s. |
| d | share of the high - low block effect carried by NAcc | One-sample t-test | 9 mice | — | n.s. |
| e | DLS previous reward response (per SD of last + running) vs 0 | One-sample t-test | 8 mice (both DLS and NAcc) | t(7) = -3.27† | .014 |
| e | NAcc previous reward response (per SD of last + running) vs 0 | One-sample t-test | 8 mice (both DLS and NAcc) | t(7) = -1.44† | .192 |
| e | DLS vs NAcc coefficient (separate fits on the same trials) | Paired t-test | 8 mice | — | .391 |
| e | DLS previous reward response (per SD of last + running) vs 0 | One-sample t-test | 8 mice (both DLS and NAcc) | t(7) = -4.90† | .002 |
| e | NAcc previous reward response (per SD of last + running) vs 0 | One-sample t-test | 8 mice (both DLS and NAcc) | t(7) = -1.42† | .198 |
| e | DLS vs NAcc coefficient (separate fits on the same trials) | Paired t-test | 8 mice | — | .003 |
| e | DLS previous reward response (per SD of last + running) vs 0 | One-sample t-test | 8 mice (both DLS and NAcc) | t(7) = -4.59† | .003 |
| e | NAcc previous reward response (per SD of last + running) vs 0 | One-sample t-test | 8 mice (both DLS and NAcc) | t(7) = -0.86† | .420 |
| e | DLS vs NAcc coefficient (separate fits on the same trials) | Paired t-test | 8 mice | — | .083 |
| **Fig. 3** |  |  |  |  |  |
| a | Almorexant vs vehicle, low reward | Paired t-test | 13 mice | t(12) = -2.76 | .017 |
| a | Almorexant vs vehicle, high reward | Paired t-test | 13 mice | t(12) = -1.58 | .14 |
| a | Semaglutide vs vehicle, low reward | Paired t-test | 8 mice | t(7) = -3.35 | .012 |
| a | Semaglutide vs vehicle, high reward | Paired t-test | 8 mice | t(7) = -4.61 | .0025 |
| c | Almorexant - vehicle, reward block 3 (low reward) | Paired t-test | 13 mice | t(12) = -2.18 | .0499 |
| c | Almorexant - vehicle, reward block 5 (low reward) | Paired t-test | 13 mice | t(12) = -3.55 | .0040 |
| c | Semaglutide - vehicle, reward block 3 (low reward) | Paired t-test | 8 mice | t(7) = -2.82 | .026 |
| c | Semaglutide - vehicle, reward block 4 (high reward) | Paired t-test | 8 mice | t(7) = -4.97 | .0016 |
| c | Semaglutide - vehicle, reward block 5 (low reward) | Paired t-test | 8 mice | t(7) = -3.77 | .0069 |
| c | Semaglutide - vehicle, reward block 6 (high reward) | Paired t-test | 8 mice | t(7) = -5.24 | .0012 |
| c | Semaglutide - vehicle, reward block 7 (low reward) | Paired t-test | 8 mice | t(7) = -6.86 | $2.4 \times 10^{-4}$ |
| d | Almorexant vs vehicle, engaged state, both rewards (overall drug effect) | Paired t-test | 13 mice | t(12) = 1.03 | .33 |
| d | Almorexant vs vehicle, disengaged state, both rewards (overall drug effect) | Paired t-test | 12 mice | t(11) = 0.88 | .40 |
| d | Semaglutide vs vehicle, engaged state, both rewards (overall drug effect) | Paired t-test | 8 mice | t(7) = 0.68 | .52 |
| d | Semaglutide vs vehicle, disengaged state, both rewards (overall drug effect) | Paired t-test | 8 mice | t(7) = -1.42 | .20 |
| e | Almorexant vs vehicle, disengage hazard, low reward | Paired t-test | 13 mice | t(12) = 3.70 | .0030 |
| e | Almorexant vs vehicle, disengage hazard, high reward | Paired t-test | 13 mice | t(12) = 1.76 | .10 |
| e | Almorexant vs vehicle, re-engage hazard, low reward | Paired t-test | 13 mice | t(12) = -1.99 | .070 |
| e | Almorexant vs vehicle, re-engage hazard, high reward | Paired t-test | 13 mice | t(12) = -1.13 | .28 |
| f | Semaglutide vs vehicle, disengage hazard, low reward | Paired t-test | 8 mice | t(7) = 2.49 | .042 |
| f | Semaglutide vs vehicle, disengage hazard, high reward | Paired t-test | 8 mice | t(7) = 4.02 | .0051 |
| f | Semaglutide vs vehicle, re-engage hazard, low reward | Paired t-test | 8 mice | t(7) = -4.64 | .0024 |
| f | Semaglutide vs vehicle, re-engage hazard, high reward | Paired t-test | 8 mice | t(7) = -4.26 | .0037 |
| **Fig. 4** |  |  |  |  |  |
| a (μ) | engaged, low reward: almorexant vs vehicle | Paired t-test | 13 mice | t(12) = 4.82 | < .001 |
| a (μ) | engaged, high reward: almorexant vs vehicle | Paired t-test | 13 mice | t(12) = 6.28 | < .001 |
| a (μ) | engaged, vehicle: low vs high reward | Paired t-test | 13 mice | t(12) = -2.61 | .023 |
| a ($\mu$) | engaged, almorexant: low vs high reward | Paired t-test | 13 mice | t(12) = -2.35 | .036 |
| a ($\mu$) | disengaged, low reward: almorexant vs vehicle | Paired t-test | 13 mice | t(12) = 2.22 | .046 |
| a ($\mu$) | disengaged, high reward: almorexant vs vehicle | Paired t-test | 13 mice | t(12) = 3.64 | .003 |
| a ($\mu$) | disengaged, vehicle: low vs high reward | Paired t-test | 13 mice | t(12) = -0.84 | .416 |
| a ($\mu$) | disengaged, almorexant: low vs high reward | Paired t-test | 13 mice | t(12) = -1.48 | .164 |
| a ($\sigma$) | engaged, low reward: almorexant vs vehicle | Paired t-test | 13 mice | t(12) = 5.23 | < .001 |
| a ($\sigma$) | engaged, high reward: almorexant vs vehicle | Paired t-test | 13 mice | t(12) = 6.07 | < .001 |
| a ($\sigma$) | engaged: drug × reward interaction | Paired t-test | 13 mice | t(12) = 3.06 | .0100 |
| a ($\sigma$) | engaged, vehicle: low vs high reward (narrowing) | Paired t-test | 13 mice | t(12) = 13.91 | < .001 |
| a ($\sigma$) | engaged, almorexant: low vs high reward (narrowing) | Paired t-test | 13 mice | t(12) = 7.79 | < .001 |
| a ($\sigma$) | disengaged, low reward: almorexant vs vehicle | Paired t-test | 13 mice | t(12) = 3.56 | .004 |
| a ($\sigma$) | disengaged, high reward: almorexant vs vehicle | Paired t-test | 13 mice | t(12) = 1.88 | .085 |
| a ($\sigma$) | disengaged, vehicle: low vs high reward (narrowing) | Paired t-test | 13 mice | t(12) = 5.59 | < .001 |
| a ($\sigma$) | disengaged, almorexant: low vs high reward (narrowing) | Paired t-test | 13 mice | t(12) = 4.50 | < .001 |
| d | pure indirect effect via disengage hazard (almorexant pupil distribution under vehicle hazards), low reward | One-sample t-test | 13 mice | — | < .001 |
| d | pure indirect effect via re-engage hazard, low reward | One-sample t-test | 13 mice | — | .002 |
| e (left) | P(disengage) almorexant vs vehicle, lowest of 6 pupil bins, low reward | Paired t-test | 13 mice (43,671 transitions) | — | .003 |
| e (left) | P(disengage) almorexant vs vehicle, pupil bins 2–6, low reward | Paired t-test | 13 mice | — | bin 2 .665, 3 .208<br>.532, 6 .536 |
| e (right) | almorexant vs vehicle, VTA oxLight bin 1 of 5, bottom 30 % pupil, low reward | Paired t-test | 6 VTA mice | — | .13 |
| e (right) | almorexant vs vehicle, VTA oxLight bin 2 of 5, bottom 30 % pupil, low reward | Paired t-test | 6 VTA mice | — | .12 |
| e (right) | almorexant vs vehicle, VTA oxLight bin 3 of 5, bottom 30 % pupil, low reward | Paired t-test | 7 VTA mice | — | .059 |
| e (right) | almorexant vs vehicle, VTA oxLight bin 4 of 5, bottom 30 % pupil, low reward | Paired t-test | 7 VTA mice | — | .040 |
| e (right) | almorexant vs vehicle, VTA oxLight bin 5 of 5, bottom 30 % pupil, low reward | Paired t-test | 7 VTA mice | — | .012 |
| f | NACC, low reward: almorexant vs vehicle | Paired t-test | 8 mice | t(7) = -0.47 | .650 |
| f | NACC, high reward: almorexant vs vehicle | Paired t-test | 8 mice | t(7) = -0.87 | .415 |
| f | DLS, low reward: almorexant vs vehicle | Paired t-test | 8 mice | t(7) = -0.13 | .900 |
| f | DLS, high reward: almorexant vs vehicle | Paired t-test | 8 mice | t(7) = -0.75 | .475 |
| g | pure indirect share of almorexant effect on P(disengaged): pupil, low reward (filled) | One-sample t-test | 8 mice | — | < .001 |
| g | pure indirect share of almorexant effect on P(disengaged): pupil, high reward (open) | One-sample t-test | 8 mice | — | .027 |
| g | pure indirect share of almorexant effect on P(disengaged): NACC, low reward | One-sample t-test | 8 mice | — | .136 |
| g | pure indirect share of almorexant effect on P(disengaged): NACC, high reward | One-sample t-test | 8 mice | — | .279 |
| g | pure indirect share of almorexant effect on P(disengaged): DLS, low reward | One-sample t-test | 8 mice | — | .608 |
| g | pure indirect share of almorexant effect on P(disengaged): DLS, high reward | One-sample t-test | 8 mice | — | .543 |

**Fig. 5**

|  |  |  |  |  |  |
| --- | --- | --- | --- | --- | --- |
| b (right) | orexin 5 Hz stim – control vs 0 | One-sample t-test | 8 orexin mice | t(7) = 2.77 | .028 |
| b (right) | orexin 20 Hz stim – control vs 0 | One-sample t-test | 8 orexin mice | t(7) = 4.48 (vs heat: Welch t(7.5) = 4.93, P = .0014) | .0029 |
| c | 5 Hz: laser days vs sham | Paired t-test | 8 orexin mice | t(7) = 2.63 | .034 |
| c | 20 Hz: laser days vs sham | Paired t-test | 8 orexin mice | t(7) = 4.56 | .003 |
| d (5 Hz) | low reward: laser vs sham | Paired t-test | 8 orexin mice | t(7) = -3.65 | .008 |
| d (5 Hz) | high reward: laser vs sham | Paired t-test | 8 orexin mice | t(7) = -1.64 | .145 |
| d (5 Hz) | bracket: low vs high (sham-corrected effect) | Paired t-test | 8 orexin mice | t(7) = -2.44 | .044 |
| d (20 Hz) | low reward: laser vs sham | Paired t-test | 8 orexin mice | t(7) = -2.62 | .034 |
| d (20 Hz) | high reward: laser vs sham | Paired t-test | 8 orexin mice | t(7) = -2.51 | .040 |
| d (20 Hz) | bracket: low vs high (sham-corrected) | Paired t-test | 8 orexin mice | t(7) = -1.97 | .090 |
| e (low reward) | 5 Hz vs control at matched pupil, laser – sham | Paired t-test | 7 orexin mice (enough trials per bin) | t(6) = -1.54 | .174 |
| e (low reward) | 20 Hz vs control at matched pupil, laser – sham | Paired t-test | 7 orexin mice | t(6) = -0.16 | .882 |

**Fig. 6**

|  |  |  |  |  |  |
| --- | --- | --- | --- | --- | --- |
| a (inset, vehicle) | reward AUC 0-3 s, trials 2-3 minus trial 1 after a switch into high (rise), vehicle | Paired t-test | 8 mice | — | < .001 |
| a (inset, semaglutide) | rise trials 2-3 minus trial 1 after a switch into high, semaglutide | Paired t-test | 8 mice | — | .25 |
| d | integrator response to a reward step, 1st switch into high, trials 1-30, semaglutide vs vehicle | Paired t-test | 8 mice | t(7) = -3.54 | .009 |
| e | model step at 1st switch vs session trend of P(disengaged), vehicle sessions of both experiments | Permutation | 120 sessions, 9 mice (16 mouse × experiment cells) | r = -0.24 | .011 |
| e | model step at 1st switch vs session trend of P(disengaged), semaglutide sessions | Permutation | 45 sessions, 8 mice | r = -0.60 | < .001 |
| e (inset) | share of semaglutide's effect on the P(disengaged) session trend mediated by the step, vehicle rule (combined-vehicle link) | Bootstrap over mice | 8 semaglutide mice (+ 9 vehicle mice for the vehicle link) | bootstrap | < .001 |
| e (inset) | share of semaglutide's effect on the P(disengaged) session trend mediated by the step, drug link (semaglutide sessions) | Bootstrap over mice | 8 semaglutide mice (+ 9 vehicle mice for the vehicle link) | bootstrap | < .001 |
| e (inset) | share of semaglutide's effect on the P(disengaged) session trend mediated by the step, mean of both links | Bootstrap over mice | 8 semaglutide mice (+ 9 vehicle mice for the vehicle link) | bootstrap | < .001 |
| f | NACC reward AUC -0.5 to 1.5 s, high-reward blocks, semaglutide vs vehicle | Paired t-test | 8 mice | t(7) = -2.69 | .031 |
| f | NACC reward AUC -0.5 to 1.5 s, low-reward blocks, semaglutide vs vehicle | Paired t-test | 8 mice | t(7) = 0.07 | .94 |
| g | DLS reward AUC -0.5 to 1.0 s, high-reward blocks, semaglutide vs vehicle | Paired t-test | 8 mice | t(7) = -3.59 | .009 |
| g | DLS reward AUC -0.5 to 1.0 s, low-reward blocks, semaglutide vs vehicle | Paired t-test | 8 mice | t(7) = 0.14 | .89 |
| g (inset) | share of semaglutide's increase in P(disengaged) mediated by the trial-level DLS response, vehicle rule (pure indirect) | Bootstrap over mice | 8 mice | bootstrap | .50 |
| g (inset) | share of semaglutide's increase in P(disengaged) mediated by the trial-level DLS response, semaglutide rule (total indirect) | Bootstrap over mice | 8 mice | bootstrap | .008 |
| g (inset) | share of semaglutide's increase in P(disengaged) mediated by the trial-level DLS response, mean of both | Bootstrap over mice | 8 mice | bootstrap | .032 |
| h (engaged) | drug vs vehicle, low reward | Paired t-test | 8 mice | t(7) = -3.91 | .006 |
| h (engaged) | drug vs vehicle, high reward | Paired t-test | 8 mice | t(7) = -4.37 | .003 |
| h (engaged) | low vs high reward, vehicle | Paired t-test | 8 mice | t(7) = -2.51 | .040 |
| h (engaged) | low vs high reward, semaglutide | Paired t-test | 8 mice | t(7) = -1.51 | .17 |
| h (disengaged) | drug vs vehicle, low reward | Paired t-test | 8 mice | t(7) = 0.03 | .98 |
| h (disengaged) | drug vs vehicle, high reward | Paired t-test | 8 mice | t(7) = -0.43 | .68 |
| h (disengaged) | low vs high reward, vehicle | Paired t-test | 8 mice | t(7) = 0.06 | .95 |
| h (disengaged) | low vs high reward, semaglutide | Paired t-test | 8 mice | t(7) = 1.00 | .35 |

##### Extended Data Fig. 1

|  |  |  |  |  |  |
| --- | --- | --- | --- | --- | --- |
| f (top) | vehicle (almorexant experiment): engaged vs disengaged | Paired t-test | 8 mice | t(7) = 3.07 | .018 |
| f (top) | almorexant: engaged vs disengaged | Paired t-test | 8 mice | t(7) = 4.14 | .0044 |
| f (top) | engaged: almorexant vs vehicle | Paired t-test | 8 mice | t(7) = -3.19 | .015 |
| f (bottom) | vehicle (semaglutide experiment): engaged vs disengaged | Paired t-test | 8 mice | t(7) = 6.99 | $2.1 \times 10^{-4}$ |
| f (bottom) | semaglutide: engaged vs disengaged | Paired t-test | 8 mice | t(7) = 5.42 | $< .001 (9.9 \times 10^{-4})$ |
| g (top) | almorexant - vehicle, low reward, engaged | One-sample t-test | 8 mice | t(7) = -1.90 | .099 |
| g (top) | almorexant - vehicle, low reward, disengaged | One-sample t-test | 8 mice | t(7) = -0.32 | .76 |
| g (top) | almorexant - vehicle, high reward, engaged | One-sample t-test | 8 mice | t(7) = -1.35 | .22 |
| g (top) | almorexant - vehicle, high reward, disengaged | One-sample t-test | 8 mice | t(7) = 0.30 | .77 |
| g (bottom) | semaglutide - vehicle, low reward, engaged | One-sample t-test | 8 mice | t(7) = 0.97 | .36 |
| g (bottom) | semaglutide - vehicle, low reward, disengaged | One-sample t-test | 8 mice | t(7) = -0.99 | .35 |
| g (bottom) | semaglutide - vehicle, high reward, engaged | One-sample t-test | 8 mice | t(7) = 0.40 | .70 |
| g (bottom) | semaglutide - vehicle, high reward, disengaged | One-sample t-test | 8 mice | t(7) = -4.51 | .0028 |

##### Extended Data Fig. 2

|  |  |  |  |  |  |
| --- | --- | --- | --- | --- | --- |
| a | high vs low reward block | Paired t-test | 8 mice | t(7) = -2.72 | .030 |
| a | high-reward blocks: curvature (quadratic term) across the 6 bins | One-sample t-test | 8 mice | t(7) = -3.06 | .018 |
| a | low-reward blocks: curvature (quadratic term) across the 6 bins | One-sample t-test | 8 mice | t(7) = -4.40 | .003 |
| a | low-reward blocks: curvature (quadratic term) across the 6 bins | One-sample t-test | 8 mice | t(7) = 2.52 | .040 |
| c | high vs low reward block | Paired t-test | 9 mice | t(8) = -1.93 | .089 |
| c | low-reward blocks: linear trend across the 6 bins | One-sample t-test | 9 mice | t(8) = -2.80 | .023 |
| c | high-reward blocks: curvature (quadratic term) across the 6 bins | One-sample t-test | 9 mice | t(8) = -2.31 | .0496 |
| c | low-reward blocks: linear trend across the 6 bins | One-sample t-test | 9 mice | t(8) = 2.70 | .027 |
| d | high vs low reward block | Paired t-test | 9 mice | t(8) = 5.46 | $6.0 \times 10^{-4}$ |
| d | low-reward blocks: curvature (quadratic term) across the 6 bins | One-sample t-test | 9 mice | t(8) = -2.83 | .022 |
| d | low-reward blocks: linear trend across the 6 bins | One-sample t-test | 9 mice | t(8) = 2.53 | .036 |
| d | low-reward blocks: curvature (quadratic term) across the 6 bins | One-sample t-test | 9 mice | t(8) = -2.66 | .029 |
| d | low-reward blocks: curvature (quadratic term) across the 6 bins | One-sample t-test | 9 mice | t(8) = 2.72 | .026 |
| e | per-trial correlation pupil vs SNc, engaged trials (lower triangle) | Fisher-z r, one-sample t-test (BH-corrected) | 8 mice | r = +0.010 | .664 (q = .738) |
| e | per-trial correlation pupil vs VTA, engaged trials (lower triangle) | Fisher-z r, one-sample t-test (BH-corrected) | 9 mice | r = -0.004 | .878 (q = .878) |
| e | per-trial correlation pupil vs NAcc, engaged trials (lower triangle) | Fisher-z r, one-sample t-test (BH-corrected) | 9 mice | r = +0.094 | .149 (q = .298) |
| e | per-trial correlation pupil vs DLS, engaged trials (lower triangle) | Fisher-z r, one-sample t-test (BH-corrected) | 9 mice | r = -0.097 | .044 (q = .146) |
| e | per-trial correlation SNc vs VTA, engaged trials (lower triangle) | Fisher-z r, one-sample t-test (BH-corrected) | 8 mice | r = +0.232 | .113 (q = .282) |
| e | per-trial correlation SNc vs NAcc, engaged trials (lower triangle) | Fisher-z r, one-sample t-test (BH-corrected) | 8 mice | r = +0.014 | .446 (q = .557) |
| e | per-trial correlation SNc vs DLS, engaged trials (lower triangle) | Fisher-z r, one-sample t-test (BH-corrected) | 8 mice | r = -0.158 | .011 (q = .057) |
| e | per-trial correlation VTA vs NAcc, engaged trials (lower triangle) | Fisher-z r, one-sample t-test (BH-corrected) | 9 mice | r = -0.046 | .286 (q = .408) |
| e | per-trial correlation VTA vs DLS, engaged trials (lower triangle) | Fisher-z r, one-sample t-test (BH-corrected) | 9 mice | r = -0.079 | .216 (q = .359) |
| e | per-trial correlation NAcc vs DLS, engaged trials (lower triangle) | Fisher-z r, one-sample t-test (BH-corrected) | 9 mice | r = +0.285 | .006 (q = .057) |
| e | per-trial correlation pupil vs SNc, disengaged trials (upper triangle) | Fisher-z r, one-sample t-test (BH-corrected) | 8 mice | r = -0.036 | .399 (q = .499) |
| e | per-trial correlation pupil vs VTA, disengaged trials (upper triangle) | Fisher-z r, one-sample t-test (BH-corrected) | 9 mice | r = -0.056 | .256 (q = .392) |
| e | per-trial correlation pupil vs NAcc, disengaged trials (upper triangle) | Fisher-z r, one-sample t-test (BH-corrected) | 9 mice | r = -0.053 | .487 (q = .541) |
| e | per-trial correlation pupil vs DLS, disengaged trials (upper triangle) | Fisher-z r, one-sample t-test (BH-corrected) | 9 mice | r = -0.107 | .083 (q = .166) |
| e | per-trial correlation SNc vs VTA, disengaged trials (upper triangle) | Fisher-z r, one-sample t-test (BH-corrected) | 8 mice | r = +0.343 | .006 (q = .020) |
| e | per-trial correlation SNc vs NAcc, disengaged trials (upper triangle) | Fisher-z r, one-sample t-test (BH-corrected) | 7 mice | r = +0.029 | .274 (q = .392) |
| e | per-trial correlation SNc vs DLS, disengaged trials (upper triangle) | Fisher-z r, one-sample t-test (BH-corrected) | 8 mice | r = -0.094 | .0014 (q = .007) |
| e | per-trial correlation VTA vs NAcc, disengaged trials (upper triangle) | Fisher-z r, one-sample t-test (BH-corrected) | 9 mice | r = -0.033 | .600 (q = .600) |
| e | per-trial correlation VTA vs DLS, disengaged trials (upper triangle) | Fisher-z r, one-sample t-test (BH-corrected) | 8 mice | r = -0.152 | .042 (q = .106) |
| e | per-trial correlation NAcc vs DLS, disengaged trials (upper triangle) | Fisher-z r, one-sample t-test (BH-corrected) | 8 mice | r = +0.386 | $1.6 \times 10^{-4}$ (q = .0) |
| f | share of the high - low block effect carried by pupil, baseline level (+ movement / licking covariates, all trials) | One-sample t-test; bootstrap over mice | 14 mice | — | .153 |
| f | share of the high - low block effect carried by pupil, baseline level (+ movement / licking covariates, all trials) | One-sample t-test; bootstrap over mice | 14 mice | — | .135 |
| f | share of the high - low block effect carried by pupil, baseline level (+ movement / licking covariates, all trials) | One-sample t-test; bootstrap over mice | 14 mice | — | .131 |
| f | share of the high - low block effect carried by SNc, baseline level (+ movement / licking covariates, all trials) | One-sample t-test; bootstrap over mice | 8 mice | — | .474 |
| f | share of the high - low block effect carried by SNc, baseline level (+ movement / licking covariates, all trials) | One-sample t-test; bootstrap over mice | 8 mice | — | .307 |
| f | share of the high - low block effect carried by SNc, baseline level (+ movement / licking covariates, all trials) | One-sample t-test; bootstrap over mice | 8 mice | — | .996 |
| f | share of the high - low block effect carried by VTA, baseline level (+ movement / licking covariates, all trials) | One-sample t-test; bootstrap over mice | 9 mice | — | .157 |
| f | share of the high - low block effect carried by VTA, baseline level (+ movement / licking covariates, all trials) | One-sample t-test; bootstrap over mice | 9 mice | — | .134 |
| f | share of the high - low block effect carried by VTA, baseline level (+ movement / licking covariates, all trials) | One-sample t-test; bootstrap over mice | 9 mice | — | .117 |
| f | share of the high - low block effect carried by NAcc, last + running reward resp. (+ movement / licking covariates, all trials) | One-sample t-test; bootstrap over mice | 9 mice | — | .116 |
| f | share of the high - low block effect carried by NAcc, last + running reward resp. (+ movement / licking covariates, all trials) | One-sample t-test; bootstrap over mice | 9 mice | — | .262 |
| f | share of the high - low block effect carried by NAcc, last + running reward resp. (+ movement / licking covariates, all trials) | One-sample t-test; bootstrap over mice | 9 mice | — | .126 |
| f | share of the high - low block effect carried by DLS, last + running reward resp. (+ movement / licking covariates, all trials) | One-sample t-test; bootstrap over mice | 9 mice | — | .039 |
| f | share of the high - low block effect carried by DLS, last + running reward resp. (+ movement / licking covariates, all trials) | One-sample t-test; bootstrap over mice | 9 mice | — | .013 |
| f | share of the high - low block effect carried by DLS, last + running reward resp. (+ movement / licking covariates, all trials) | One-sample t-test; bootstrap over mice | 9 mice | — | .021 |
| f | share of the high - low block effect carried by all five signals (+ movement / licking covariates, all trials) | One-sample t-test; bootstrap over mice | 9 mice | — | .026 |
| f | share of the high - low block effect carried by all five signals (+ movement / licking covariates, all trials) | One-sample t-test; bootstrap over mice | 9 mice | — | .025 |
| f | share of the high - low block effect carried by all five signals (+ movement / licking covariates, all trials) | One-sample t-test; bootstrap over mice | 9 mice | — | .011 |
| g | pupil: slope on the recent reward rate (tau 10 trials) vs 0 | One-sample t-test | 14 mice | — | .058 |
| g | SNc: slope on the recent reward rate (tau 10 trials) vs 0 | One-sample t-test | 8 mice | — | .013 |
| g | VTA: slope on the recent reward rate (tau 10 trials) vs 0 | One-sample t-test | 9 mice | — | .048 |
| g | NAcc: slope on the recent reward rate (tau 10 trials) vs 0 | One-sample t-test | 9 mice | — | .013 |
| g | DLS: slope on the recent reward rate (tau 10 trials) vs 0 | One-sample t-test | 8 mice | — | .496 |
| h | pupil × SNc interaction, disengage hazard, low-reward blocks | One-sample t-test (BH-corrected) | 8 mice | — | .594 (q = .659) |
| h | pupil × VTA interaction, disengage hazard, low-reward blocks | One-sample t-test (BH-corrected) | 9 mice | — | .483 (q = .659) |
| h | pupil × NAcc interaction, disengage hazard, low-reward blocks | One-sample t-test (BH-corrected) | 9 mice | — | .580 (q = .659) |
| h | pupil × DLS interaction, disengage hazard, low-reward blocks | One-sample t-test (BH-corrected) | 9 mice | — | .462 (q = .659) |
| h | SNc × VTA interaction, disengage hazard, low-reward blocks | One-sample t-test (BH-corrected) | 8 mice | — | .329 (q = .658) |
| h | SNc × NAcc interaction, disengage hazard, low-reward blocks | One-sample t-test (BH-corrected) | 8 mice | — | .209 (q = .557) |
| h | SNc × DLS interaction, disengage hazard, low-reward blocks | One-sample t-test (BH-corrected) | 8 mice | — | .861 (q = .861) |
| h | VTA × NAcc interaction, disengage hazard, low-reward blocks | One-sample t-test (BH-corrected) | 9 mice | — | .223 (q = .557) |
| h | VTA × DLS interaction, disengage hazard, low-reward blocks | One-sample t-test (BH-corrected) | 9 mice | — | .033 (q = .280) |
| h | NAcc × DLS interaction, disengage hazard, low-reward blocks | One-sample t-test (BH-corrected) | 9 mice | — | .056 (q = .280) |
| h | pupil × SNc interaction, disengage hazard, high-reward blocks | One-sample t-test (BH-corrected) | 8 mice | — | .525 (q = .815) |
| h | pupil × VTA interaction, disengage hazard, high-reward blocks | One-sample t-test (BH-corrected) | 9 mice | — | .009 (q = .045) |
| h | pupil × NAcc interaction, disengage hazard, high-reward blocks | One-sample t-test (BH-corrected) | 9 mice | — | .002 (q = .024) |
| h | pupil × DLS interaction, disengage hazard, high-reward blocks | One-sample t-test (BH-corrected) | 9 mice | — | .631 (q = .815) |
| h | SNc × VTA interaction, disengage hazard, high-reward blocks | One-sample t-test (BH-corrected) | 8 mice | — | .950 (q = .950) |
| h | SNc × NAcc interaction, disengage hazard, high-reward blocks | One-sample t-test (BH-corrected) | 8 mice | — | .652 (q = .815) |
| h | SNc × DLS interaction, disengage hazard, high-reward blocks | One-sample t-test (BH-corrected) | 8 mice | — | .384 (q = .815) |
| h | VTA × NAcc interaction, disengage hazard, high-reward blocks | One-sample t-test (BH-corrected) | 9 mice | — | .884 (q = .950) |
| h | VTA × DLS interaction, disengage hazard, high-reward blocks | One-sample t-test (BH-corrected) | 9 mice | — | .461 (q = .815) |
| h | NAcc × DLS interaction, disengage hazard, high-reward blocks | One-sample t-test (BH-corrected) | 9 mice | — | .468 (q = .815) |
| h | pupil × SNc interaction, re-engage hazard, low-reward blocks | One-sample t-test (BH-corrected) | 8 mice | — | .041 (q = .408) |
| h | pupil × VTA interaction, re-engage hazard, low-reward blocks | One-sample t-test (BH-corrected) | 9 mice | — | .199 (q = .714) |
| h | pupil × NAcc interaction, re-engage hazard, low-reward blocks | One-sample t-test (BH-corrected) | 9 mice | — | .214 (q = .714) |
| h | pupil × DLS interaction, re-engage hazard, low-reward blocks | One-sample t-test (BH-corrected) | 9 mice | — | .292 (q = .730) |
| h | SNc × VTA interaction, re-engage hazard, low-reward blocks | One-sample t-test (BH-corrected) | 8 mice | — | .591 (q = .940) |
| h | SNc × NAcc interaction, re-engage hazard, low-reward blocks | One-sample t-test (BH-corrected) | 8 mice | — | .802 (q = .963) |
| h | SNc × DLS interaction, re-engage hazard, low-reward blocks | One-sample t-test (BH-corrected) | 8 mice | — | .504 (q = .940) |
| h | VTA × NAcc interaction, re-engage hazard, low-reward blocks | One-sample t-test (BH-corrected) | 9 mice | — | .967 (q = .967) |
| h | VTA × DLS interaction, re-engage hazard, low-reward blocks | One-sample t-test (BH-corrected) | 9 mice | — | .658 (q = .940) |
| h | NAcc × DLS interaction, re-engage hazard, low-reward blocks | One-sample t-test (BH-corrected) | 9 mice | — | .867 (q = .963) |
| h | pupil × SNc interaction, re-engage hazard, high-reward blocks | One-sample t-test (BH-corrected) | 7 mice | — | .833 (q = .924) |
| h | pupil × VTA interaction, re-engage hazard, high-reward blocks | One-sample t-test (BH-corrected) | 8 mice | — | .193 (q = .873) |
| h | pupil × NAcc interaction, re-engage hazard, high-reward blocks | One-sample t-test (BH-corrected) | 8 mice | — | .628 (q = .873) |
| h | pupil × DLS interaction, re-engage hazard, high-reward blocks | One-sample t-test (BH-corrected) | 8 mice | — | .646 (q = .873) |
| h | SNc × VTA interaction, re-engage hazard, high-reward blocks | One-sample t-test (BH-corrected) | 7 mice | — | .372 (q = .873) |
| h | SNc × NAcc interaction, re-engage hazard, high-reward blocks | One-sample t-test (BH-corrected) | 7 mice | — | .328 (q = .873) |
| h | SNc × DLS interaction, re-engage hazard, high-reward blocks | One-sample t-test (BH-corrected) | 7 mice | — | .658 (q = .873) |
| h | VTA × NAcc interaction, re-engage hazard, high-reward blocks | One-sample t-test (BH-corrected) | 8 mice | — | .924 (q = .924) |
| h | VTA × DLS interaction, re-engage hazard, high-reward blocks | One-sample t-test (BH-corrected) | 8 mice | — | .506 (q = .873) |
| h | NAcc × DLS interaction, re-engage hazard, high-reward blocks | One-sample t-test (BH-corrected) | 8 mice | — | .698 (q = .873) |

**Extended Data Fig. 3**

|  |  |  |  |  |  |
| --- | --- | --- | --- | --- | --- |
| a | Almorexant vs vehicle | Paired t-test | 13 mice | t(12) = -2.27 | .043 |
| a | Semaglutide vs vehicle | Paired t-test | 8 mice | t(7) = -3.36 | .012 |
| b | Almorexant, low - high reward RT vs 0 | One-sample t-test | 13 mice | t(12) = 5.28 | < .001 |
| b | Vehicle (almorexant experiment), low - high reward RT vs 0 | One-sample t-test | 13 mice | t(12) = 3.94 | .0020 |
| b | Almorexant vs vehicle, low - high reward RT | Paired t-test | 13 mice | t(12) = 4.69 | < .001 |
| b | Semaglutide, low - high reward RT vs 0 | One-sample t-test | 8 mice | t(7) = 2.79 | .027 |
| b | Vehicle (semaglutide experiment), low - high reward RT vs 0 | One-sample t-test | 8 mice | t(7) = 4.05 | .0049 |
| b | Semaglutide vs vehicle, low - high reward RT | Paired t-test | 8 mice | t(7) = 1.14 | .29 |
| f | Almorexant vs vehicle, constant reward | Paired t-test | 8 mice | t(7) = 2.89 | .023 |
| f | Semaglutide vs vehicle, constant reward | Paired t-test | 8 mice | t(7) = -6.62 | $3 \times 10^{-4}$ |
| g | Almorexant vs vehicle, constant reward | Paired t-test | 8 mice | t(7) = 1.25 | .25 |
| g | Semaglutide vs vehicle, constant reward | Paired t-test | 8 mice | t(7) = -6.95 | $2 \times 10^{-4}$ |
| h | Almorexant vs vehicle, constant reward, disengage hazard | Paired t-test | 8 mice | t(7) = -3.17 | .016 |
| h | Almorexant vs vehicle, constant reward, re-engage hazard | Paired t-test | 8 mice | t(7) = 0.45 | .67 |
| h | Semaglutide vs vehicle, constant reward, disengage hazard | Paired t-test | 8 mice | t(7) = 3.12 | .017 |
| h | Semaglutide vs vehicle, constant reward, re-engage hazard | Paired t-test | 8 mice | t(7) = -4.54 | .0027 |

**Extended Data Fig. 4**

|  |  |  |  |  |  |
| --- | --- | --- | --- | --- | --- |
| a (middle) | slope of baseline pupil over the session, almorexant vs vehicle, all trials | Paired t-test | 13 mice | t(12) = -2.86 | .014 |
| a (middle) | slope of baseline pupil over the session, almorexant vs vehicle, engaged trials | Paired t-test | 13 mice | t(12) = -3.08 | .009 |
| d | pupil-movement coupling slope, low reward: almorexant vs vehicle | Paired t-test | 8 mice (wheel) | t(7) = 3.54 | .0095 |
| d | pupil-movement coupling slope, high reward: almorexant vs vehicle | Paired t-test | 8 mice | t(7) = 1.64 | .145 |
| e | baseline pupil on trials without movement, engaged: almorexant vs vehicle | Paired t-test | 8 mice | t(7) = 3.01 | .020 |
| e | baseline pupil on trials without movement, disengaged: almorexant vs vehicle | Paired t-test | 6 mice | t(5) = -4.93 | .004 |
| g | dilation 0–2 s after reward from pre-stimulus baseline, low reward: almorexant vs vehicle | Paired t-test | 13 mice | t(12) = 3.03 | .011 |
| g | dilation 0–2 s after reward from pre-stimulus baseline, high reward: almorexant vs vehicle | Paired t-test | 13 mice | t(12) = 1.17 | .264 |
| h | dilation 0–2 s after reward from pre-reward pupil, low reward: almorexant vs vehicle | Paired t-test | 13 mice | t(12) = 3.32 | .006 |
| h | dilation 0–2 s after reward from pre-reward pupil, high reward: almorexant vs vehicle | Paired t-test | 13 mice | t(12) = 0.75 | .465 |
| i | almorexant vs vehicle: dark pupil, start of session | Paired t-test | 6 mice | t(5) = -2.72 | .042 |
| i | almorexant vs vehicle: dark pupil, end of session | Paired t-test | 5 mice | t(4) = -1.53 | .20 |
| i | almorexant vs vehicle: light pupil, start of session | Paired t-test | 6 mice | t(5) = 1.14 | .30 |
| i | almorexant vs vehicle: light pupil, end of session | Paired t-test | 5 mice | t(4) = -0.65 | .55 |
| i | almorexant vs vehicle: constriction t50, start of session | Paired t-test | 6 mice | t(5) = 0.16 | .876 |
| i | almorexant vs vehicle: constriction t50, end of session | Paired t-test | 5 mice | t(4) = -0.35 | .741 |
| i | almorexant vs vehicle: task baseline pupil (laser-free trials, no-laser days, both rewards) | Paired t-test | 5 mice | t(4) = 4.36 | .012 |

**Extended Data Fig. 5**

|  |  |  |  |  |  |
| --- | --- | --- | --- | --- | --- |
| a (per-mouse fits) | generative arousal route (pure indirect): almorexant, low reward, via disengage hazard | Wilcoxon signed-rank | 13 mice | signed-rank | .021 |
| a (per-mouse fits) | generative arousal route (pure indirect): almorexant, low reward, via re-engage hazard | Wilcoxon signed-rank | 13 mice | signed-rank | .068 |
| a (per-mouse fits) | generative arousal route (pure indirect): almorexant, high reward, via disengage hazard | Wilcoxon signed-rank | 13 mice | signed-rank | .017 |
| a (per-mouse fits) | generative arousal route (pure indirect): almorexant, high reward, via re-engage hazard | Wilcoxon signed-rank | 13 mice | signed-rank | .340 |
| a (per-mouse fits) | generative arousal route (pure indirect): semaglutide, low reward, via disengage hazard | Wilcoxon signed-rank | 8 mice | signed-rank | .195 |
| a (per-mouse fits) | generative arousal route (pure indirect): semaglutide, low reward, via re-engage hazard | Wilcoxon signed-rank | 8 mice | signed-rank | .383 |
| a (per-mouse fits) | generative arousal route (pure indirect): semaglutide, high reward, via disengage hazard | Wilcoxon signed-rank | 8 mice | signed-rank | .742 |
| a (per-mouse fits) | generative arousal route (pure indirect): semaglutide, high reward, via re-engage hazard | Wilcoxon signed-rank | 8 mice | signed-rank | .844 |
| a (hierarchical fit) | generative arousal route (pure indirect): almorexant, low reward, via disengage hazard | Bootstrap over mice | 13 mice | bootstrap | .0010 |
| a (hierarchical fit) | generative arousal route (pure indirect): almorexant, low reward, via re-engage hazard | Bootstrap over mice | 13 mice | bootstrap | .0010 |
| a (hierarchical fit) | generative arousal route (pure indirect): almorexant, high reward, via disengage hazard | Bootstrap over mice | 13 mice | bootstrap | .0010 |
| a (hierarchical fit) | generative arousal route (pure indirect): almorexant, high reward, via re-engage hazard | Bootstrap over mice | 13 mice | bootstrap | .036 |
| a (hierarchical fit) | generative arousal route (pure indirect): semaglutide, low reward, via disengage hazard | Bootstrap over mice | 8 mice | bootstrap | .65 |
| a (hierarchical fit) | generative arousal route (pure indirect): semaglutide, low reward, via re-engage hazard | Bootstrap over mice | 8 mice | bootstrap | .952 |
| a (hierarchical fit) | generative arousal route (pure indirect): semaglutide, high reward, via disengage hazard | Bootstrap over mice | 8 mice | bootstrap | .80 |
| a (hierarchical fit) | generative arousal route (pure indirect): semaglutide, high reward, via re-engage hazard | Bootstrap over mice | 8 mice | bootstrap | .494 |
| b (low reward) | P(engaged) almorexant vs vehicle, pupil bin 1 of 6, low reward | Paired t-test | 13 mice | t(12) = -3.43 | .005 |
| b (low reward) | P(engaged) almorexant vs vehicle, pupil bin 2 of 6, low reward | Paired t-test | 13 mice | t(12) = -2.28 | .042 |
| b (low reward) | P(engaged) almorexant vs vehicle, pupil bin 4 of 6, low reward | Paired t-test | 13 mice | t(12) = -2.21 | .047 |
| b (low reward) | P(engaged) almorexant vs vehicle, bins 3, 5, 6, low reward | Paired t-test | 13 mice | — | bin 3 .070, 5 .116 |
| b (high reward) | P(engaged) almorexant vs vehicle, bins 1–6, high reward | Paired t-test | 13 mice | — | P = .155 to .509 |
| c | engaged, low reward: semaglutide vs vehicle | Paired t-test | 8 mice | t(7) = 0.33 | .751 |
| c | engaged, high reward: semaglutide vs vehicle | Paired t-test | 8 mice | t(7) = 1.33 | .224 |
| c | engaged, vehicle: low vs high reward (narrowing) | Paired t-test | 8 mice | t(7) = 7.63 | < .001 |
| c | engaged, semaglutide: low vs high reward (narrowing) | Paired t-test | 8 mice | t(7) = 3.84 | .006 |
| c | disengaged, low reward: semaglutide vs vehicle | Paired t-test | 8 mice | t(7) = 3.51 | .0098 |
| c | disengaged, high reward: semaglutide vs vehicle | Paired t-test | 8 mice | t(7) = 3.49 | .0101 |
| c | disengaged, vehicle: low vs high reward (narrowing) | Paired t-test | 8 mice | t(7) = 2.80 | .026 |
| c | disengaged, semaglutide: low vs high reward (narrowing) | Paired t-test | 8 mice | t(7) = 0.81 | .446 |

##### Extended Data Fig. 6

|  |  |  |  |  |  |
| --- | --- | --- | --- | --- | --- |
| a (left, bottom 30 % pupil) | almorexant vs vehicle, VTA oxLight bin 4 (z +0.43), low reward | Paired t-test | 7 VTA mice (5/7 lower) | — | .040 |
| a (left, bottom 30 % pupil) | almorexant vs vehicle, VTA oxLight bin 5 (z +1.42), low reward | Paired t-test | 7 VTA mice (7/7 lower) | t(6) = -3.54 (same cell, 'high release' line) | .012 |
| a (left) | bins 1-3 (z -1.25, -0.55, -0.09) | Paired t-test | 6, 6, 7 mice | — | .127 / .119 / .059 |
| a (right, top 30 % pupil) | five bins | Paired t-test | 7 VTA mice | — | .688 / .429 / .152 |
| b (VTA, top 30 % pupil) | growth of the almorexant – vehicle difference, +1.5 vs -1 z VTA oxLight | Bootstrap over mice | 7 VTA mice, 7,220 transitions | bootstrap | .243 |
| b (SNc, bottom 30 % pupil) | growth of the drug difference along SNc oxLight | Bootstrap over mice | 8 SNc mice, 8,423 transitions | bootstrap | .269 |
| b (SNc, top 30 % pupil) | growth of the drug difference along SNc oxLight | Bootstrap over mice | 8 SNc mice, 7,150 transitions | bootstrap | .973 |
| c (SNc) | almorexant vs vehicle | Paired t-test | 8 SNc mice | t(7) = 0.54 | .61 |
| c (VTA) | almorexant vs vehicle | Paired t-test | 7 VTA mice | t(6) = 1.44 | .20 |
| d (SNc) | almorexant vs vehicle | Paired t-test | 8 SNc mice | t(7) = 2.11 | .073 |
| d (VTA) | almorexant vs vehicle | Paired t-test | 7 VTA mice | t(6) = 0.47 | .65 |
| e (disengage, SNc) | almorexant vs vehicle | Paired t-test | 8 SNc mice | — | .755 |
| e (disengage, VTA) | almorexant vs vehicle | Paired t-test | 8 VTA mice | — | .973 |
| e (re-engage, SNc) | almorexant vs vehicle | Paired t-test | 7 SNc | — | .34 |
| e (re-engage, VTA) | almorexant vs vehicle | Paired t-test | 7 VTA mice | — | .329 |
| f (low reward) | almorexant vs vehicle, anticipatory | Paired t-test | 13 mice | t(12) = 1.24 | .240 |
| f (low reward) | almorexant vs vehicle, slow | Paired t-test | 13 mice | t(12) = 2.35 | .037 |
| f (low reward) | almorexant vs vehicle, lapse | Paired t-test | 13 mice | t(12) = 3.75 | .003 |
| f (high reward) | almorexant vs vehicle, anticipatory | Paired t-test | 13 mice | t(12) = -0.88 | .397 |
| f (high reward) | almorexant vs vehicle, slow | Paired t-test | 13 mice | t(12) = 1.86 | .088 |
| f (high reward) | almorexant vs vehicle, lapse | Paired t-test | 13 mice | t(12) = 0.76 | .463 |
| g (low reward) | almorexant vs vehicle, engaged trials | Paired t-test | 13 mice | t(12) = 4.82 | < .001 |
| g (low reward) | almorexant vs vehicle, anticipatory trials | Paired t-test | 3 mice | t(2) = 1.46 | .282 |
| g (low reward) | almorexant vs vehicle, slow trials | Paired t-test | 13 mice | t(12) = 3.41 | .005 |
| g (low reward) | almorexant vs vehicle, lapse trials | Paired t-test | 11 mice | t(10) = -0.75 | .473 |
| g (high reward) | almorexant vs vehicle, engaged trials | Paired t-test | 13 mice | t(12) = 6.28 | < .001 |
| g (high reward) | almorexant vs vehicle, anticipatory trials | Paired t-test | < 3 mice | not tested (n < 3) |  |
| g (high reward) | almorexant vs vehicle, slow trials | Paired t-test | 13 mice | t(12) = 3.88 | .002 |
| g (high reward) | almorexant vs vehicle, lapse trials | Paired t-test | 4 mice | t(3) = -1.55 | .219 |
| h (slow) | almorexant vs vehicle, session third 1st | Paired t-test | 13 mice | t(12) = 2.39 | .034 |
| h (slow) | almorexant vs vehicle, session third 2nd | Paired t-test | 13 mice | t(12) = 2.18 | .050 |
| h (slow) | almorexant vs vehicle, session third 3rd | Paired t-test | 13 mice | t(12) = 2.02 | .067 |
| h (lapse) | almorexant vs vehicle, session third 1st | Paired t-test | 13 mice | t(12) = 1.22 | .247 |
| h (lapse) | almorexant vs vehicle, session third 2nd | Paired t-test | 13 mice | t(12) = 3.53 | .004 |
| h (lapse) | almorexant vs vehicle, session third 3rd | Paired t-test | 13 mice | t(12) = 1.91 | .080 |
| i (NAcc) | almorexant vs vehicle, high reward | Paired t-test | 8 mice | t(7) = -0.95 | .372 |
| i (NAcc) | almorexant vs vehicle, low reward | Paired t-test | 8 mice | t(7) = -0.20 | .846 |
| i (DLS) | almorexant vs vehicle, high reward | Paired t-test | 8 mice | t(7) = -0.88 | .407 |
| i (DLS) | almorexant vs vehicle, low reward | Paired t-test | 8 mice | t(7) = -0.07 | .947 |

##### Extended Data Fig. 7

|  |  |  |  |  |  |
| --- | --- | --- | --- | --- | --- |
| a (5 Hz) | low reward: orexin stim – control vs 0 | One-sample t-test | 8 orexin mice | t(7) = -9.74 | < .001 |
| a (5 Hz) | low reward: orexin vs heat | Welch t-test | 8 orexin, 7 heat mice | t(8.0) = -5.43 | < .001 |
| a (5 Hz) | high reward: orexin stim – control vs 0 | One-sample t-test | 8 orexin mice | t(7) = -5.67 | < .001 |
| a (5 Hz) | high reward: orexin vs heat | Welch t-test | 8 orexin, 7 heat mice | t(7.9) = -2.88 | .021 |
| a (5 Hz) | bracket: low vs high within orexin | Paired t-test | 8 orexin mice | t(7) = -5.27 | .001 |
| a (20 Hz) | low reward: orexin stim – control vs 0 | One-sample t-test | 8 orexin mice | t(7) = -3.79 | .007 |
| a (20 Hz) | low reward: orexin vs heat | Welch t-test | 8 orexin, 7 heat mice | t(10.5) = -4.25 | .002 |
| a (20 Hz) | high reward: orexin stim – control vs 0 | One-sample t-test | 8 orexin mice | t(7) = -5.21 | .001 |
| a (20 Hz) | high reward: orexin vs heat | Welch t-test | 8 orexin, 7 heat mice | t(10.1) = -3.35 | .007 |
| a (20 Hz) | bracket: low vs high within orexin | Paired t-test | 8 orexin mice | t(7) = -2.55 | .038 |
| b (disengage) | 5 Hz, low reward: laser vs sham | Paired t-test | 8 orexin mice | t(7) = -0.62 | .554 |
| b (disengage) | 20 Hz, low reward: laser vs sham | Paired t-test | 7 orexin mice | t(6) = 0.20 | .850 |
| b (disengage) | 5 Hz, high reward: laser vs sham | Paired t-test | 8 orexin mice | t(7) = -2.16 | .068 |
| b (disengage) | 20 Hz, high reward: laser vs sham | Paired t-test | 8 orexin mice | t(7) = 0.95 | .372 |
| b (re-engage) | 5 Hz, low reward: laser vs sham | Paired t-test | 7 orexin mice | t(6) = -1.36 | .222 |
| b (re-engage) | 20 Hz, low reward: laser vs sham | Paired t-test | 7 orexin mice | t(6) = 0.31 | .764 |
| b (re-engage) | 5 Hz, high reward: laser vs sham | Paired t-test | 4 orexin mice | t(3) = 0.14 | .898 |
| b (re-engage) | 20 Hz, high reward: laser vs sham | Paired t-test | 4 orexin mice | t(3) = -0.05 | .962 |
| c (5 Hz, laser – sham) | bracket: low vs high reward block | Paired t-test | 8 orexin mice | t(7) = 2.57 | .037 |
| c (20 Hz, laser – sham) | bracket: low vs high reward block | Paired t-test | 8 orexin mice | t(7) = 3.24 | .014 |
| d (middle, continuous, 5 Hz) | low vs high reward block | Paired t-test | 8 orexin mice | t(7) = 2.06 | .078 |
| d (middle, continuous, 20 Hz) | low vs high reward block | Paired t-test | 8 orexin mice | t(7) = 3.50 | .010 |
| d (right, control) | low vs high reward block, control trials on laser days | Paired t-test | 8 orexin mice | t(7) = -1.35 | .220 |
| d (right, 20 Hz) | low vs high reward block, 20-Hz stimulated trials | Paired t-test | 8 orexin mice | t(7) = 3.44 | .011 |
| e (low reward) | 5 Hz, arousal route via the disengage hazard, laser vs sham | Paired t-test; bootstrap over mice | 8 orexin mice | t(7) = -0.06 | .957 |
| e (low reward) | 5 Hz, arousal route via the re-engage hazard, laser vs sham | Paired t-test; bootstrap over mice | 8 orexin mice | t(7) = -1.10 | .307 |
| e (low reward) | 20 Hz, arousal route via the disengage hazard, laser vs sham | Paired t-test; bootstrap over mice | 8 orexin mice | t(7) = 3.18 | .016 |
| e (low reward) | 20 Hz, arousal route via the re-engage hazard, laser vs sham | Paired t-test; bootstrap over mice | 8 orexin mice | t(7) = 1.09 | .310 |
| e (high reward) | 5 Hz, arousal route via the disengage hazard, laser vs sham | Paired t-test; bootstrap over mice | 7 orexin mice | t(6) = 0.22 | .835 |
| e (high reward) | 5 Hz, arousal route via the re-engage hazard, laser vs sham | Paired t-test; bootstrap over mice | 7 orexin mice | t(6) = 1.35 | .226 |
| e (high reward) | 20 Hz, arousal route via the disengage hazard, laser vs sham | Paired t-test; bootstrap over mice | 7 orexin mice | t(6) = 2.47 | .048 |
| e (high reward) | 20 Hz, arousal route via the re-engage hazard, laser vs sham | Paired t-test; bootstrap over mice | 7 orexin mice | t(6) = 2.73 | .034 |
| f (left, P(engaged)) | post-stimulation trials, laser – sham at matched pupil, low reward: all matched trials | One-sample t-test | 8 orexin mice | t(7) = 2.63 | .034 |
| f (left, P(engaged)) | post-stimulation trials, laser – sham at matched pupil, low reward: low pupil third | One-sample t-test | 8 orexin mice | t(7) = 2.52 | .040 |
| f (left, P(engaged)) | post-stimulation trials, laser – sham at matched pupil, low reward: mid pupil third | One-sample t-test | 8 orexin mice | t(7) = 2.16 | .068 |
| f (left, P(engaged)) | post-stimulation trials, laser – sham at matched pupil, low reward: high pupil third | One-sample t-test | 8 orexin mice | t(7) = 2.61 | .035 |
| f (middle, P(engaged \| prev. eng.)) | post-stimulation trials, laser – sham at matched pupil, low reward: all matched trials | One-sample t-test | 8 orexin mice | t(7) = 2.44 | .045 |
| f (middle, P(engaged \| prev. eng.)) | post-stimulation trials, laser – sham at matched pupil, low reward: low pupil third | One-sample t-test | 8 orexin mice | t(7) = 1.28 | .24 |
| f (middle, P(engaged \| prev. eng.)) | post-stimulation trials, laser – sham at matched pupil, low reward: mid pupil third | One-sample t-test | 8 orexin mice | t(7) = 2.27 | .057 |
| f (middle, P(engaged \| prev. eng.)) | post-stimulation trials, laser – sham at matched pupil, low reward: high pupil third | One-sample t-test | 8 orexin mice | t(7) = 2.48 | .042 |
| f (right, P(disengaged next)) | post-stimulation trials, laser – sham at matched pupil, low reward: all / low / mid / high third | One-sample t-test | 8 orexin mice | t(7) = -1.92 / -1.01 / -2.32 / -1.33 | .097 / .346 / .054 |
| g (5 Hz) | low reward: vGlut stim – control vs 0 | One-sample t-test | 5 vGlut mice | t(4) = -3.07 | .037 |
| g (5 Hz) | low reward: vGlut vs heat | Welch t-test | 5 vGlut, 7 heat | t(5.8) = -3.57 | .013 |
| g (5 Hz) | high reward: vGlut stim – control vs 0 | One-sample t-test | 5 vGlut mice | t(4) = -2.34 | .080 |
| g (5 Hz) | high reward: vGlut vs heat | Welch t-test | 5 vGlut, 7 heat | t(4.4) = -2.47 | .064 |
| g (5 Hz) | bracket: low vs high within vGlut | Paired t-test | 5 vGlut mice | t(4) = 0.51 | .634 |
| g (20 Hz) | low reward: vGlut stim – control vs 0 | One-sample t-test | 5 vGlut mice | t(4) = -9.97 | < .001 |
| g (20 Hz) | low reward: vGlut vs heat | Welch t-test | 5 vGlut, 7 heat | t(5.2) = -9.99 | < .001 |
| g (20 Hz) | high reward: vGlut stim – control vs 0 | One-sample t-test | 5 vGlut mice | t(4) = -29.43 | < .001 |
| g (20 Hz) | high reward: vGlut vs heat | Welch t-test | 5 vGlut, 7 heat | t(9.5) = -21.51 | < .001 |
| g (20 Hz) | bracket: low vs high within vGlut | Paired t-test | 5 vGlut mice | t(4) = 2.18 | .095 |
| h (right) | vGlut 5 Hz stim – control vs 0 | One-sample t-test | 5 vGlut mice | t(4) = 4.47 | .011 |
| h (right) | vGlut 20 Hz stim – control vs 0 | One-sample t-test | 5 vGlut mice | t(4) = 8.64 | < .001 |

##### Extended Data Fig. 8

|  |  |  |  |  |  |
| --- | --- | --- | --- | --- | --- |
| a (inset, vehicle) | switch into low: trials 2-3 minus trial 1, vehicle | Paired t-test | 8 mice | — | .007 |
| a (inset, semaglutide) | switch into low: trials 2-3 minus trial 1, semaglutide | Paired t-test | 8 mice | — | .42 |
| b (left) | reward AUC change after a switch into high, trials 2-10, semaglutide vs vehicle | Paired t-test | 8 mice | t(7) = -3.81 | .007 |
| b (right) | reward AUC change after a switch into low, trials 2-10, semaglutide vs vehicle | Paired t-test | 8 mice | t(7) = 5.37 | .001 |
| c | within-block slope, low reward blocks, semaglutide vs vehicle | Paired t-test | 8 mice | t(7) = -4.71 | .002 |
| c | within-block slope, high reward blocks, semaglutide vs vehicle | Paired t-test | 8 mice | t(7) = -3.79 | .007 |
| d | inside-block reward kernel, summed weight of lags 2-5, semaglutide vs vehicle | Paired t-test | 8 mice | t(7) = -6.00 | < .001 |
| e (right) | hard – easy reward weight, predicted (confidence-weighted PE integrator) vs observed, vehicle, semaglutide experiment | Paired t-test | 8 mice | t(7) = -3.60 | .009 |
| f (left) | inside-block reward kernel, lags 2-5, almorexant vs vehicle | Paired t-test | 8 mice | t(7) = -0.33 | .75 |
| f (right) | integrator response at 1st switch into high, almorexant vs vehicle (trials 1-30) | Paired t-test | 8 mice | t(7) = -0.22 | .83 |
| h | surprise: 1st reward – mean(2nd, 3rd), both directions, semaglutide vs vehicle | Paired t-test | 8 mice | — | .24 |
| h | switch step: 1st reward – pre, both directions, semaglutide vs vehicle | Paired t-test | 8 mice | — | .29 |
| h | steady NAcc reward response, low reward: semaglutide vs vehicle | Paired t-test | 8 mice | t(7) = –0.83 | .43 |
| h | steady NAcc reward response, high reward: semaglutide vs vehicle | Paired t-test | 8 mice | t(7) = –2.65 | .033 |

##### Extended Data Fig. 9

|  |  |  |  |  |  |
| --- | --- | --- | --- | --- | --- |
| a | constant-reward task, NAcc reward response, semaglutide vs vehicle | Paired t-test | 8 mice | t(7) = 0.83 | .43 |
| a | constant-reward task, DLS reward response, semaglutide vs vehicle | Paired t-test | 8 mice | t(7) = 1.02 | .34 |
| b | NAcc high – low reward response, blocks 1-2, semaglutide vs vehicle | Paired t-test | 8 mice (45 / 50 sessions) | t(7) = –3.97 | .005 |
| b | NAcc high – low reward response, blocks 3 onwards, semaglutide vs vehicle | Paired t-test | 8 mice (45 / 50 sessions) | t(7) = –3.44 | .011 |
| c | tonic utility code, semaglutide – vehicle | Paired t-test | 8 mice | t(7) = –3.90 | .006 |
| c | phasic utility code, semaglutide – vehicle | Paired t-test | 8 mice | t(7) = –3.97 | .005 |
| d | early phasic utility code (blocks 1-2) vs P(disengaged) blocks >= 3, vehicle sessions | Fisher-z r, one-sample t-test | 8 mice (50 vehicle / 45 semaglutide sessions) | r = +0.06 | .83 |
| d | early phasic utility code (blocks 1-2) vs P(disengaged) blocks >= 3, semaglutide sessions | Fisher-z r, one-sample t-test | 8 mice (50 vehicle / 45 semaglutide sessions) | r = –0.56 | .024 |
| d | early phasic utility code (blocks 1-2) vs P(disengaged) blocks >= 3, both drugs, within mouse | Fisher-z r, one-sample t-test | 8 mice (50 vehicle / 45 semaglutide sessions) | r = –0.55; t(7) = –4.47 | .003 |
| e (low) | NAcc reward response 0.1-0.6 s, low-reward blocks, matched pairs, semaglutide vs vehicle | Paired t-test | 8 mice (14 pairs) | t(7) = –0.34 | .74 |
| e (high) | NAcc reward response 0.1-0.6 s, high-reward blocks, matched pairs, semaglutide vs vehicle | Paired t-test | 8 mice (14 pairs) | t(7) = –3.28 | .013 |
| f | high – low reward-aligned NAcc response, 0.1-0.6 s, matched pairs, semaglutide vs vehicle | Paired t-test | 8 mice (14 pairs) | t(7) = –4.08 | .005 |
| g | tonic code (high – low pre-stimulus NAcc level), brightness-matched session pairs, semaglutide vs vehicle | Paired t-test | 8 mice (14 matched pairs, caliper 10 %) | t(7) = –2.97 | .021 |
| h (VTA) | mouse-level r, early oxLight code vs late P(disengaged), almorexant, against 0 | One-sample t-test | 7 mice | t(6) = 0.76 | .48 |
| h (VTA) | mouse-level r, early oxLight code vs late P(disengaged), vehicle (almorexant experiment), against 0 | One-sample t-test | 7 mice | t(6) = –0.80 | .45 |
| h (VTA) | mouse-level r, early oxLight code vs late P(disengaged), semaglutide, against 0 | One-sample t-test | 7 mice | t(6) = 1.53 | .18 |
| h (VTA) | mouse-level r, early oxLight code vs late P(disengaged), vehicle (semaglutide experiment), against 0 | One-sample t-test | 8 mice | t(7) = 0.81 | .45 |
| h (SNc) | mouse-level r, early oxLight code vs late P(disengaged), almorexant, against 0 | One-sample t-test | 8 mice | t(7) = 0.73 | .49 |
| h (SNc) | mouse-level r, early oxLight code vs late P(disengaged), vehicle (almorexant experiment), against 0 | One-sample t-test | 8 mice | t(7) = 1.74 | .13 |
| h (SNc) | mouse-level r, early oxLight code vs late P(disengaged), semaglutide, against 0 | One-sample t-test | 6 mice | t(5) = 0.85 | .43 |
| h (SNc) | mouse-level r, early oxLight code vs late P(disengaged), vehicle (semaglutide experiment), against 0 | One-sample t-test | 7 mice | t(6) = –0.71 | .50 |

##### Extended Data Fig. 10

|  |  |  |  |  |  |
| --- | --- | --- | --- | --- | --- |
| a | DLS cue value (high – low cue response), semaglutide vs vehicle | Paired t-test | 8 mice | t(7) = –4.16 | .004 |
| a | DLS confidence slope (easy – hard cue response), semaglutide vs vehicle | Paired t-test | 8 mice | t(7) = –2.10 | .073 |
| b | DLS reward AUC –0.5 to 1.0 s, low-reward blocks, semaglutide vs vehicle | Paired t-test | 8 mice | t(7) = 0.14 | .89 |
| b | DLS reward AUC –0.5 to 1.0 s, high-reward blocks, semaglutide vs vehicle | Paired t-test | 8 mice | t(7) = –3.59 | .009 |
| e (left) | reward-size prediction error at a switch, semaglutide vs vehicle | Paired t-test | 8 mice | — | .018 |
| e (middle) | switch step: 1st reward – pre, both directions, semaglutide vs vehicle | Paired t-test | 8 mice | t(7) = –2.90 | .023 |
| e (right) | steady DLS reward response, low reward: semaglutide vs vehicle | Paired t-test | 8 mice | t(7) = –0.35 | .74 |
| e (right) | steady DLS reward response, high reward: semaglutide vs vehicle | Paired t-test | 8 mice | t(7) = –4.04 | .0049 |
| f | integrator step at the 1st switch into high vs session trend of the phasic DLS reward response, vehicle sessions of both experiments | Permutation | 111 sessions, 9 mice | r = 0.23 | .022 |
| f | integrator step at the 1st switch into high vs session trend of the within-block SD of baseline pupil, vehicle sessions of both experiments | Permutation | 113 sessions, 9 mice | r = –0.32 | < .001 |
| g | NAcc reward step (model D), semaglutide vs vehicle | Paired t-test; bootstrap over mice | 8 mice | t(7) = –3.54 | .009 |
| g | NAcc utility code (tonic), semaglutide vs vehicle | Paired t-test; bootstrap over mice | 8 mice | t(7) = –3.90 | .006 |
| g | NAcc size code (phasic), semaglutide vs vehicle | Paired t-test; bootstrap over mice | 8 mice | t(7) = –2.81 | .026 |
| g | NAcc cue value, semaglutide vs vehicle | Paired t-test; bootstrap over mice | 8 mice | t(7) = –2.45 | .044 |
| g | NAcc switch RPE (step), semaglutide vs vehicle | Paired t-test; bootstrap over mice | 8 mice | t(7) = –1.14 | .29 |
| g | NAcc cue confidence, semaglutide vs vehicle | Paired t-test; bootstrap over mice | 8 mice | t(7) = –1.26 | .25 |
| g | NAcc reward response, semaglutide vs vehicle | Paired t-test; bootstrap over mice | 8 mice | t(7) = –1.68 | .14 |
| g | DLS cue value, semaglutide vs vehicle | Paired t-test; bootstrap over mice | 8 mice | t(7) = –4.16 | .004 |
| g | DLS switch RPE (step), semaglutide vs vehicle | Paired t-test; bootstrap over mice | 8 mice | t(7) = –2.90 | .023 |
| g | DLS cue confidence, semaglutide vs vehicle | Paired t-test; bootstrap over mice | 8 mice | t(7) = –2.10 | .073 |
| g | DLS reward response, semaglutide vs vehicle | Paired t-test; bootstrap over mice | 8 mice | $t(7) = -0.97$ | .37 |
